# Diffusion MRI Indices and their Relation to Cognitive Impairment in Brain Aging: The updated multi-protocol approach in ADNI3

**DOI:** 10.1101/476721

**Authors:** Artemis Zavaliangos-Petropulu, Talia M. Nir, Sophia I. Thomopoulos, Robert I. Reid, Matt A. Bernstein, Bret Borowski, Clifford R. Jack, Michael W. Weiner, Neda Jahanshad, Paul M. Thompson

**Affiliations:** Imaging Genetics Center, Mark & Mary Stevens Neuroimaging & Informatics Institute, Keck School of Medicine, University of Southern California, Marina del Rey, CA, USA; Department of Information Technology, Mayo Clinic and Foundation, Rochester, MN, USA; Department of Radiology, Mayo Clinic and Foundation, Rochester, MN, USA; Department of Radiology, University of California San Francisco School of Medicine, San Francisco, CA, USA

**Keywords:** Alzheimer’s disease, ADNI3, White Matter, DTI, Multi-site, Harmonization, TDF, ComBat

## Abstract

Brain imaging with diffusion-weighted MRI (dMRI) is sensitive to microstructural white matter changes associated with brain aging and neurodegeneration. In its third phase, the Alzheimer’s Disease Neuroimaging Initiative (ADNI3) is collecting data across multiple sites and scanners using different dMRI acquisition protocols, to better understand disease effects. It is vital to understand when data can be pooled across scanners, and how the choice of dMRI protocol affects the sensitivity of extracted measures to differences in clinical impairment. Here, we analyzed ADNI3 data from 317 participants (mean age: 75.4±7.9 years; 143 men/174 women), who were each scanned at one of 47 sites with one of six dMRI protocols using scanners from three different manufacturers. We computed four standard diffusion tensor imaging (DTI) indices including fractional anisotropy (FA^DTI^) and mean, radial, and axial diffusivity, and one FA index based on the tensor distribution function (FA^TDF^), in 24 bilaterally averaged white matter regions of interest. We found that protocol differences significantly affected dMRI indices, in particular FA^DTI^. We ranked the diffusion indices for their strength of association with four clinical assessments. In addition to diagnosis, we evaluated cognitive impairment as indexed by three commonly used screening tools for detecting dementia and Alzheimer’s disease: the Alzheimer’s Disease Assessment Scale (ADAS-cog), the Mini-Mental State Examination (MMSE), and the Clinical Dementia Rating scale sum-of-boxes (CDR-sob). Using a nested random-effects model to account for protocol and site, we found that across all dMRI indices and clinical measures, the hippocampal-cingulum and fornix (*crus*) / *stria terminalis* regions most consistently showed strong associations with clinical impairment. Overall, the greatest effect sizes were detected in the hippocampal-cingulum and uncinate fasciculus for associations between axial or mean diffusivity and CDR-sob. FA^TDF^ detected robust widespread associations with clinical measures, while FA^DTI^ was the weakest of the five indices for detecting associations. Ultimately, we were able to successfully pool dMRI data from multiple acquisition protocols from ADNI3 and detect consistent and robust associations with clinical impairment and age.

## 1 Introduction

Alzheimer’s disease (AD) is the most common type of dementia, affecting approximately 10% of the population over age 65 (Alzheimer’s Association, 2018). As life expectancy increases, there is an ever-increasing need for sensitive biomarkers of AD - to better understand the disease, and to serve as surrogate markers of disease burden for use in treatment and prevention trials. The Alzheimer’s Disease Neuroimaging Initiative (ADNI) is an ongoing large-scale, multi-center, longitudinal study designed to improve methods for clinical trials by identifying brain imaging, clinical, cognitive, and molecular biomarkers of AD and aging. Now in its third phase (ADNI3), ADNI continues to incorporate newer technologies as they become established (Jack et al., 2015); data from ADNI, collected at participating sites across the U.S. and Canada, is publicly available and has been used in a diverse range of publications (Veitch et al., 2018).

ADNI’s second phase (ADNI2) introduced, to the initiative, the use of diffusion-weighted MRI (dMRI) as an additional approach for tracking AD progression (Jack et al., 2015). dMRI has since been used in numerous studies to understand the effects of AD on white matter (WM) microstructure and brain connectivity (Daianu et al., 2013a,b; Nir et al., 2013; Prasad et al., 2013). Some of these approaches assess dMRI indices in normal appearing WM (Giulietti et al., 2018), while others use tractography and graph-theoretic analysis to study abnormalities in structural brain networks (Nir et al., 2015; Hu et al., 2016; Maggipinto et al., 2017; Sulaimany et al., 2017; Powell et al., 2018; Sanchez-Rodriguez et al., 2018). In aggregate, these studies point to WM abnormalities in AD, which may play a key role in early pathogenesis and diagnosis (Sachdev et al., 2013).

ADNI2 acquired dMRI data with one acquisition protocol from approximately one third of enrolled participants at the subset of ADNI sites that used 3 tesla General Electric (GE) scanners. To ensure that dMRI could be collected from all enrolled participants, ADNI3 developed new dMRI protocols for all GE, Siemens and Philips scanners used across ADNI sites. Now, data is being acquired with seven different dMRI acquisition protocols (see methods for details; http://adni.loni.usc.edu/methods/documents/mri-protocols/). ADNI3 began in October 2016, and has already acquired data from over 300 participants. dMRI spatial resolution was improved between ADNI2 and ADNI3 by reducing the voxel size from 2.7×2.7×2.7 mm to 2.0×2.0×2.0 mm. While voxel size (i.e., spatial resolution) remains consistent across all seven ADNI3 protocols, angular resolution (the number of gradient directions) varies across protocols to accommodate scanner restrictions and to ensure that the multi-modal scanning session is completed in under 60 minutes. Although many large-scale multi-site DTI studies have obtained consistent results even when acquisition protocols across sites are not harmonized in advance (Jahanshad et al., 2013; Kochunov et al., 2014; Acheson et al., 2017; Kelly et al., 2018), differences in dMRI acquisition parameters, including vendor, voxel size, and angular resolution, are known to affect derived dMRI measures (Alexander et al., 2001; Cercignani et al., 2003; Zhan et al., 2010; Zhu et al., 2011). As a result, improved harmonization of multi-site diffusion data is of great interest (Grech-Sollars et al., 2015; Pohl et al., 2016; Palacios et al., 2017). For example, ComBat - originally developed to model and remove batch effects from genomic microarray data (Johnson et al., 2007) - was one of the most effective methods for harmonizing DTI measures in a recent comparison of such techniques (Fortin et al., 2017).

Here we tested whether standard diffusion tensor imaging (DTI)-derived anisotropy and diffusivity indices, calculated from multiple imaging protocols in ADNI3, can be pooled and harmonized to show robust associations with age and four clinical assessments. In addition to diagnosis, cognitive impairment was assessed with three commonly used screening tools for detecting dementia and Alzheimer’s disease: the Alzheimer’s Disease Assessment Scale (ADAS-cog; Rosen et al., 1984), the Mini-Mental State Examination (MMSE; Folstein et al., 1975), and the Clinical Dementia Rating scale sum-of-boxes (CDR-sob; Berg, 1988). For the rest of the paper we refer to these tools as “cognitive measures”. In addition to standard DTI indices - the fractional anisotropy (FA^DTI^), mean diffusivity (MD^DTI^), radial diffusivity (RD^DTI^), and axial diffusivity (AxD^DTI^) - we also evaluated a modified measure of FA, derived from the tensor distribution function (FA^TDF^; Leow et al., 2009) which can be more sensitive to neurodegenerative disease-related WM abnormalities than FA^DTI^ across high- and low-angular resolution dMRI (Nir et al., 2017). The TDF model addresses well-established limitations of the standard single-tensor diffusion model - which cannot resolve complex profiles of WM architecture such as crossing or mixing fibers, present in up to 90% of WM voxels (Tournier et al., 2004; Descoteaux et al., 2007, 2009; Jeurissen et al., 2013).

In 24 WM regions of interest (ROIs), we ranked these five anisotropy and diffusivity indices, in terms of their strength of association with key clinical measures, to identify dMRI indices that may help understand and track AD progression. We hypothesized that the diffusion indices from ADNI2 (Nir et al., 2013, 2017) would still be associated with clinical measures of disease burden in ADNI3 - despite the variation in protocols. We hypothesized that when data were pooled across ADNI3 protocols: (1) higher diffusivity and lower anisotropy in the temporal lobe white matter would be most sensitive to cognitive impairment, with highest effect sizes for associations with CDR-sob, and (2) FA^TDF^ would detect associations with clinical impairment with higher effect sizes than FA^DTI^.

## 2 Methods

### 2.1 ADNI participants

Baseline MRI, DTI, diagnosis, demographics, and cognitive measures were downloaded from the ADNI database (https://ida.loni.usc.edu/). This analysis was performed when data collection for ADNI3 was still ongoing (May 2018), and reflects the data available on April 30, 2018. Of the 381 participants scanned to date, 55 were excluded after quality assurance: this included ensuring complete clinical and demographic information, and image-level quality control (removing scans with severe motion, missing volumes, or corrupt files). To ensure sufficient statistical power to assess differences in data collected with different protocols, we evaluated only those protocols with complete available data for at least 10 participants at the time of download; we did not assess protocol GE36, for which scans from 9 of 12 participants passed quality assurance. Details on excluded participants are outlined in **Supplementary Table 1**.

317 remaining participants - from 47 scanning sites - were included in the analysis (mean age: 75.4±7.9 yrs; 143 men, 174 women; **Table 1**): 211 were elderly cognitively normal controls (CN; mean age: 74.5±7.3 yrs; 84 men, 127 women), 84 were diagnosed with mild cognitive impairment (MCI); mean age: 76.3±8.1 yrs; 48 men, 36 women), and 22 were diagnosed with Alzheimer’s disease (AD; mean age: 80.6±10.5 yrs; 11 men, 11 women). We note that two of the ADNI2 diagnostic categories - CN and Significant Memory Concern (SMC) - were combined and identified as CN in ADNI3. ADNI2’s early and late MCI categories were combined and identified as MCI in ADNI3.

**Table 1.**
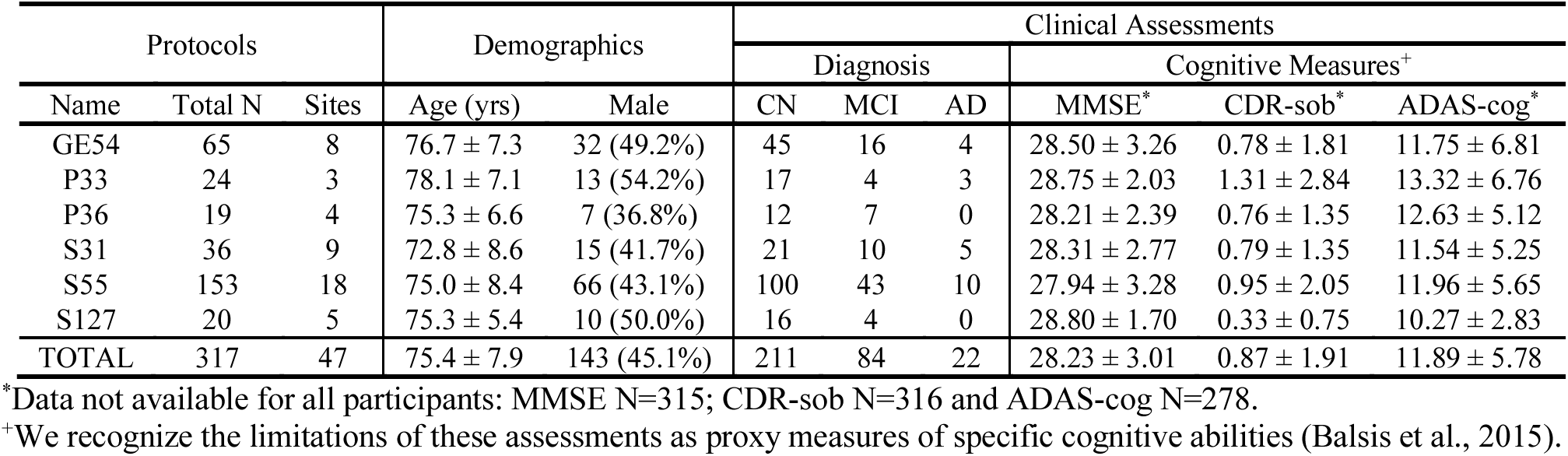
Demographic and clinical measures for participants in ADNI3, subdivided by dMRI protocol. We report the average age, MMSE, CDR-sob, and ADAS-cog measures, and their standard deviations.

| Protocols |  |  | Demographics |  | Clinical Assessments |  |  |  |  |  |
| --- | --- | --- | --- | --- | --- | --- | --- | --- | --- | --- |
|  |  |  |  |  | Diagnosis |  |  | Cognitive Measures<sup>†</sup> |  |  |
| Name | Total N | Sites | Age (yrs) | Male | CN | MCI | AD | MMSE<sup>*</sup> | CDR-sob<sup>*</sup> | ADAS-cog<sup>*</sup> |
| GE54 | 65 | 8 | 76.7 ± 7.3 | 32 (49.2%) | 45 | 16 | 4 | 28.50 ± 3.26 | 0.78 ± 1.81 | 11.75 ± 6.81 |
| P33 | 24 | 3 | 78.1 ± 7.1 | 13 (54.2%) | 17 | 4 | 3 | 28.75 ± 2.03 | 1.31 ± 2.84 | 13.32 ± 6.76 |
| P36 | 19 | 4 | 75.3 ± 6.6 | 7 (36.8%) | 12 | 7 | 0 | 28.21 ± 2.39 | 0.76 ± 1.35 | 12.63 ± 5.12 |
| S31 | 36 | 9 | 72.8 ± 8.6 | 15 (41.7%) | 21 | 10 | 5 | 28.31 ± 2.77 | 0.79 ± 1.35 | 11.54 ± 5.25 |
| S55 | 153 | 18 | 75.0 ± 8.4 | 66 (43.1%) | 100 | 43 | 10 | 27.94 ± 3.28 | 0.95 ± 2.05 | 11.96 ± 5.65 |
| S127 | 20 | 5 | 75.3 ± 5.4 | 10 (50.0%) | 16 | 4 | 0 | 28.80 ± 1.70 | 0.33 ± 0.75 | 10.27 ± 2.83 |
| TOTAL | 317 | 47 | 75.4 ± 7.9 | 143 (45.1%) | 211 | 84 | 22 | 28.23 ± 3.01 | 0.87 ± 1.91 | 11.89 ± 5.78 |

<sup>\*</sup>Data not available for all participants: MMSE N=315; CDR-sob N=316 and ADAS-cog N=278.
<sup>†</sup>We recognize the limitations of these assessments as proxy measures of specific cognitive abilities (Balsis et al., 2015).

### 2.2 Clinical assessments

In addition to diagnosis, we indexed cognitive impairment using total scores from commonly used screening tools for detecting dementia and AD (**Table 1**): the Alzheimer’s Disease Assessment Scale 13 (ADAS-cog), the Mini-Mental State Examination (MMSE), and the Clinical Dementia Rating scale sum-of-boxes (CDR-sob). We refer to these tools as “cognitive measures”, but recognize the limitations of these assessments as proxy measures of specific cognitive abilities (Balsis et al., 2015). The ADAS-cog is frequently used in pharmaceutical trials, with scores ranging from 0-70; higher scores represent more severe cognitive dysfunction (Rosen et al., 1984). MMSE is more often used by clinicians and researchers in assessing cognitive aging. Scores for MMSE range from 0-30; lower scores typically indicate greater cognitive dysfunction (Folstein et al., 1975). CDR-sob is used primarily in clinical trials and in clinical practice for evaluating disease severity including the mild and early symptomatic stages of dementia. It is calculated based on the sum of severity ratings in six domains (‘boxes’) - memory, orientation, judgment and problem solving, community affairs, home and hobbies, and personal care. Scores range from 0 (no dementia) to 3 (severe dementia; Rosen et al., 1984). These evaluations are among the measures used in diagnosing ADNI participants. Not all cognitive measures were available for every participant (MMSE, N=315; CDR-sob, N=316, and ADAS-cog, N=278; **Supplementary Table 2** lists these by protocol).

### 2.3 Diffusion MRI acquisition protocols

ADNI3 incorporated dMRI protocols for 3 tesla Siemens, Philips, and GE scanners. ADNI2, the first phase of ADNI to include diffusion MRI, only prescribed dMRI protocols for GE scanners. The available scanners span a wide range of software capabilities, such as support (or the lack of it) for custom diffusion gradient tables and/or simultaneous multi-slice acceleration. Including additional scanners while staying in a 7-10 minute scan duration resulted in data acquired with seven different acquisition protocols - of which six had sufficient sample sizes to be evaluated here. Protocols varied in the number of diffusion weighted imaging (DWI) directions (i.e., angular resolution), and the number of non-diffusion sensitized gradients (*b*_0_ images), which serve as a reference to assess diffusion-related decay of the MR signal. Voxel size across all ADNI3 protocols was 2.0×2.0×2.0 mm^3^ and 2.7×2.7×2.7 mm^3^ in ADNI2. **Table 2** summarizes the different protocols.

**Table 2.** ADNI diffusion MRI acquisition protocols

| | Name | Scanner | Protocol | $b_0$<br>Volumes | DWI<br>Volumes | Total<br>Volumes | Time<br>(min) | Total<br>N |
| --- | --- | --- | --- | --- | --- | --- | --- | --- |
| ADNI3 | GE36 | GE | Basic Widebore 25x | 4 $b=0$ s/mm<sup>2</sup> | 32 $b=1000$ s/mm<sup>2</sup> | 36 | 9:52 | -- |
| | GE54 | GE | Basic 25x | 6 $b=0$ s/mm<sup>2</sup> | 48 $b=1000$ s/mm<sup>2</sup> | 54 | 7:09 | 65 |
| | P33 | Philips | Basic Widebore | 1 $b=0$ s/mm<sup>2</sup> | 32 $b=1000$ s/mm<sup>2</sup> | 33 | 7:32 | 24 |
| | P36 | Philips | Basic Widebore R3 | 1 $b=0$ s/mm<sup>2</sup><br>3 $b=2$ s/mm<sup>2</sup> | 32 $b=1000$ s/mm<sup>2</sup> | 36 | 6:54 | 19 |
| | P54 | Philips | Basic R5 | 1 $b=0$ s/mm<sup>2</sup><br>5 $b=2$ s/mm<sup>2</sup> | 48 $b=1000$ s/mm<sup>2</sup> | 54 | 8:05 | -- |
| | S31 | Siemens | Basic VB17 | 1 $b=0$ s/mm<sup>2</sup> | 30 $b=1000$ s/mm<sup>2</sup> | 31 | 7:02 | 36 |
| | S55 | Siemens | Basic Skyra E11 & Prisma D13 | 7 $b=0$ s/mm<sup>2</sup> | 48 $b=1000$ s/mm<sup>2</sup> | 55 | 9:18 | 153 |
| | S127 | Siemens | Advanced Prisma VE11C | 13 $b=0$ s/mm<sup>2</sup> | 48 $b=1000$ s/mm<sup>2</sup> | 61 | 7:25* | 20 |
| ADNI2 | G46 | GE | Discovery MR750 & MR750w, Signa HDx & HDxt | 5 $b=0$ s/mm<sup>2</sup> | 41 $b=1000$ s/mm<sup>2</sup> | 46 | 7:00-10:00 | 59 |

\*reflects the time to acquire the full multi-shell protocol (127 volumes), not the single-shell subset

There is currently one multi-shell multiband protocol for Siemens Advanced Prisma scanners (S127). As ADNI3 is still in its early stages, GE and Philips protocols for multi-shell acquisition have not yet been finalized, so only 20 multi-shell scans were available for analysis at the time of writing. Here our goal was to evaluate single-shell dMRI indices across protocols, so we used a subsample of the 127 DWI volumes from the S127 multi-shell protocol to include only 13 *b*=0 and 48 *b*=1000 s/mm^2^ DWI volumes (removing 6 *b*=500 s/mm^2^ and 60 *b*=2000 s/mm^2^ volumes).

The Philips Basic Widebore R3 protocol (P36) included three *b*=2 s/mm^2^ volumes and one *b*=0 s/mm^2^, because Philips scanners cannot acquire more than one *b*=0 s/mm^2^. The Philips Basic Widebore (P33) was not a prescribed protocol, but rather acquired from Philips sites with a software version less than 5.0 that could not acquire the *b*=2 s/mm^2^ volumes.

### 2.4 dMRI preprocessing and scalar indices

All DWI were preprocessed using the ADNI2 diffusion tensor imaging (DTI) analysis protocol as in Nir et al., (2013). Briefly, we corrected for head motion and eddy current distortion, removed extra-cerebral tissue, and registered each participant’s DWI to the respective T1-weighted brain to correct for echo planar imaging (EPI) distortion. Details of the preprocessing steps may be found here: *https://adni.bitbucket.io/reference/docs/DTIROI/DTI-ADNI_Methods-Thompson-Oct2012.pdf*. All DWI and T1-weighted images were visually checked for quality assurance.

Scalar dMRI indices were derived from two reconstruction models: the single tensor model (DTI; Basser et al., 1994) and the tensor distribution function (TDF; Leow et al., 2009). From the single tensor model, FA^DTI^, AxD^DTI^, MD^DTI^, and RD^DTI^ scalar maps were generated. In contrast to the single tensor model, the TDF represents the diffusion profile as a probabilistic mixture of tensors that optimally explain the observed DWI data, allowing for the reconstruction of multiple underlying fibers per voxel, together with a distribution of weights, from which the TDF-derived form of FA (FA^TDF^) was calculated (Nir et al., 2017).

### 2.5 White matter tract atlas ROI summary measures

Images were processed as reported previously (Nir et al., 2013). Briefly, the FA image from the Johns Hopkins University single subject Eve atlas (JHU-DTI-SS; http://cmrm.med.jhmi.edu/cmrm/atlas/human_data/file/AtlasExplanation2.htm) was registered to each participant’s corrected FA image using an inverse consistent mutual information based registration (Leow et al., 2007); the transformation was then applied to the atlas WM parcellation map (WMPM) ROI labels (as shown in **Figure 7**; Mori et al., 2008) using nearest neighbor interpolation. Mean anisotropy and diffusivity indices were extracted from 24 WM ROIs total (**Table 3**): 22 ROIs averaged bilaterally, the full corpus callosum, and a summary across all ROIs (full WM).

**Table 3.** The following 24 ROIs from the JHU atlas (Mori et al., 2008) were analyzed.

|  |  |  |  |
| --- | --- | --- | --- |
| **CST** | Corticospinal tract | **SLF** | Superior longitudinal fasciculus |
| **CP** | Cerebral peduncle | **SFO** | Superior fronto-occipital fasciculus |
| **ALIC** | Anterior limb of internal capsule | **IFO** | Inferior fronto-occipital fasciculus |
| **PLIC** | Posterior limb of internal capsule | **SS** | Sagittal *stratum* |
| **RLIC** | Retrolenticular part of internal capsule | **EC** | External capsule |
| **PTR** | Posterior thalamic radiation | **UNC** | Uncinate fasciculus |
| **ACR** | Anterior *corona radiata* | **GCC** | *Genu* of corpus callosum |
| **SCR** | Superior *corona radiata* | **BCC** | Body of corpus callosum |
| **PCR** | Posterior *corona radiata* | **SCC** | Splenium of corpus callosum |
| **CGC** | Cingulum (cingulate gyrus) | **CC** | Full corpus callosum |
| **CGH** | Cingulum (hippocampal bundle) | **TAP** | *Tapetum* |
| **Fx/ST** | Fornix (*crus*) / *stria terminalis* | **Full WM** | Full white matter |

### 2.6 Comparing the ADNI2 and ADNI3 protocols in cognitively normal participants

#### 2.6.1 Sample sizes for the ADNI2 and ADNI3 cognitively normal participants

We evaluated the six ADNI3 protocols and the ADNI2 protocol using scans from cognitively normal (CN) participants only. Of 85 CN participants in ADNI2 with dMRI, 30 rolled over to ADNI3. To avoid duplication, and boost the number of scans available for each protocol, we did not include all these roll-over participants in the ADNI3 group. 26 CN roll-over participants were included in the ADNI3 group. Four CN roll-over participants were scanned with the S55 protocol, and due to the larger sample size already available for that protocol (N=156), we included these four in the ADNI2 group. In total, 59 out of 85 ADNI2 CN participants were included in the ADNI2 group and the remaining 26 were kept in the ADNI3 group for a total of 207 ADNI3 CN participants (see **Supplementary Table 3** for CN demographics by ADNI phase and protocol).

#### 2.6.2 Assessing age effects

In CN participants, multivariate random-effects linear regressions were used to assess whether to assess whether WM dMRI indices from each ADNI protocol individually were associated with age, controlling for sex and age*sex interactions as fixed variables, and acquisition site as a random variable. dMRI indices for the CN group were subsequently pooled across ADNI3 protocols (N=207), or ADNI3 and ADNI2 protocols (N=266) and tested for associations with age using an analogous model, but with protocol and acquisition site as nested random variables (e.g., 8 sites used protocol GE54, and 3 sites used protocol P33, so the acquisition site grouping variable is nested within the protocol grouping variable). We used the false discovery rate (FDR) procedure to correct for multiple comparisons (*q* = 0.05; Benjamini and Hochberg, 1995) across the 24 ROIs assessed for each dMRI index. Regions that survive a more stringent Bonferroni correction at an alpha of 0.05 (*p* ≤ 0.05/24=0.0021) are also shown in the Supplements.

#### 2.6.3 Effect of protocol on dMRI indices

In CN participants, we tested for significant differences in dMRI indices between the seven ADNI protocols using analyses of covariance (ANCOVAs), adjusting for age, sex, and age*sex interactions as fixed variables and acquisition site as a random variable. For each dMRI index, we used FDR to correct for multiple comparisons across the 24 ROIs assessed. Pairwise tests were performed to directly compare protocols. In total, there were 504 tests per dMRI index: 24 ROIs * 21 pairs of protocol comparisons (protocol 1 vs 2, protocol 1 vs 3, etc). As before, we used FDR to account for multiple comparisons.

#### 2.6.4 dMRI harmonization with ComBat

ComBat uses an empirical Bayes framework to reduce unwanted variation in multi-site data due to differences in acquisition protocol, while preserving the desired biological variation in the data (Fortin et al., 2017). In the CN participants from ADNI2 and ADNI3, we ran ComBat on each of the dMRI indices, including age, sex, age*sex, and information from all 24 ROIs to inform the statistical properties of the protocol effects. Random-effects regressions tested for dMRI microstructural associations with age, covarying for sex and age*sex as fixed variables and site as a random variable; ANCOVAs and pairwise tests of dMRI differences between protocols were repeated for the harmonized ROI data.

### 2.7 Clinical assessments and their relation to pooled ADNI3 diffusion indices

Multivariate random-effects linear regressions were used to test associations between 5 dMRI indices in each of the 24 WM ROIs and the three cognitive measures (ADAS, MMSE, CDR-sob), and with diagnosis. Due to the limited available sample size for AD participants (N=22), and their uneven distribution across the acquisition protocols tested here, we compared only groups of people with CN and MCI diagnoses. Age, sex, and age*sex interactions were controlled for as fixed effects, and the protocol and acquisition site were modeled as nested random variables. FDR was again used to correct for 24 ROI tests (*q* = 0.05; Benjamini and Hochberg, 1995), in addition to Bonferroni corrections (*p* ≤ 0.05/24=0.0021) available in the Supplements. Effect sizes for associations were determined using the *d*-value standardized coefficient (Rosenthal and Rosnow, 1991):

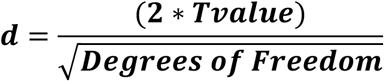

## 3 Results

### 3.1 ADNI2 and ADNI3 protocols in cognitively normal participants

#### 3.1.1 Age effects in cognitively normal participants from ADNI2 and ADNI3 protocols

When data were pooled across ADNI2 and ADNI3, significant associations with age were detected throughout the WM. **Figure 1a** shows effect sizes for ROIs significantly associated with age after FDR multiple comparisons correction (tabulated results and more stringent Bonferroni thresholds are shown in **Supplementary Table 4**). Lower FA^TDF^ and higher diffusivity indices were significantly associated with older age in all 24 ROIs. For FA^DTI^, 22 ROIs were significantly associated with age. The largest effect size was detected with FA^TDF^ in the Fornix *(crus) / stria terminalis* (Fx/ST; *d* = −1.459*; p* = 5.07×10^−21^). The Fx/ST, corpus *callosum genu* (GCC) and full WM consistently showed one of the 10 largest effect sizes across dMRI indices.

**Figure 1.**
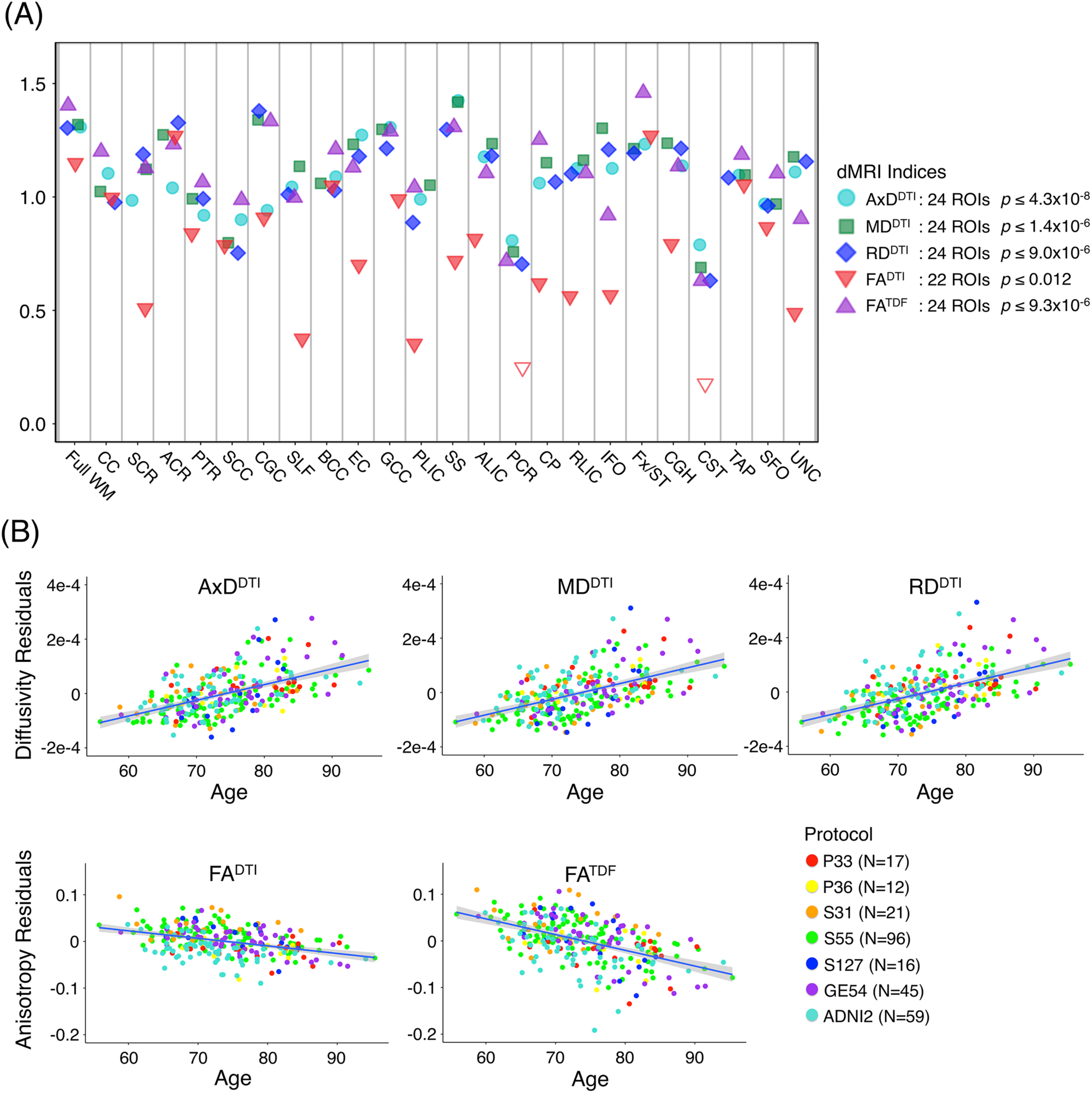
**(A)** For each dMRI index, the absolute values of effect sizes (*d*-value) are plotted for regional WM microstructural associations with age when all ADNI3 dMRI data are pooled, adjusting for any site or protocol effects. For each test, we note the number of significant ROIs, as indicated by filled shapes, and the corresponding FDR significance *p*-value threshold (*q* = 0.05). See **Supplementary Table 4** for complete tabulated results. **(B)** Here, we plot the residuals of diffusivity and anisotropy indices in the full WM (*y*-axis) against age (*x*-axis) after regressing out the effects of sex in CN participants from each protocol separately. Individual level residuals from each protocol are plotted with a different color. Despite protocol differences, age effects are evident across protocols.

The mean ages of the CN participants assessed in the two phases of ADNI were significantly different (*p* = 0.049; ADNI2 mean age: 72.4±6.6 yrs; ADNI3 mean age: 74.5±7.4 yrs; demographics in **Supplementary Table 3**). Pairwise tests comparing the mean age of CN participants scanned with each protocol also showed significant differences between those scanned with S31 and two other protocols: GE54 and S31 (*p =* 0.026); P33 and S31 (*p =* 0.0037). Due to differences in age and sample size between protocols and phases, effect sizes could not be directly compared (Button et al., 2013), but the directions of associations with age were largely consistent for ADNI2 and ADNI3 phases separately, and each ADNI3 protocol (**Figures 1–2**). Each ADNI protocol showed directionally consistent associations in more than 89% of tests (24 ROIs * 5 dMRI indices), except for P36 which was consistent in 81%, but had the smallest sample size (N=12; **Figure 2b; Supplementary Tables 5-11**). FA^TDF^ and all three diffusivity indices were consistent in ≥ 96% of tests (24 ROIs * 8 protocols/phases), while FA^DTI^ was only consistent in 88% of tests. Most of the associations detected in the unexpected direction for each protocol were driven by FA^DTI^. None of the associations in the unexpected direction were significant after multiple comparisons correction, and only 2 had a *p* ≤ 0.05.

**Figure 2.**
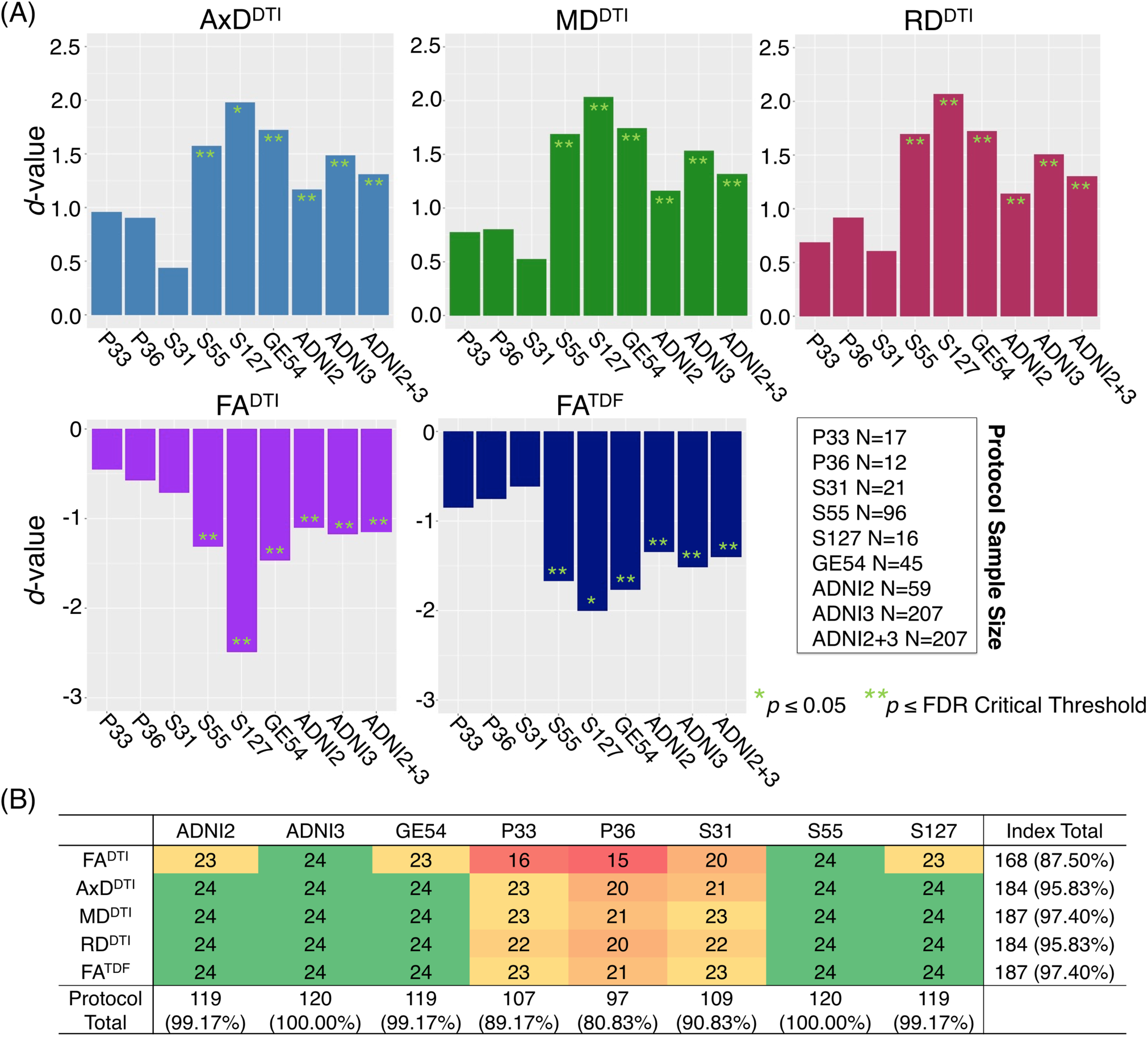
**(A)** Effect sizes (*d*-value) for each ADNI protocol and phase show the direction of dMRI associations with age in the full WM are consistent. Due to differences in age and sample size between protocols and phases, effect sizes could not be directly compared. **(B)** For each protocol and phase, the number of ROIs (out of 24), that show the expected association direction, regardless of significance, are reported for each dMRI index, revealing consistent associations across tests, except for protocol P36 which has the smallest sample size, and FA^DTI^, which shows the smallest effect sizes and fewest significant associations across protocols when pooled.

Figure 2 shows consistent associations in the full WM by protocol. As demographic and sample size variability between protocols affect detected effect sizes, we also evaluated full WM dMRI associations with age in an age- and sex-matched subset of 12 participants from each protocol (total N=84; demographics in **Supplementary Table 3**); a comparison of the effect sizes between protocols suggests that the protocols with greatest total number of diffusion-weighted (*b*=1000 s/mm^2^) and non-diffusion sensitized (*b*_0_) gradients may detect larger effects (S127 followed by S55; **Supplementary Figure 1**).

#### 3.1.2 Effect of protocol on dMRI indices from cognitively normal controls

The influence of dMRI acquisition protocol on mean values of the diffusion indices is evident in boxplots of dMRI indices in the full WM for each protocol. When modeling the mean full WM values for each diffusion index, the residuals of the statistical model become closer to 0 after fitting the effect of protocol and site (nested as a random variable with age, sex, and age*sex interactions as fixed effects) than when we plot the residuals of just age, sex, and age*sex interactions (**Figure 3**).

**Figure 3.**
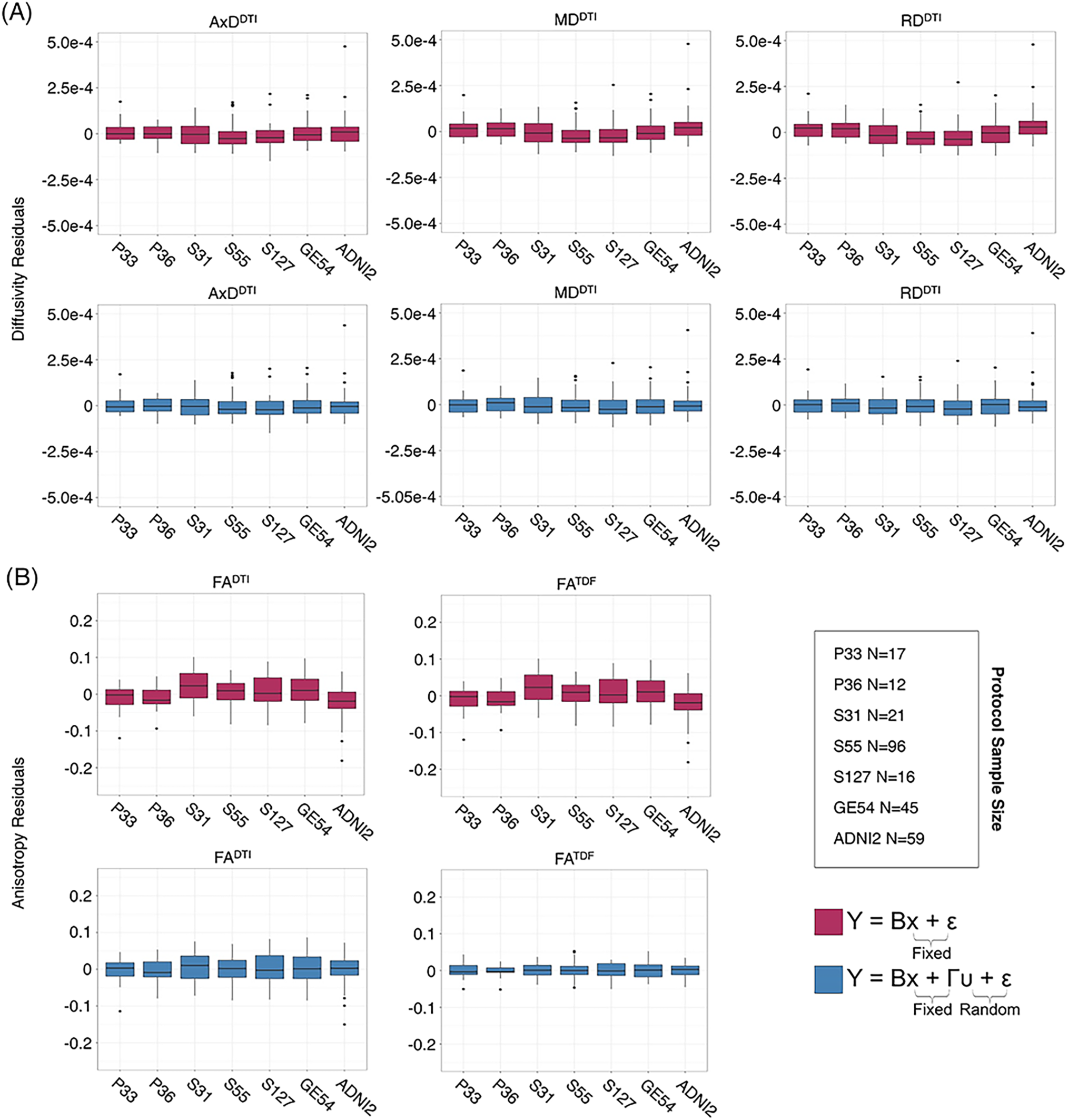
Full WM mean **(A)** AxD^DTI^, MD^DTI^, and RD^DTI^, and **(B)** FA^DTI^ and FA^TDF^ residuals for each protocol after fitting effects of age, sex, and age*sex interactions are plotted here in the top rows (*red*). Protocol has an effect on anisotropy and diffusivity measures. The lower panels (*blue*) show residuals after additionally fitting protocol and site as nested random-effects, after which the residuals across protocols are closer to 0.

ANCOVAs and pairwise tests for each ROI suggest there are significant differences between protocols for all 5 dMRI indices across most ROIs (**Figure 4**). ANCOVAs revealed significant protocol differences for 22 ROIs for FA^DTI^ and FA^TDF^, with the highest overall effect size detected in the anterior limb of the internal capsule (ALIC) and the external capsule (EC) for FA^DTI^ (ALIC: *d =* 0.648; EC: *d* = 0.652). AxD^DTI^ had the smallest effect size, overall, in the splenium of the corpus callosum (SCC; *d* = 0.106), and only 13 ROIs showed significant AxD^DTI^ differences between protocols.

**Figure 4.**
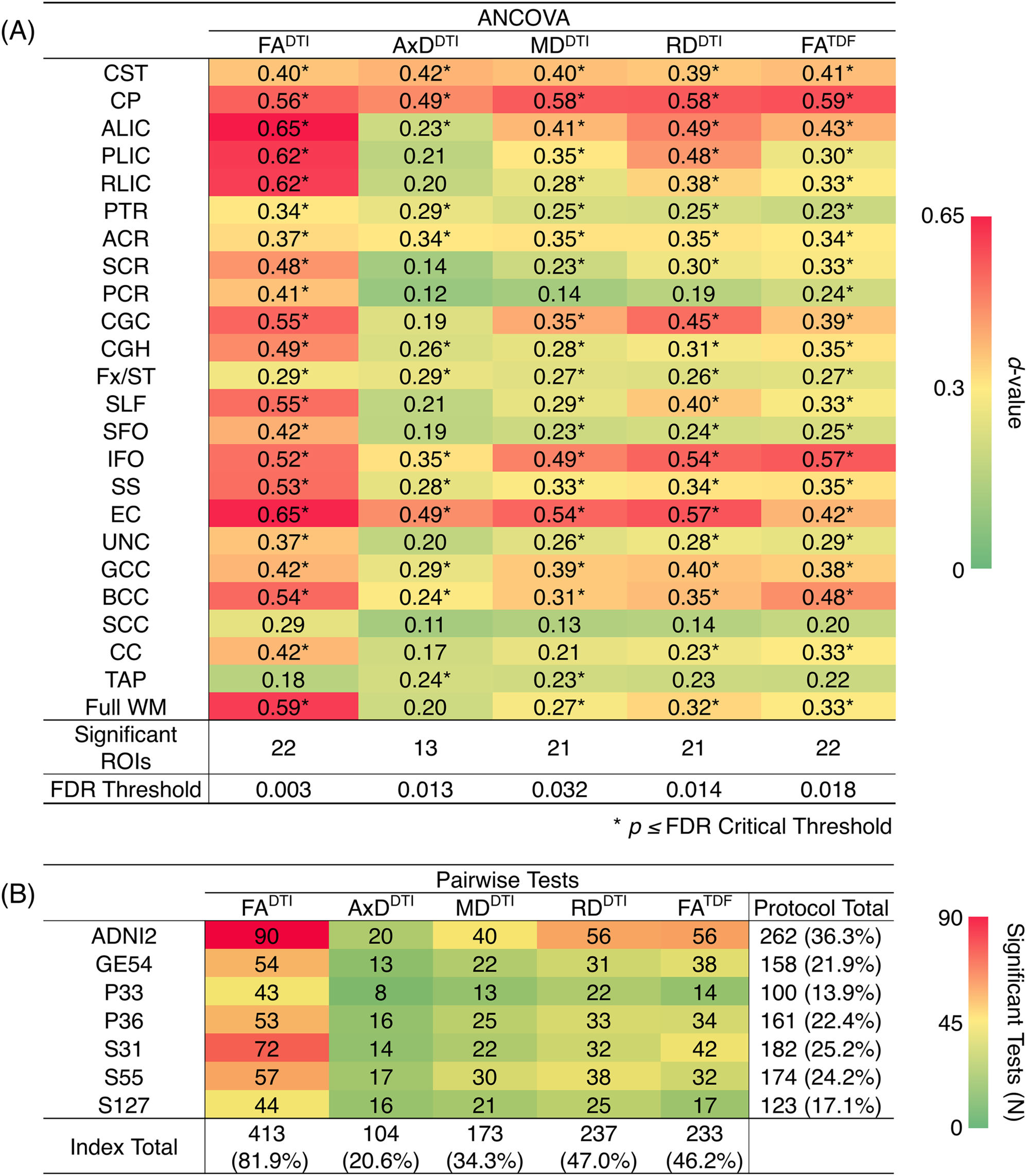
**(A)** *d*-values from the ANCOVAs assessing differences in dMRI indices between protocols, for each of the 24 ROIs; FA^DTI^ shows the greatest significant differences (largest *d*-values; *dark red*) between protocols and AxD^DTI^ the fewest (*dark green*). **(B)** We report the number of times each protocol and each dMRI index showed significant differences in pairwise tests between protocols (out of 504 tests per index and 720 tests per protocol); AxD^DTI^ was the most stable dMRI index across protocols, while FA^DTI^ was the least stable.

In pairwise analyses, AxD^DTI^ was the most stable index across protocols, as significant protocol differences were detected in only 20.6% of pairwise tests (24 ROIs * 21 pairwise tests), compared to FA^DTI^, the most variable index, which showed significant protocol differences in 81.9% of tests (**Figure 4b**). ADNI2 was the most divergent protocol across dMRI indices, showing differences in 36.3% of tests.

#### 3.1.3 Diffusion MRI harmonization with ComBat

After using ComBat to harmonize dMRI indices across protocols, ANCOVAs revealed that significant protocol differences in dMRI indices were all but eliminated across ROIs (**Supplementary Figure 2a**); significant protocol differences were detected only in the CST, for each of the dMRI indices. The number of pairwise tests for which each protocol showed significant differences in dMRI indices decreased by 93.8% with ComBat (**Supplementary Figure 2b**).

After harmonization, we still detected significant associations between age and dMRI indices from ADNI2 and ADNI3 pooled in the same number of ROIs (**Supplementary Table 12**). ComBat correction did not significantly change effect sizes, while correcting for effects of protocol (**Supplementary Figure 3**). In **Figure 5** we show effect sizes before and after harmonization with ComBat in the Full WM, Fx/ST, and GCC, the three ROIs that consistently showed one of the 10 largest effect sizes for associations with age across all five diffusion indices (for changes by protocol see **Supplementary Figures 4-6**). As harmonization with ComBat did not improve or change results found with random-effect linear regressions, we proceeded to test clinical associations without applying a ComBat transformation.

**Figure 5.**
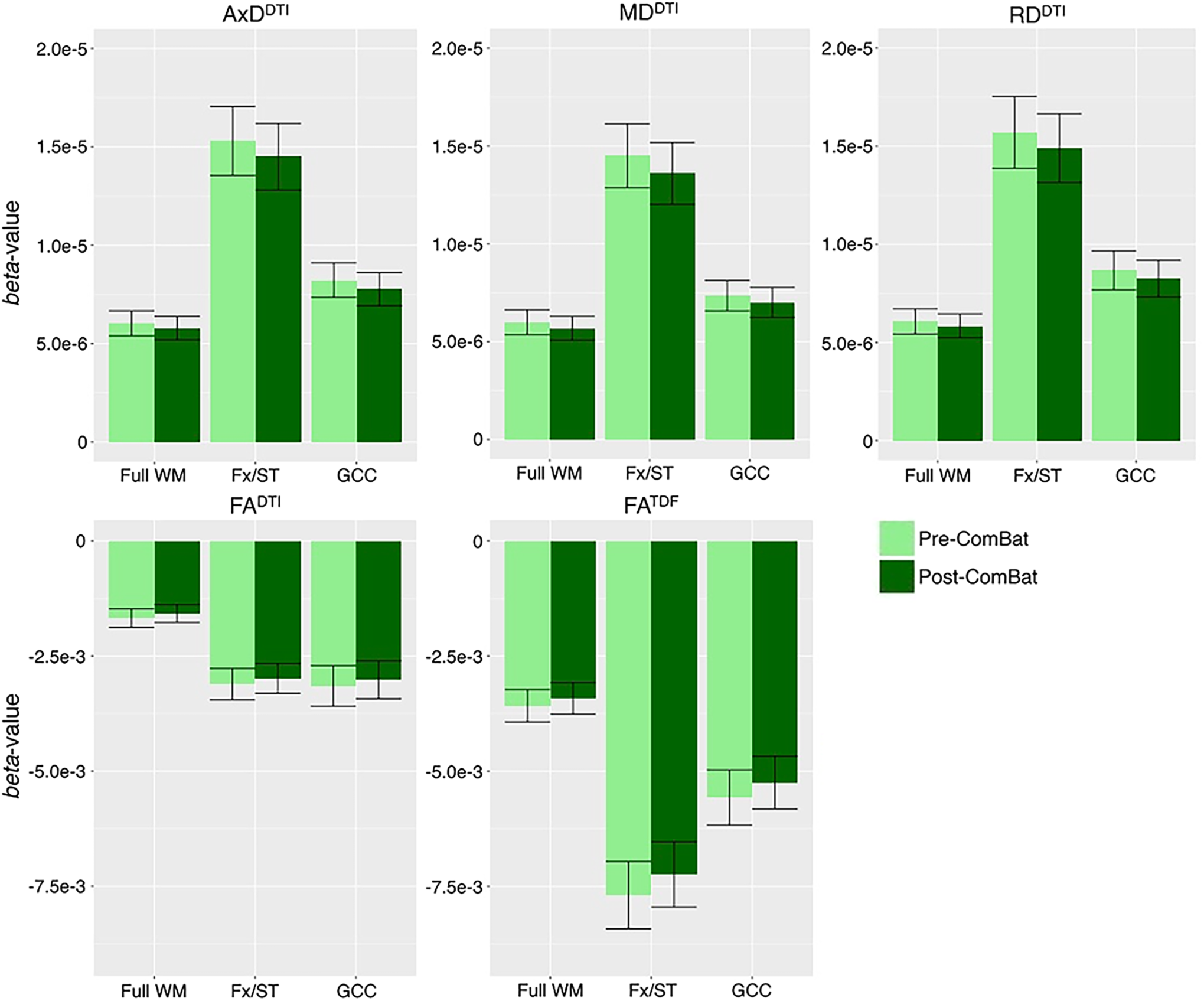
*Beta*-values and error bars representing standard error from the association between each diffusion index and age in CN participants, before and after ComBat harmonization. We show the three ROIs that consistently showed one of the 10 largest effect sizes for associations with age across all five diffusion indices (see **Supplementary Figure 3** for all ROIs). Compared to pre-ComBat analyses, effect sizes are marginally different across indices, but still within the standard error.

### 3.2 Cognitive measure associations with pooled ADNI3 dMRI indices

Pooling data across ADNI3, we detected significant associations between all three cognitive measures and regional dMRI indices throughout the WM. Greater cognitive impairment was associated with lower anisotropy and higher diffusivity. **Figure 6a-c** shows effect sizes for ROIs significantly associated with each cognitive measure after FDR multiple comparisons correction (for tabulated results and more stringent Bonferroni corrections, please see **Supplementary Tables 13-15**). Across tests (5 dMRI indices * 3 cognitive measures), the hippocampal-cingulum (CGH), fornix *(crus)* / *stria terminalis* region (Fx/ST), and the full WM consistently showed one of the 10 largest effect sizes (see **Supplementary Figures 7-9** for associations with indices in the CGH, Fx/ST, and full WM, by protocol). In 14 of 15 tests, the CGH consistently showed one of the top two largest effect sizes (CGH FA^DTI^ association with CDR-sob was the third largest), along with the uncinate fasciculus (UNC), which was top two in 12 of 15 tests (while significant, cognitive associations with UNC FA^DTI^ never showed one of the largest effect sizes).

**Figure 6.**
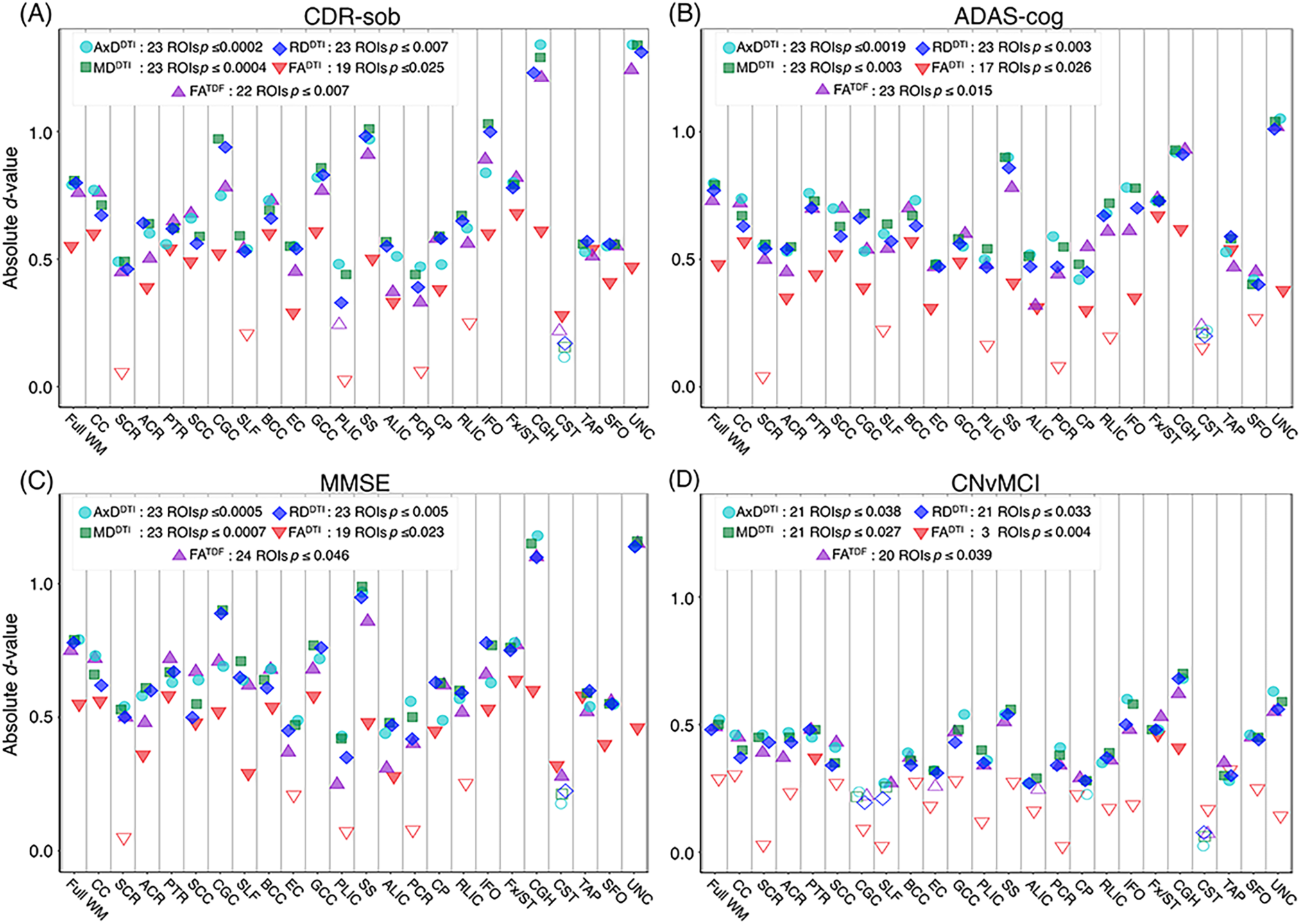
For each dMRI index, the absolute values of effect sizes (*d*-value) are plotted for regional WM microstructural associations with clinical measures. Lower anisotropy and higher diffusivity were significantly associated with **(A)** higher CDR-sob, **(B)** lower MMSE, **(C)** higher ADAS-cog, and **(D)** an MCI diagnosis, when all ADNI3 dMRI data are pooled, adjusting for any site or protocol effects. For each test, we note the number of significant ROIs, as indicated by filled shapes, and the corresponding FDR significance *p*-value threshold (*q* = 0.05). See **Supplementary Tables 13-16** for complete tabulated results.

FA^DTI^ showed significant associations in the fewest ROIs: 55 out of 72 tests (76.4%; 24 ROIs * 3 cognitive measures) were significant. FA^TDF^ showed more widespread associations with cognitive measures throughout WM ROIs: 69 out of 72 tests (94.4%; 24 ROIs * 3 cognitive measures) were significant. Effect sizes were consistently lower for FA^DTI^ than for the other dMRI indices, across all 3 cognitive measures; the largest FA^DTI^ effect size was consistently found in the Fx/ST, followed by the CGH or the GCC. The strongest FA^DTI^ association was in the Fx/ST with CDR-sob (*d* = −0.681*, p =* 7.01×10^−8^). Compared to FA^DTI^, FA^TDF^ showed larger effect sizes; across cognitive tests, the strongest FA^TDF^ associations were detected in the uncinate fasciculus (UNC) with CDR-sob (*d =* −1.244; *p =* 1.39×10^−20^), followed by the CGH (*d =* −1.213; *p =* 8.86×10^−20^). CDR-sob effect sizes for FA^DTI^ and FA^TDF^ in the CGH, UNC, Fx/ST, and full WM are depicted by protocol in **Supplementary Figure 10**, revealing consistently larger effect sizes for FA^TDF^ across protocols.

Cognitive associations with all of the diffusivity indices were widespread: significant associations were detected in 207 out of 216 tests (95.8%; 24 ROIs * 3 cognitive measures * 3 diffusivity indices). Regional measures of AxD^DTI^ consistently showed the largest effect sizes across all cognitive measures (CDR-sob and the UNC: *d =* 1.344*, p =* 3.13×10^−23^; MMSE and the CGH: *d =* −1.178*, p =* 7.87×10^−19^; ADAS-cog and the UNC: *d =* 1.048*, p =* 1.09×10^−13^).

Of the three cognitive measures, CDR-sob associations showed the largest effect sizes across dMRI indices (in the UNC followed by the CGH for all indices except FA^DTI^); the largest effect sizes across all tests were detected with AxD^DTI^ (UNC: *d =* 1.344) and MD^DTI^ (UNC: *d =* 1.34*2, p =* 3.47×10^−23^). **Figure 7** shows the distribution of the effect sizes for CDR-sob throughout the brain. Temporal lobe regions (UNC, CGH, IFO, SS) frequently showed greatest effect sizes (for ADAS-cog and MMSE figures, see **Supplementary Figures 11-12**). Effect size was not correlated with ROI size (**Supplementary Figure 13**), consistent with prior studies of other disorders (Kelly et al., 2018).

**Figure 7.**
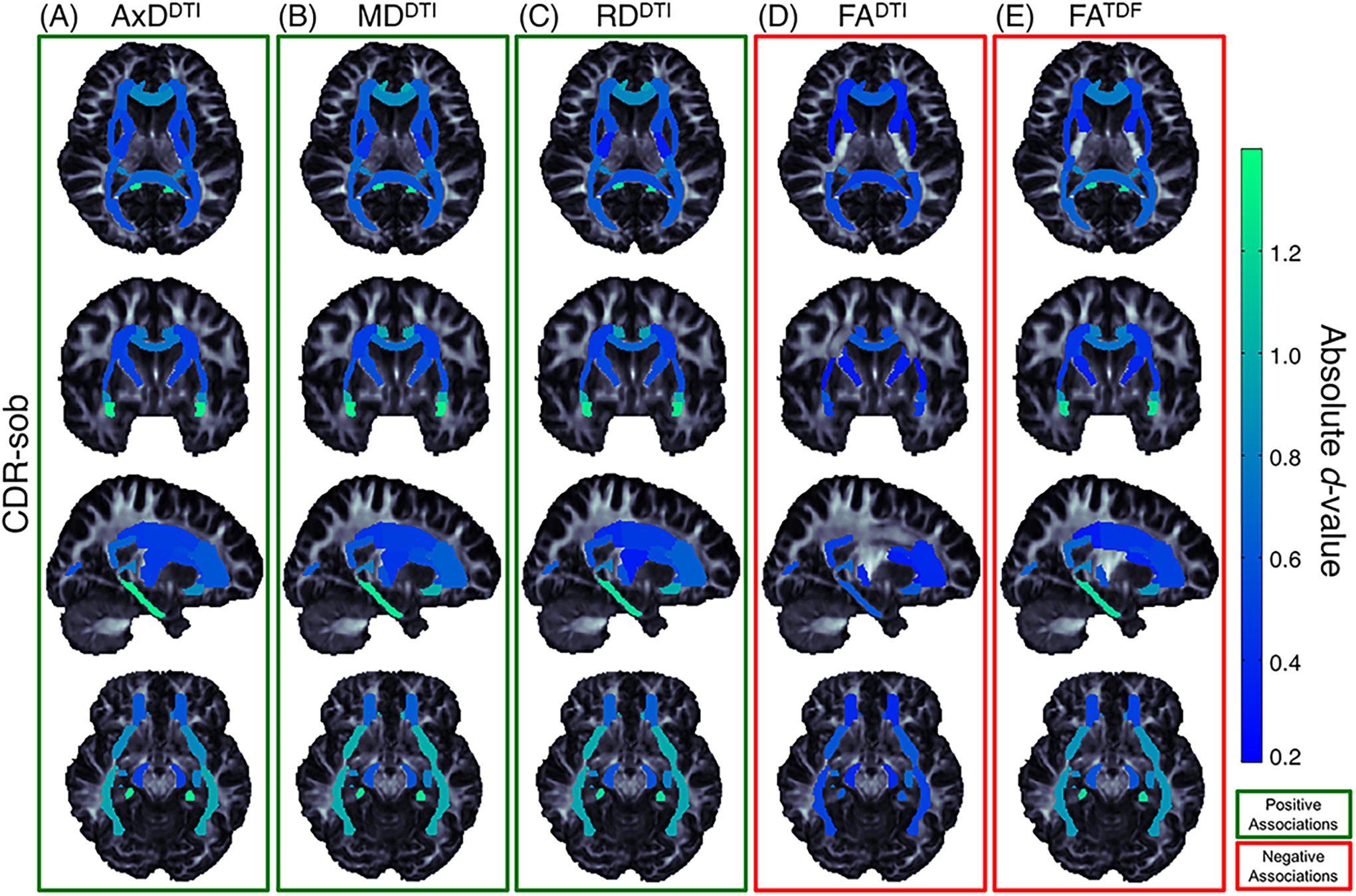
Effect size (absolute *d*-value) maps of WM regions that show significant associations with CDR-sob - the cognitive measure with the largest effect sizes - reveal widespread associations throughout the WM, with particularly strong associations in the temporal lobes (SS, IFO, UNC, and CGH; *light green* regions show the largest effect sizes). As expected, positive associations were detected between CDR-sob and **(A)** AxD^DTI^ (FDR critical threshold *p* = 1.78×10^−4^) **(B)** MD^DTI^ (FDR critical threshold *p* = 3.64×10^−4^) and **(C)** RD^DTI^ (FDR critical threshold *p* = 6.92×10^−3^); higher diffusivity was associated with greater cognitive impairment. Lower **(D)** FA^DTI^ (FDR critical threshold *p* = 0.025) and **(E)** FA^TDF^ (FDR critical threshold *p* = 7.73×10^−3^) were also associated with greater impairment, but FA^DTI^ associations were detected in fewer regions with weaker effect sizes compared to FA^TDF^.

### 3.3 CN vs MCI diagnosis associations with pooled ADNI3 dMRI indices

For each diffusion index, **Figure 6d** shows the significant regional effect sizes for differences between CN and MCI participants. Widespread diffusivity differences were detected, with significantly higher diffusivity in MCI participants in 21 out of 24 ROIs (**Supplementary Table 16** and **Supplementary Figure 14**). Only three regions showed significantly lower FA^DTI^ in MCI participants – Fx/ST (*d* = −0.460; *p* = 3.89×10^−4^), CGH (*d =* −0.410; *p* = 1.53×10^−3^), and the posterior thalamic radiation (PTR; *d =* 0.367; *p* = 4.55×10^−3^). On the other hand, FA^TDF^ was significant in 20 out of 24 ROIs, similar to diffusivity indices. FA^TDF^ and diffusivity indices in the CGH showed the largest effect sizes (AxD^DTI^ *d =* 0.681; *p =* 2.26×10^−7^, MD^DTI^ *d =* 0.700*; p =* 1.15×10^−7^; RD^DTI^ *d =* 0.679*; p =* 2.41×10^−7^; FA^TDF^ *d = -*0.622*; p =* 2.00×10^−6^).

For all three cognitive measures, and in the comparison between CN and MCI participants, the CGH and Fx/ST were the only regions that survived multiple comparisons correction across all dMRI indices. The Fx/ST always had the largest effect size in FA^DTI^ tests. The UNC showed either the first or second largest effect size (alternating with CGH) across diffusivity indices and FA^TDF^ tests, but was significant only for cognitive measure associations with FA^DTI^ (i.e., 3 of 4 clinical tests).

## 4 Discussion

This study has three main findings: (1) When data were pooled from the six available diffusion MRI protocols used in ADNI3, anisotropy and diffusivity indices showed robust associations with MCI diagnosis, and with three common cognitive measures: MMSE, ADAS-cog, and CDR-sob; (2) When using a higher-order diffusion model, the tensor distribution function (TDF), the derived measure of anisotropy (FA^TDF^) showed stronger and more widespread associations with clinical impairment than the standard DTI anisotropy measure (FA^DTI^); (3) Despite significant differences in protocols, for each dMRI index, we were able to detect consistent associations with clinical measures in ADNI3 participants, and age in ADNI2 and ADNI3 CN participants.

Accumulation of amyloid plaques and neurofibrillary tangles (NFT) in the brain (Braak & Braak, 1991; Braak & Braak 1996; Frank et al., 2003, Shaw et al., 2007) can directly impact WM (Lee et al., 2004; Roth et al., 2005), promoting myelin degeneration and axonal loss (Braak and Braak 1996; Kneynsberg et al., 2017). While many factors drive anisotropy and diffusivity measures from DTI, higher anisotropy values may indicate, in part, more coherent intact axons, while lower anisotropy and higher diffusivity may reflect factors such as axonal injury and demyelination, among other factors (Beaulieu et al., 2002; Song et al., 2003; Song et al., 2005; Harsan et al., 2006; Le Bihan and Johansen-Berg 2012; Kantarci et al., 2017; Moore et al., 2018). In this paper, lower anisotropy values and higher diffusivity values were correlated with clinical impairment, most strongly in the hippocampal-cingulum and uncinate fasciculus. Along with the full WM, reflecting global WM effects, the largest effect sizes were most frequently detected in the hippocampal-cingulum and fornix *(crus)* / *stria terminalis*, WM bundles connecting hippocampal and parahippocampal regions to the rest of the brain, consistent with patterns of AD pathology. The histopathological validity of these findings has been supported, specifically in a recent study that compared NFT stages in autopsy material along with ante-mortem MRI. Elevated MD^DTI^ and lower FA^DTI^ significantly correlated with higher postmortem NFT stage, particularly in the *crus* of the fornix, the ventral cingulum tracts, the precuneus, and entorhinal WM (Kantarci et al., 2017).

The participants recruited for ADNI3 tend to be younger and healthier, on average, than those in ADNI2, as they were recruited with the intention of studying the transition from CN to mild AD (Jack et al., 2015). With few AD patients enrolled so far in ADNI3, the primary focus of this paper was to assess three cognitive assessments (ADAS-cog, CDR-sob, and MMSE), and to compare CN to MCI participants. MCI is now the focus of intense research, as it is essential to find ways to clinically categorize the transitional stages between normal aging and AD to evaluate targeted treatments, as pathophysiological mechanisms may differ or change throughout the course of AD (Mueller et al., 2005). As in our prior analysis of ADNI2 (Nir et al., 2013), FA^DTI^ was the least sensitive DTI measure. In ADNI3, AxD^DTI^ and MD^DTI^ showed the largest effect sizes. Lower FA^DTI^ and higher MD^DTI^ are most frequently reported in studies of AD (Kavcic et al., 2008; Clerx et al., 2012; Nir et al., 2013; Maggipinto et al., 2017; Mayo et al., 2017), but AxD^DTI^ may be more sensitive to unspecific microscopic cellular loss earlier in the disease (O’Dwyer et al., 2011), perhaps making it more sensitive in the healthier participants of the ADNI3 dataset. Similarly, in ADNI2, AxD^DTI^ was the most sensitive to differences between CN and MCI diagnoses (Nir et al., 2013).

Among the three cognitive assessments, CDR-sob showed the strongest correlations with dMRI indices, in line with prior ADNI brain imaging studies (Hua et al., 2009; Nir et al., 2013). The largest of these effects were found in temporal WM tracts including the hippocampal-cingulum, uncinate fasciculus, sagittal stratum, and inferior fronto-occipital fasciculus. These are all regions that show early degenerative changes in MCI and AD (Mielke et al., 2009; Nir et al., 2013; Maggipinto et al., 2017; Powell et al., 2018). While associations with clinical impairment were detected throughout the WM, the region that most frequently showed the lowest effect sizes and was significant in only 3 of the 20 clinical tests, was the corticospinal tract (CST). However, the CST ROI from the JHU WMPM atlas is limited to a small region in the inferior portion of the brain and has been shown to be the least reliable and reproducible ROI (Jahanshad et al., 2013; Acheson et al., 2017), suggesting alternate approaches, such as tractography-based evaluations (Jin et al., 2017), or the use of the probabilistic JHU atlas (Hua et al., 2008), may be more appropriate for studying the CST. Our analysis focused on white matter microstructure, but future work assessing tract geometry and properties of anatomical brain networks using tractography may reveal more detailed information. The validation and harmonization of tractography methods and derived network metrics is a vast field of research with active ongoing work (Maier-Hein et al., 2017).

DTI is widely recognized as a useful tool for studying neurodegenerative disorders such as AD (Oishi et al., 2011; Müller and Kassubek, 2013; Abhinav et al., 2014; Acosta-Cabronero et al., 2014; Maggipinto et al., 2017). However, at the spatial resolutions now used, a single voxel typically captures partial volumes of different tissue compartments, e.g., the intra- and extra-cellular compartments, the vascular compartment, the CSF and myelin; each affects water diffusion and the MR signal. The DTI model cannot differentiate these components or even crossing fibers (Tuch et al., 2002; Jbabdi et al., 2010), which are estimated to occur in up to 90% of WM voxels at the typical dMRI resolution (Descoteaux et al., 2009; Jeurissen et al., 2013). In healthy tissue with crossing fibers, the DTI model may show low FA. FA^DTI^ may paradoxically appear to increase in regions where crossing fibers deteriorate in neurodegenerative diseases such as AD (Douaud et al., 2011). FA^TDF^ addresses this limitation even in low angular resolution data (Nir et al., 2017). Here, compared to FA^DTI^, FA^TDF^ showed more widespread associations with cognitive measures and diagnosis throughout WM ROIs: FA^TDF^ was significant in 89 of the 96 tests (92.7%; 24 regions in 4 clinical tests), while FADTI was only significant in 58 (60.4%). The greatest difference was seen for diagnostic associations (CN vs MCI): FATDF was significant in 20 out of 24 ROIs while FADTI was only significant in 3. FATDF also showed stronger effect sizes across the protocols, suggesting that tensor limitations have likely confounded previous diffusion studies of cognitive decline that have found little or no effects with FA (Acosta-Cabronero et al., 2010). Recently proposed biophysical models of brain tissue may help to relate diffusion signals directly to underlying microstructure and different tissue compartments (Harms et al., 2017). We may be able to further disentangle questions of orientation coherence (dispersing and ‘kissing’ fibers), fiber diameter, fiber density, membrane permeability, and myelination, which all influence classic anisotropy and diffusivity measures derived from DTI. Several AD studies have already used multi-shell protocols to compute diffusion indices from models that do not assume mono-exponential decay, such as diffusion kurtosis imaging (DKI; Jensen et al., 2005; Chen et al., 2017; Cheng et al., 2018; Wang et al., 2018), and multi-compartment models such as neurite orientation dispersion and density imaging (NODDI; Zhang et al., 2012; Colgan et al., 2016; Slattery et al., 2017; Parker et al., 2018). To date, approximately 20 participants in ADNI have been scanned with multi-shell diffusion protocols; in a future report, we will relate these measures to those examined here.

Large-scale, multi-site neuroimaging studies can increase the power of statistical analyses and establish greater confidence and generalizability for findings. Most multi-site neuroimaging studies are susceptible to variability across sites. Variability in dMRI studies is due in part to heterogeneity in acquisition protocols, scanning parameters, and scanner manufacturers (Zhu et al., 2009; Zhu et al., 2011; Zhu et al., 2018). Anisotropy and diffusivity maps are affected by angular and spatial resolution (Alexander et al., 2001; Kim et al., 2006; Zhan et al., 2010), the number of DWI directions (Giannelli et al., 2009), and the number of acquired *b*-values (Correia et al., 2009). All five dMRI indices were significantly different between protocols; AxD^DTI^ was the most stable index, while FA^DTI^ was the least stable, reflective of their performance in detecting associations with cognitive measures. ADNI2 was the most divergent protocol across dMRI indices, perhaps due to the larger voxel size in ADNI2 (2.7 mm versus 2.0 mm isotropic voxels used in ADNI3). This is consistent with the notion that DTI measures vary with voxel size due to partial voluming (Zhan et al., 2013). Despite differences in protocols, the directions of associations were consistent across protocols.

ADNI3 extends dMRI acquisitions across scanner manufacturers and platforms to maximize the number of participants scanned with dMRI; this makes it necessary to account for site-related heterogeneities and confounds in analytical models where data are pooled. Multi-site dMRI studies are becoming increasingly common, and new data harmonization methods to adjust for site and acquisition protocol are being developed and tested. A thorough investigation of dMRI harmonization methods is now possible with ADNI3, one of the few publically available multi-site datasets acquired with multiple protocols. As regional dMRI measures are available for download as part of the ADNI database, we highlight two ways that the data may be pooled across sites: 1) performing statistical analyses with nested random-effects models to account for site and acquisition protocol differences, and 2) harmonizing the derived regional measures before aggregating the data across sites. In a preliminary analysis, we showed that one harmonization method performed on these regional measures, ComBat, reduced cross-site differences in dMRI indices, while preserving biological relationships with age in CN controls. The only region where differences remained after ComBat, was the CST, the ROI with the weakest associations with clinical measures, and previously identified as least reliable (Acheson et al., 2017). In Fortin et al. (2017), compared to other harmonization methods, ComBat increased the number of voxels where significant associations between age and FA^DTI^ or MD^DTI^ were detected. Here, the number of significant ROIs and the magnitude of effect sizes were comparable for ComBat and nested random-effects model approaches. This discrepancy between our findings and that of Fortin et al, may be due to differences between studies: 1) ADNI3 includes more sites and protocols, 2) the number of ROIs is far less than the number of participants, and 3) the age effects in the elderly populations tested here are stronger than the effects tested in adolescents in Fortin et al. When effects are more readily detected, one harmonization approach may not be more advantageous than others. In addition to exploring additional harmonization techniques, future work should evaluate voxel-wise ComBat approaches and the effects of harmonization beyond CN participants (i.e., across the entire ADNI cohort).

In addition to ComBat, a number of harmonization approaches have recently been proposed at various stages of analysis (Tax et al., 2018; Zhu et al., 2018). Site differences can be accounted for at the time of overall group inference, such as with the random-effects regression level correction used here, or by using a meta-analysis approach in lieu of pooling data (Thompson et al., 2014). The data may also be transformed prior to multi-site group-level statistics. Some methods, such as ComBat and RAVEL, use the distribution of derived features, such as diffusivity and anisotropy measures (Fortin et al., 2016, 2017). Alternatively, several proposed methods use information from the raw image to adjust for acquisition variability (Zhu et al., 2018). For example, Kochunov et al. (2018) calculated the signal to noise ratio for each protocol and include it in their regression models. Mirzaalian et al. (2018) use voxel-wise spherical harmonic residual networks to derive local correction parameters. Finding the best method to harmonize dMRI data is an active topic at ‘hackathons’ and technical challenges; in 2017 and 2018, the International Conference on Medical Image Computing and Computer Assisted Intervention (MICCAI) hosted a computational diffusion MRI challenge to explore approaches for data harmonization. With so many available approaches, the preliminary random-effects regression and ComBat results from this paper serve as a first step towards future work establishing robust approaches for combining data in ADNI3 and other multi-site studies.

The current study is limited in that the sample sizes, and sample demographics, available for each protocol vary, complicating direct comparison of the protocols (Button et al., 2013). A matched comparison might be possible if a group of participants or a phantom were scanned using every protocol. Even so, separating protocol differences from differences in scanner manufacturer is difficult. We also could not directly compare all diagnostic groups in ADNI3, as few participants with AD were scanned.

A more complete picture of brain changes in aging and AD would include imaging metrics from other modalities, such as perfusion imaging, resting state functional MRI (Wang et al., 2017), and radiotracer methods such as FDG-PET (Popuri et al., 2018), or amyloid- and tau-sensitive PET (Grothe et al., 2017; Phillips et al., 2018). Genetic and other ‘omics’ data could be analyzed as well, and may help to predict diagnostic classification and brain aging, when combined with other neuroimaging markers (Ding et al., 2018; Kauppi et al., 2018). While these data are all being collected as part of ADNI3 and other studies of brain aging, our focus here was on the variety of available dMRI measures, calculated using different protocols. With this in mind, the optimal dMRI indices to include in a multimodal study may be those that contribute the greatest independent information beyond that available from anatomical MRI and other standard imaging modalities. Multivariate methods - such as machine learning (Zhou et al., 2017; Wang et al., 2018) and even deep learning (Liu et al., 2017) - may also help to extract and capitalize on features that predict clinical decline beyond those studied here.

In addition to providing a roadmap for the new ADNI3 dMRI data, these preliminary analyses show that despite differences in the updated dMRI protocols, diffusion indices can be pooled to detect white matter microstructural differences associated with aging and Alzheimer’s disease.

## Supporting information

## Author contributions Statement

RIR, MAB, BB, CRJ, PT and MWW designed the ADNI3 diffusion MRI study. ST, AZP, and TMN performed the image analysis. TMN, AZP, NJ and PT conceived and designed the image analysis study. TMN, AZP, ST, NJ and PT drafted the manuscript. All authors contributed to interpreting the results and critically revised the manuscript for intellectual content.

## Acknowledgements

This study builds on preliminary findings in a conference paper entitled, *Ranking Diffusion Tensor Measures of Brain Aging & Alzheimer’s Disease,* which may be found in the conference proceedings from the 14th International Symposium on Medical Information Processing and Analysis (SIPAIM; Zavaliangos-Petropulu et al., 2018).

