## Supplementary material for "Diffusion MRI Indices and their Relation to Cognitive Impairment in Brain Aging: The updated multi-protocol approach in ADNI3"

Paul M. Thompson, for the Alzheimer’s Disease Neuroimaging Initiative (ADNI)

*Denotes equal contribution

**1.	Supplementary Data**

**1.1	ADNI3 Excluded Participants**

**Supplementary Table 1.** Of the 381 raw dMRI scans available for download from the ADNI database as of April 2018 (<https://ida.loni.usc.edu/>), 64 were excluded from the study for the reasons summarized below. Participants with missing diagnosis, age, or with no available cognitive scores were excluded. Images were manually inspected for quality, and were removed if there were scanner-related artifacts or pathology inconsistent with the ADNI inclusion criteria, or if the data available from the IDA was corrupted. Protocols that included participants with overlapping exclusion criteria are marked with an asterisk.

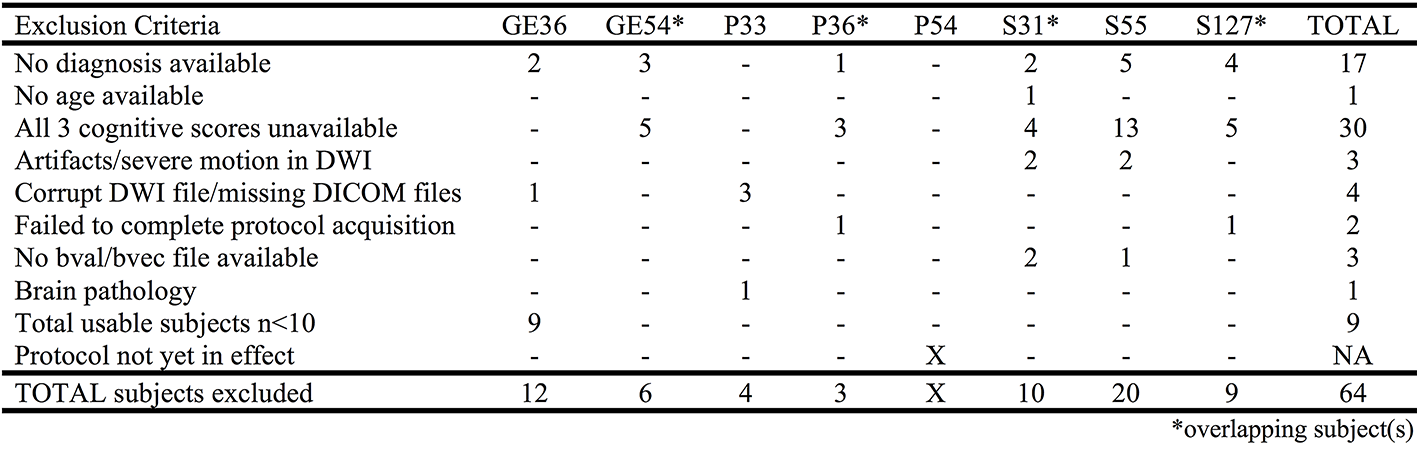

**1.2	Available Cognitive Evaluations**

**Supplementary Table 2.** Some ADNI3 participants only had a subset of cognitive assessments. Here, we break down the number of participants that have available scores for each cognitive evaluation.

| Protocol | MMSE | CDR-sob | ADAS-cog |
| --- | --- | --- | --- |
| GE54 | 64 | 65 | 61 |
| P33 | 24 | 24 | 20 |
| P36 | 19 | 19 | 18 |
| S31 | 35 | 36 | 27 |
| S55 | 153 | 152 | 132 |
| S127 | 20 | 20 | 20 |
| TOTAL | 315 | 316 | 278 |

**1.3	Demographic Data for Cognitively Normal Controls**

**Supplementary Table 3.** Demographics for ADNI2 and ADNI3 cognitively normal (CN) participants, by protocol.

| Protocol | All | | | Matched | | |
| --- | --- | --- | --- | --- | --- | --- |
|  | Total  N | Age  Mean ± SD | Male  N (%) | Total  N | Age  Mean ± SD | Male  N (%) |
| GE54 | 45 | 75.8±7.0 | 20 (44.4) | 12 | 76.4±3.8 | 4 (33.3) |
| P33 | 17 | 78.5±7.3 | 9 (52.9) | 12 | 77.0±6.3 | 4 (33.3) |
| P36 | 12 | 75.9±6.4 | 4 (33.3) | 12 | 75.9±6.4 | 4 (33.3) |
| S31 | 21 | 71.6±7.0 | 8 (38.1) | 12 | 75.3±6.9 | 4 (33.3) |
| S55 | 96 | 73.5±7.6 | 36 (37.5) | 12 | 75.6±3.5 | 4 (33.3) |
| S127 | 16 | 75.2±6.0 | 6 (37.5) | 12 | 76.5±4.8 | 4 (33.3) |
| ADNI3 | 207 | 74.5±7.4 | 83 (40.1) | 72 | 76.1±5.3 | 24 (33.3) |
| ADNI2 | 59 | 72.4±6.6 | 24 (40.7) | 12 | 76.5±3.8 | 4 (33.3) |
| Total | 266 | 74.0±7.3 | 107 (40.2) | 84 | 76.2±5.1 | 4 (33.3) |

**2.	ADNI2 and ADNI3 dMRI Associations with Age in Cognitively Normal Controls**

**2.1	ADNI2 and ADNI3 *Pooled* dMRI ROI Associations with Age**

**Supplementary Table 4.** *P*-values and corresponding effect sizes (*d*-values) are reported for associations between age and 5 dMRI anisotropy and diffusivity indices in pooled ADNI2 and ADNI3 control participants (N=266). For each test, the ROIs are ordered by *d*-value. Regions that were significant after FDR (*q* = 0.05) or Bonferroni (α = 0.05) multiple comparisons correction are delineated by a dotted or solid line respectively.

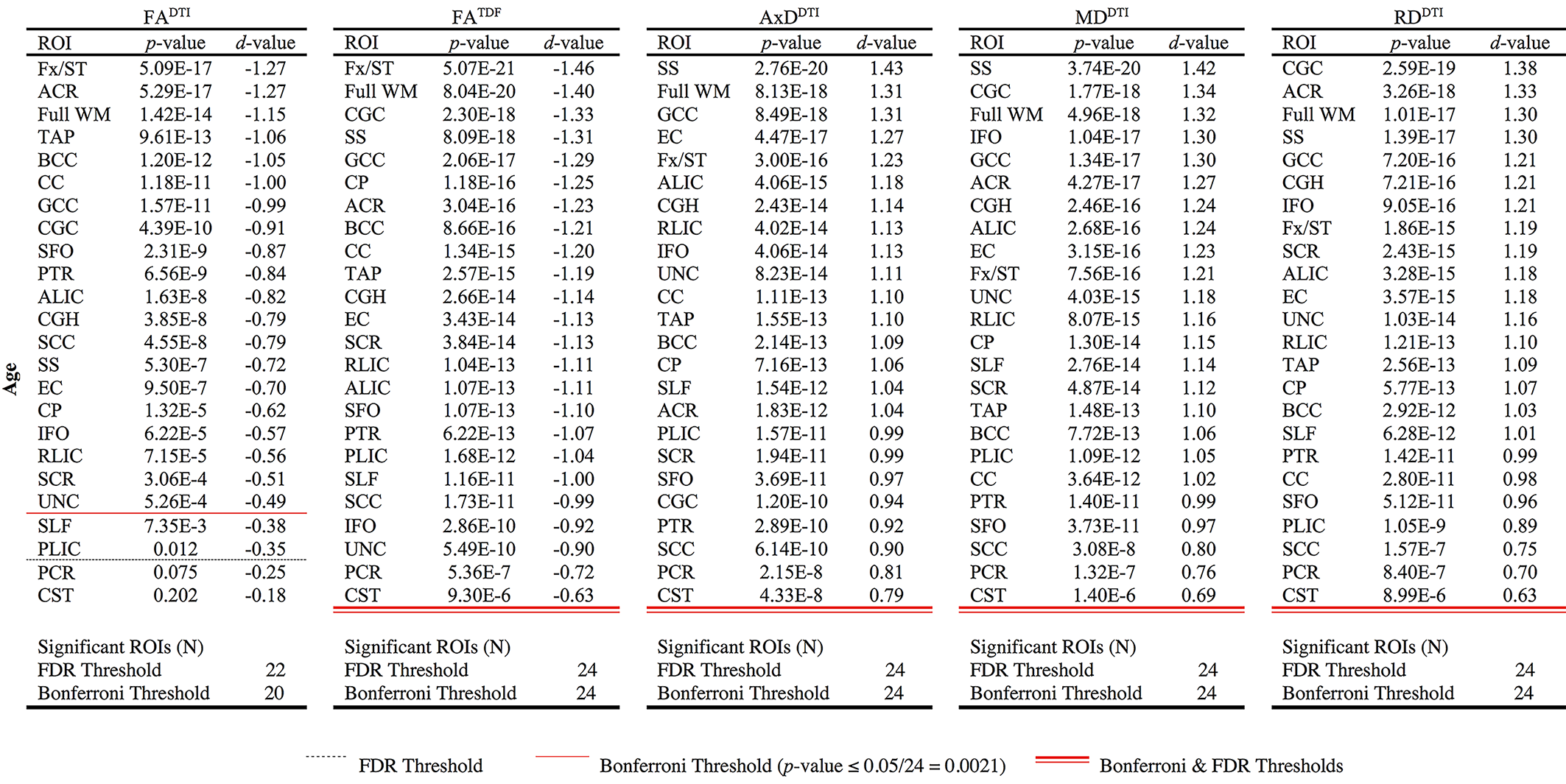

**2.2	dMRI ROI Associations with Age *by Protocol***

**2.2.1	GE54 Associations with Age**

**Supplementary Table 5.** For CN participants scanned with protocol GE54 (N=45), *p*-values and corresponding effect sizes (*d*-values) are reported for associations between age and 5 dMRI anisotropy and diffusivity indices. For each test, the ROIs are ordered by *d*-value. Regions that were significant after FDR (*q* = 0.05) or Bonferroni (α = 0.05) multiple comparisons correction are delineated by a dotted or solid line respectively.

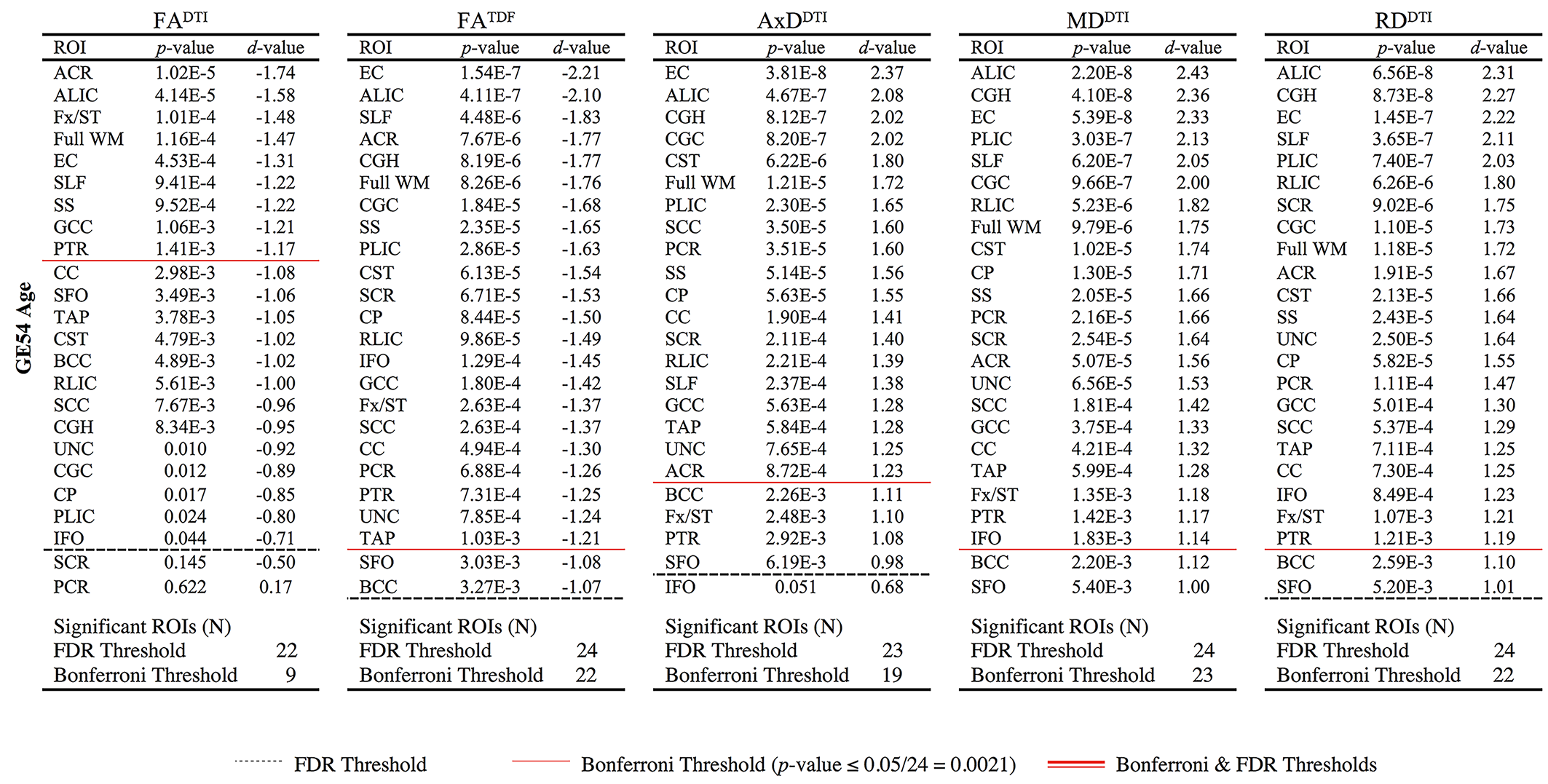

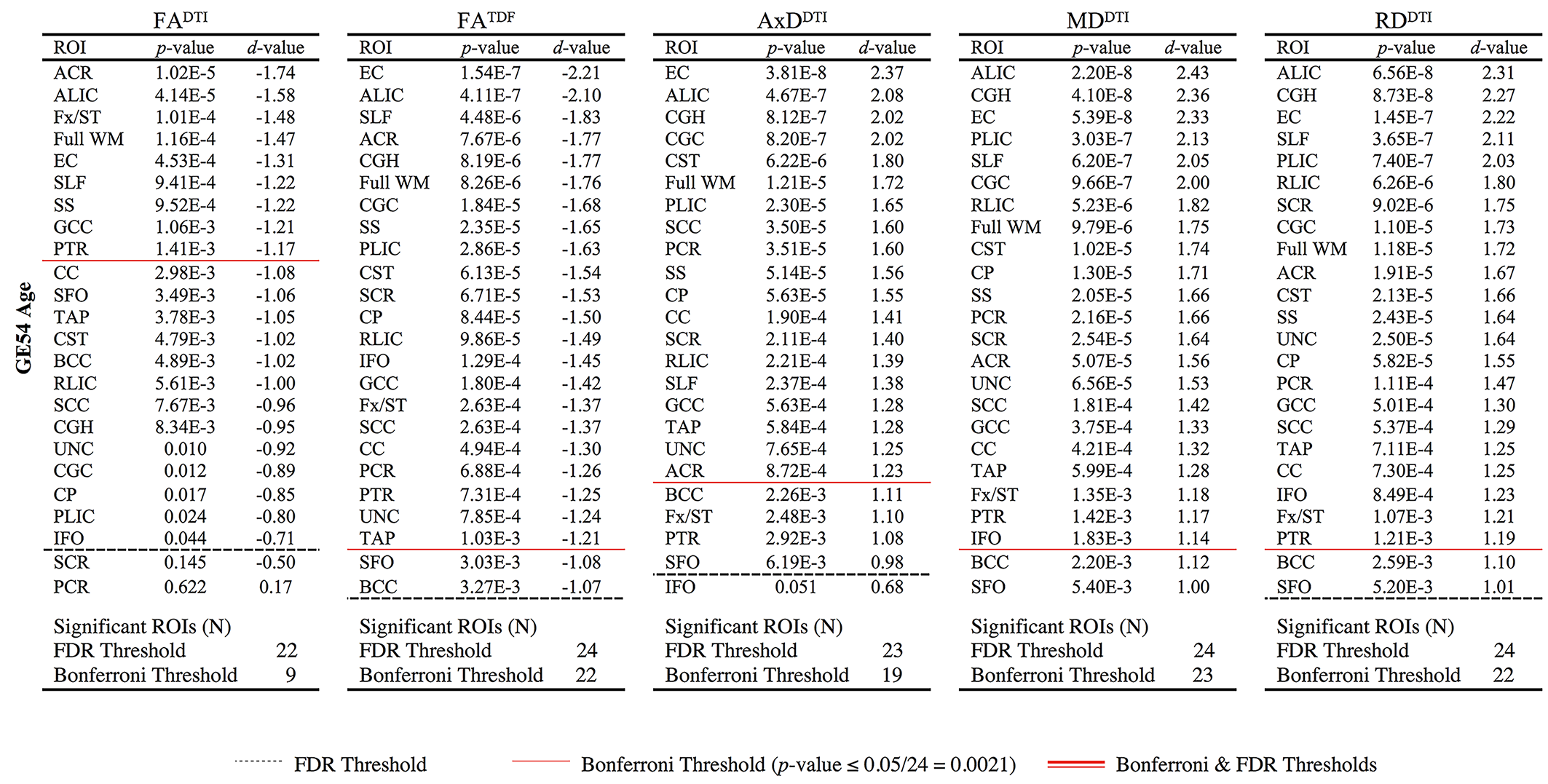

**2.2.2	P33 Associations with Age**

**Supplementary Table 6.** For CN participants scanned with protocol P33 (N=17), *p*-values and corresponding effect sizes (*d*-values) are reported for associations between age and 5 dMRI anisotropy and diffusivity indices. For each test, the ROIs are ordered by *d*-value. Regions that were significant after FDR (*q* = 0.05) or Bonferroni (α = 0.05) multiple comparisons correction are delineated by a dotted or solid line respectively.

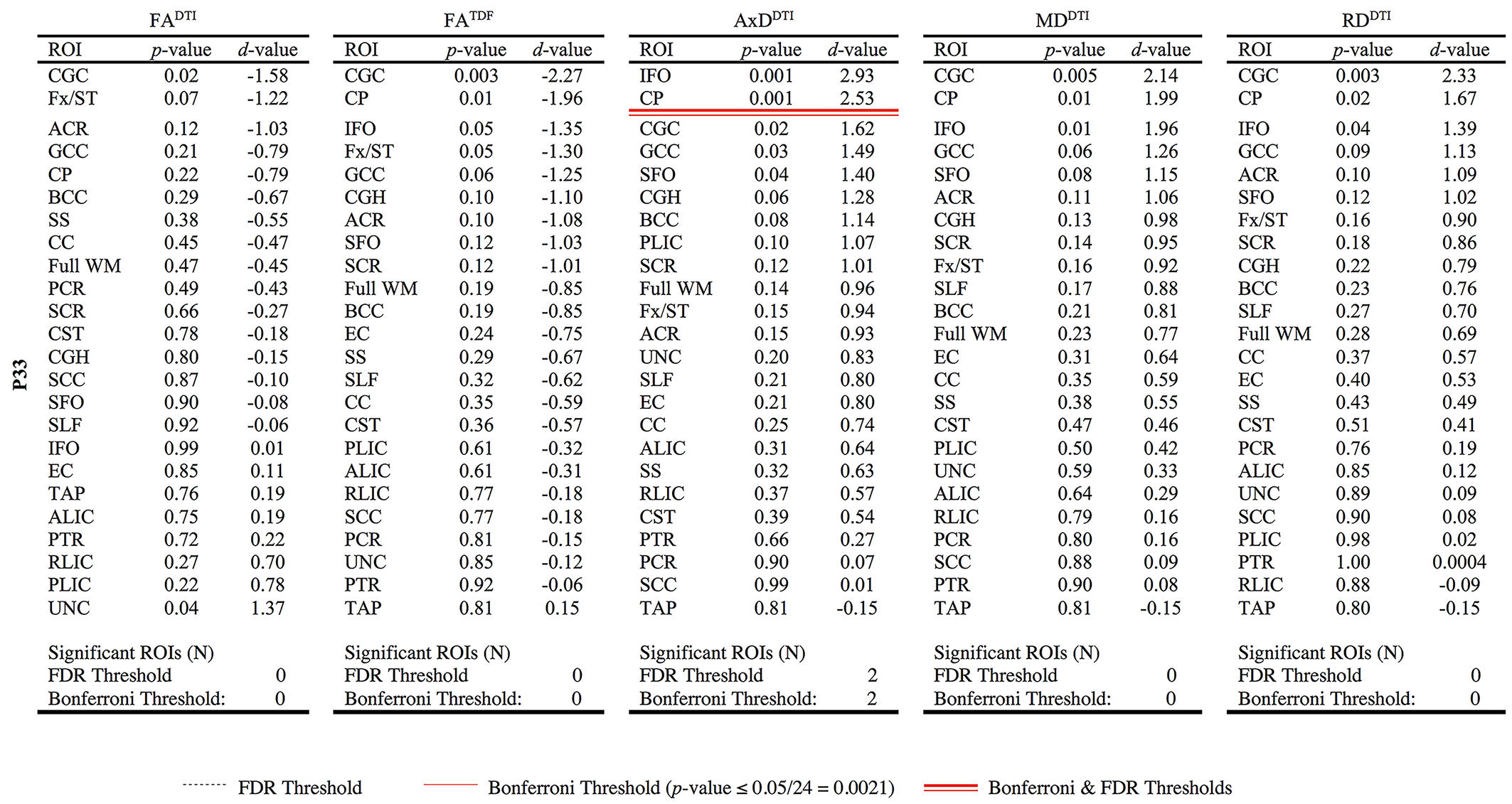

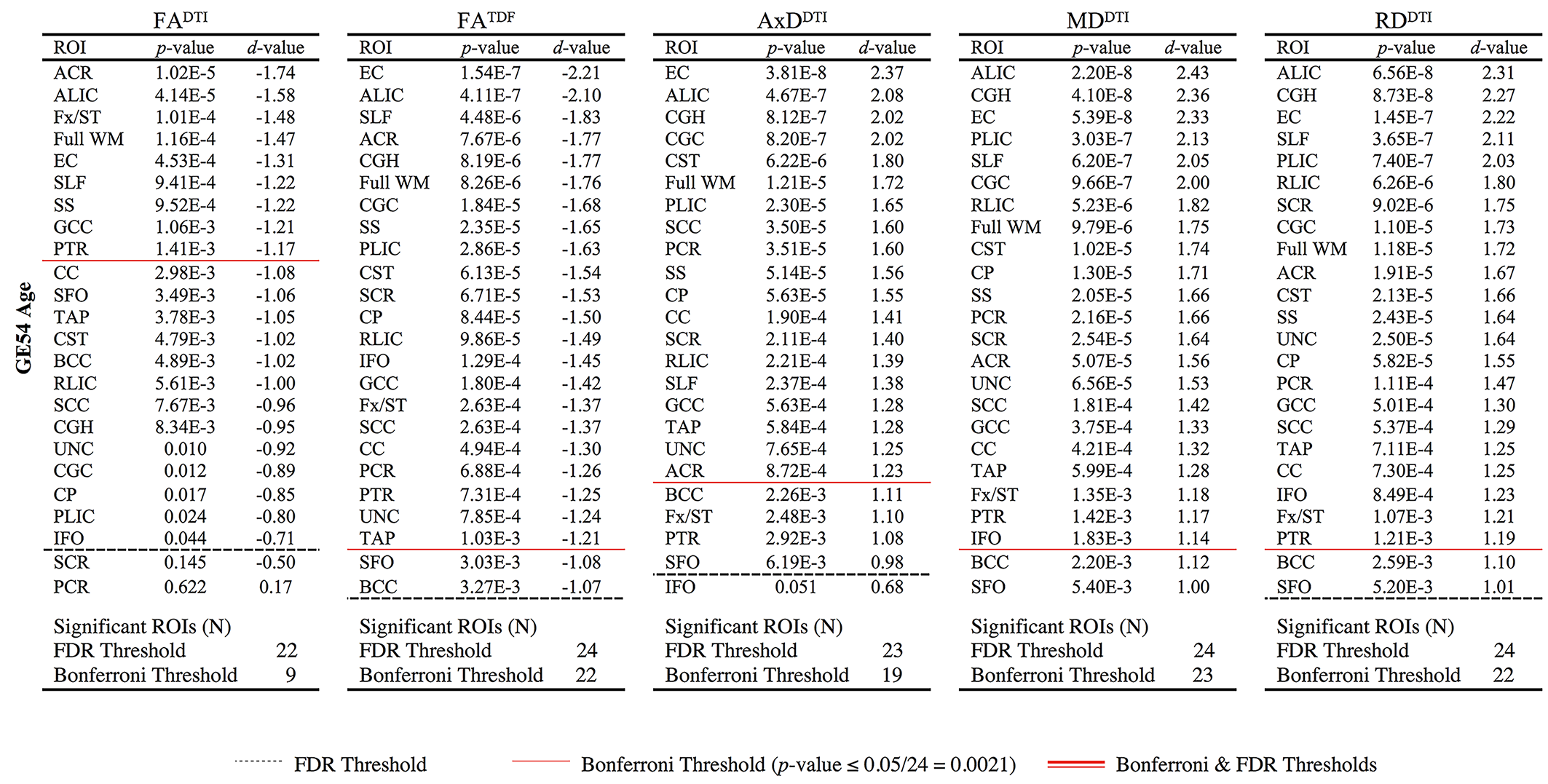

**2.2.3	P36 Associations with Age**

**Supplementary Table 7.** For CN participants scanned with protocol P36 (N=12), *p*-values and corresponding effect sizes (*d*-values) are reported for associations between age and 5 dMRI anisotropy and diffusivity indices. For each test, the ROIs are ordered by *d*-value. No associations were significant after correction for multiple comparisons.

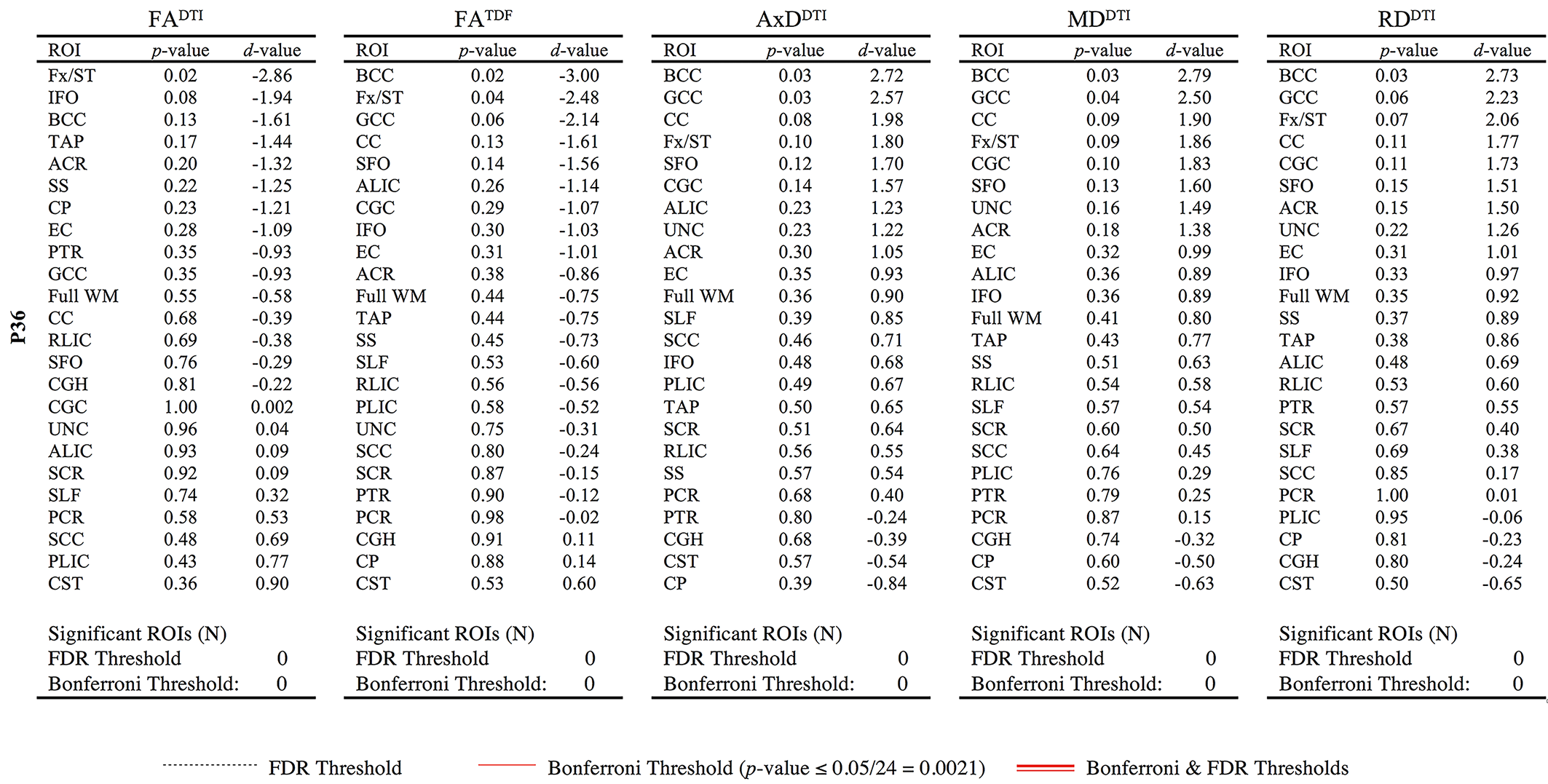

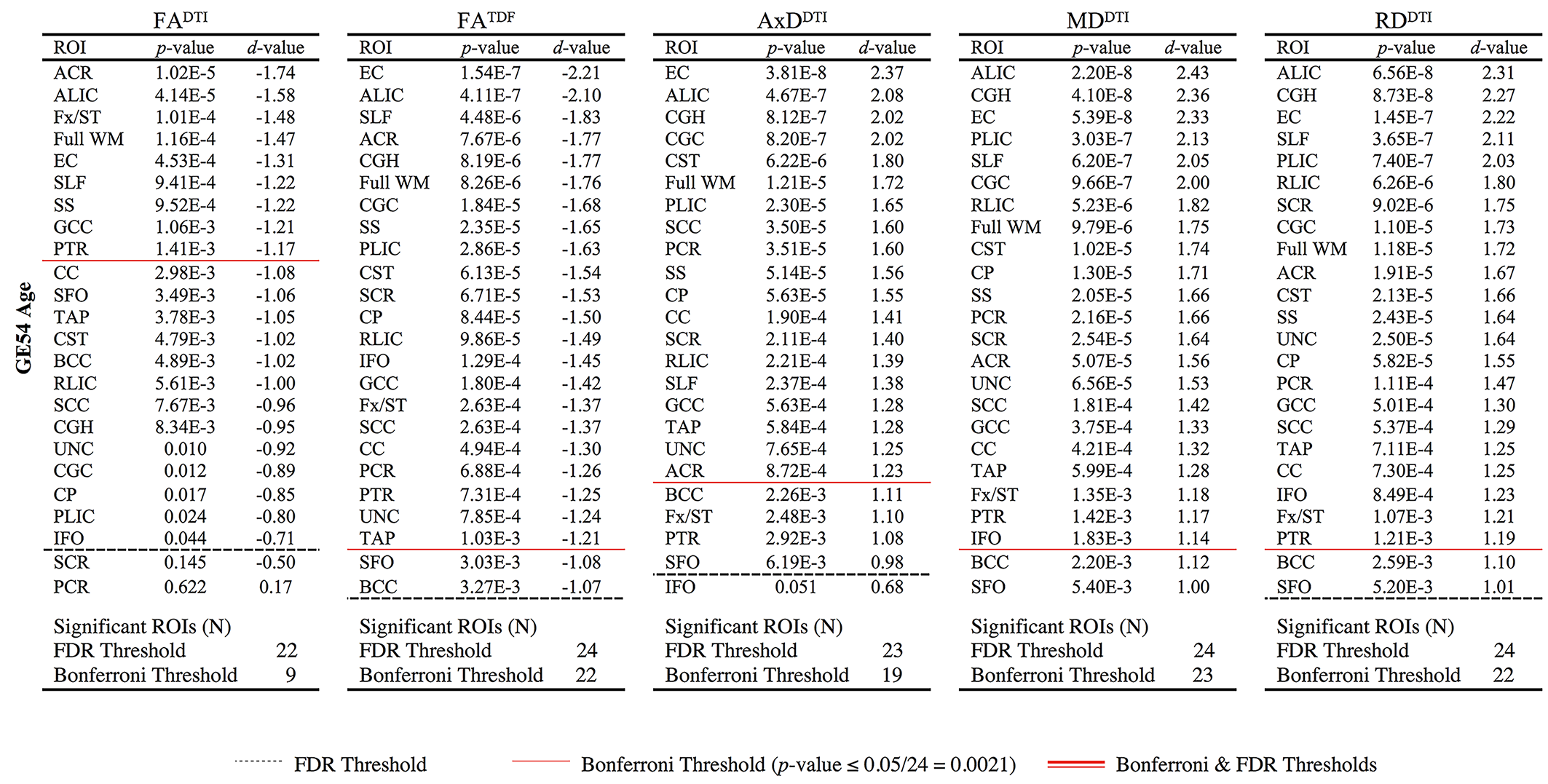

**2.2.4	S31 Associations with Age**

**Supplementary Table 8.** For CN participants scanned with protocol S31 (N=21), *p*-values and corresponding effect sizes (*d*-values) are reported for associations between age and 5 dMRI anisotropy and diffusivity indices. For each test, the ROIs are ordered by *d*-value. No associations were significant after correction for multiple comparisons.

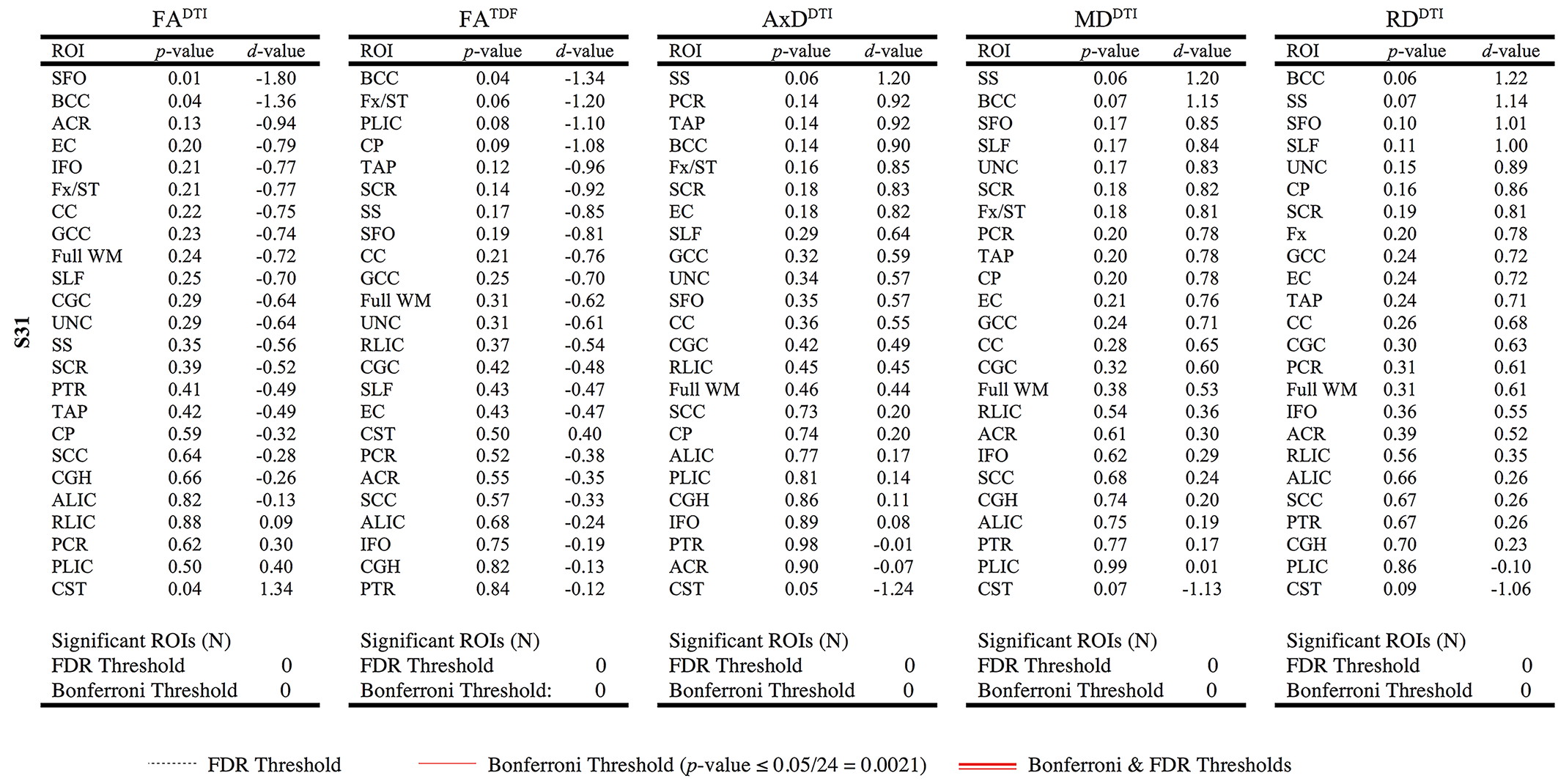

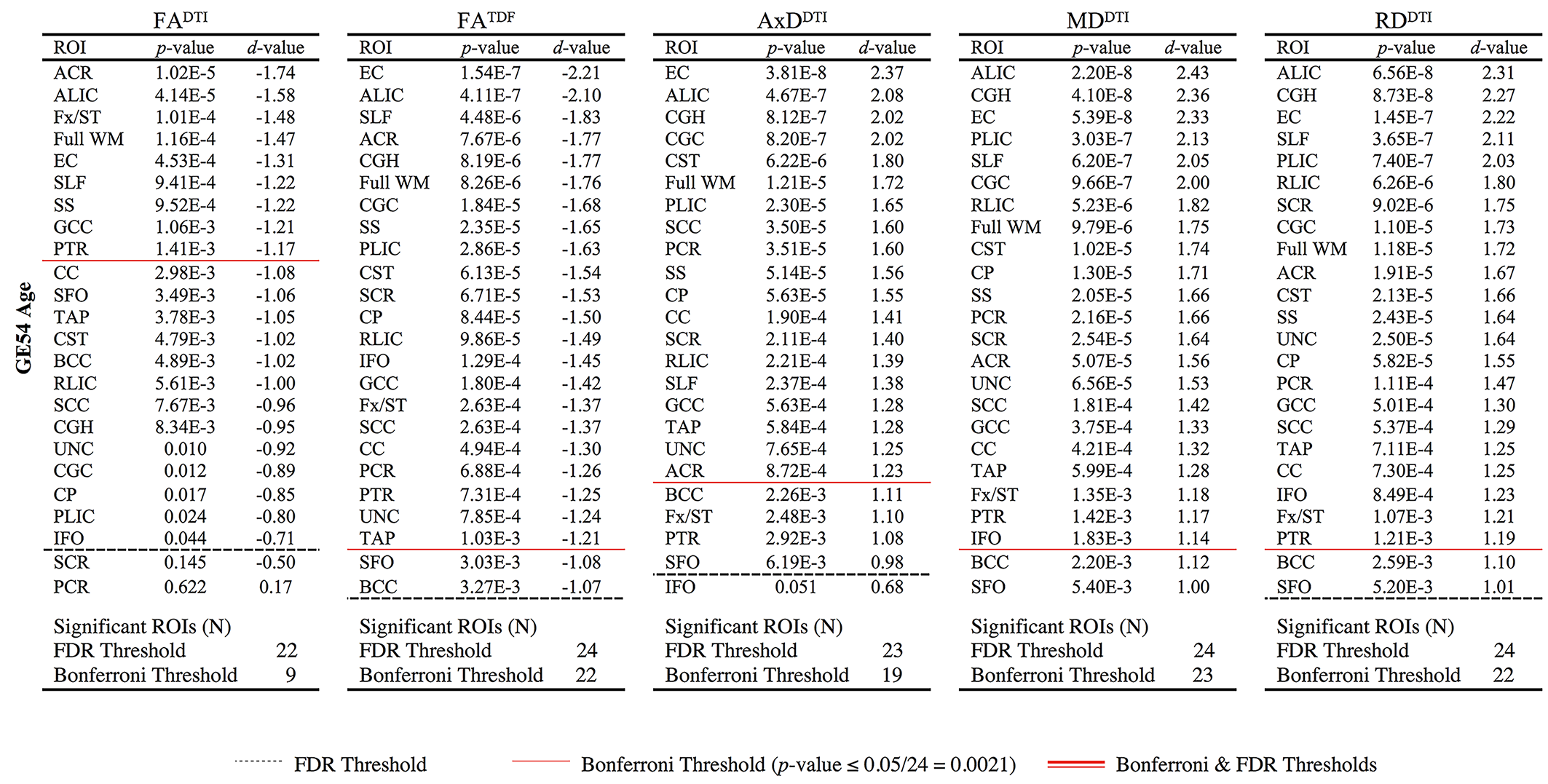

**2.2.5	S55 Associations with Age**

**Supplementary Table 9.** For CN participants scanned with protocol S55 (N=96), *p*-values and corresponding effect sizes (*d*-values) are reported for associations between age and 5 dMRI anisotropy and diffusivity indices. For each test, the ROIs are ordered by *d*-value. Regions that were significant after FDR (*q* = 0.05) or Bonferroni (α = 0.05) multiple comparisons correction are delineated by a dotted or solid line respectively.

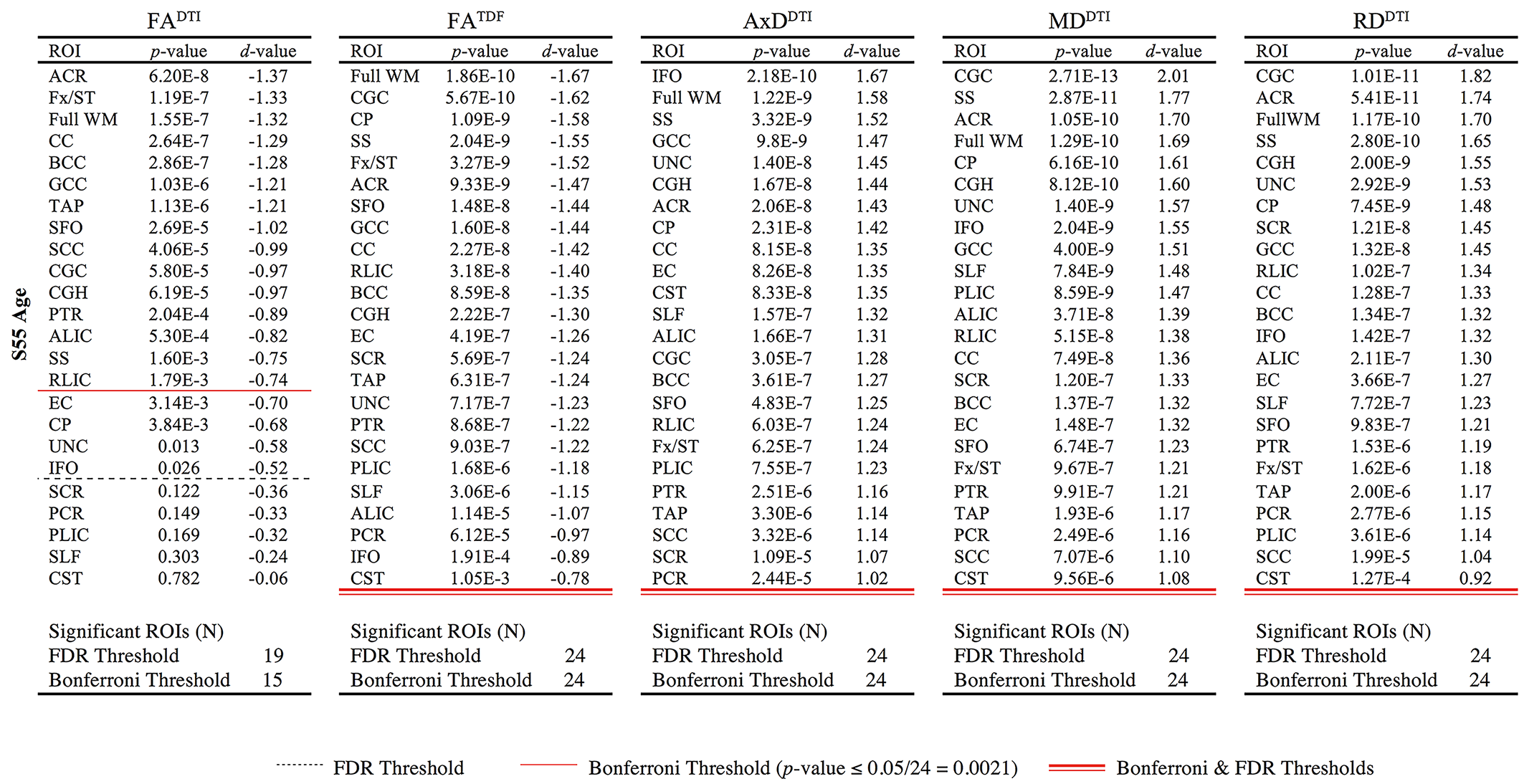

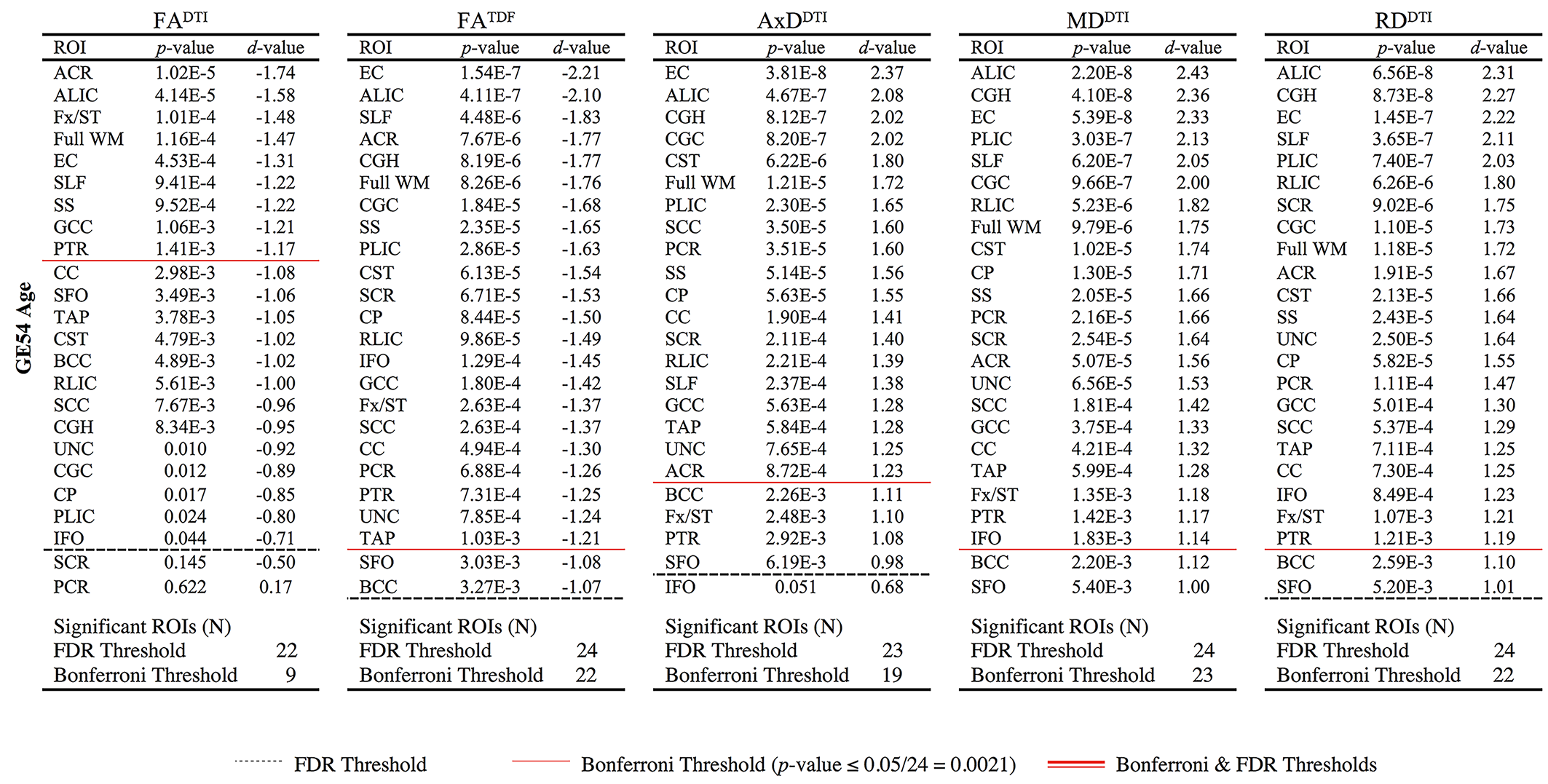

**2.2.6	S127 Associations with Age**

**Supplementary Table 10.** For CN participants scanned with protocol S127 (N=16), *p*-values and corresponding effect sizes (*d*-values) are reported for associations between age and 5 dMRI anisotropy and diffusivity indices. For each test, the ROIs are ordered by *d*-value. Regions that were significant after FDR (*q* = 0.05) or Bonferroni (α = 0.05) multiple comparisons correction are delineated by a dotted or solid line respectively.

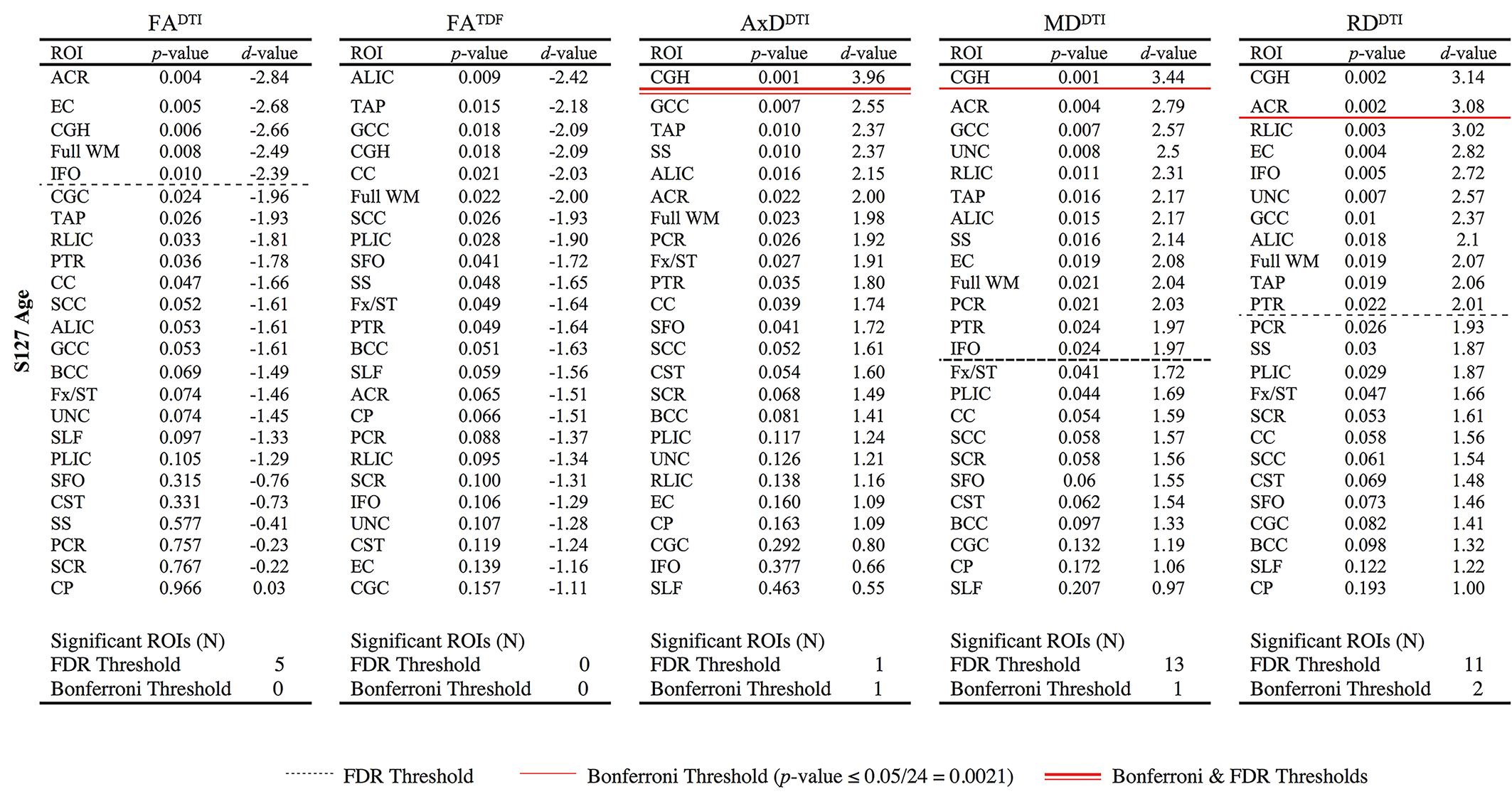

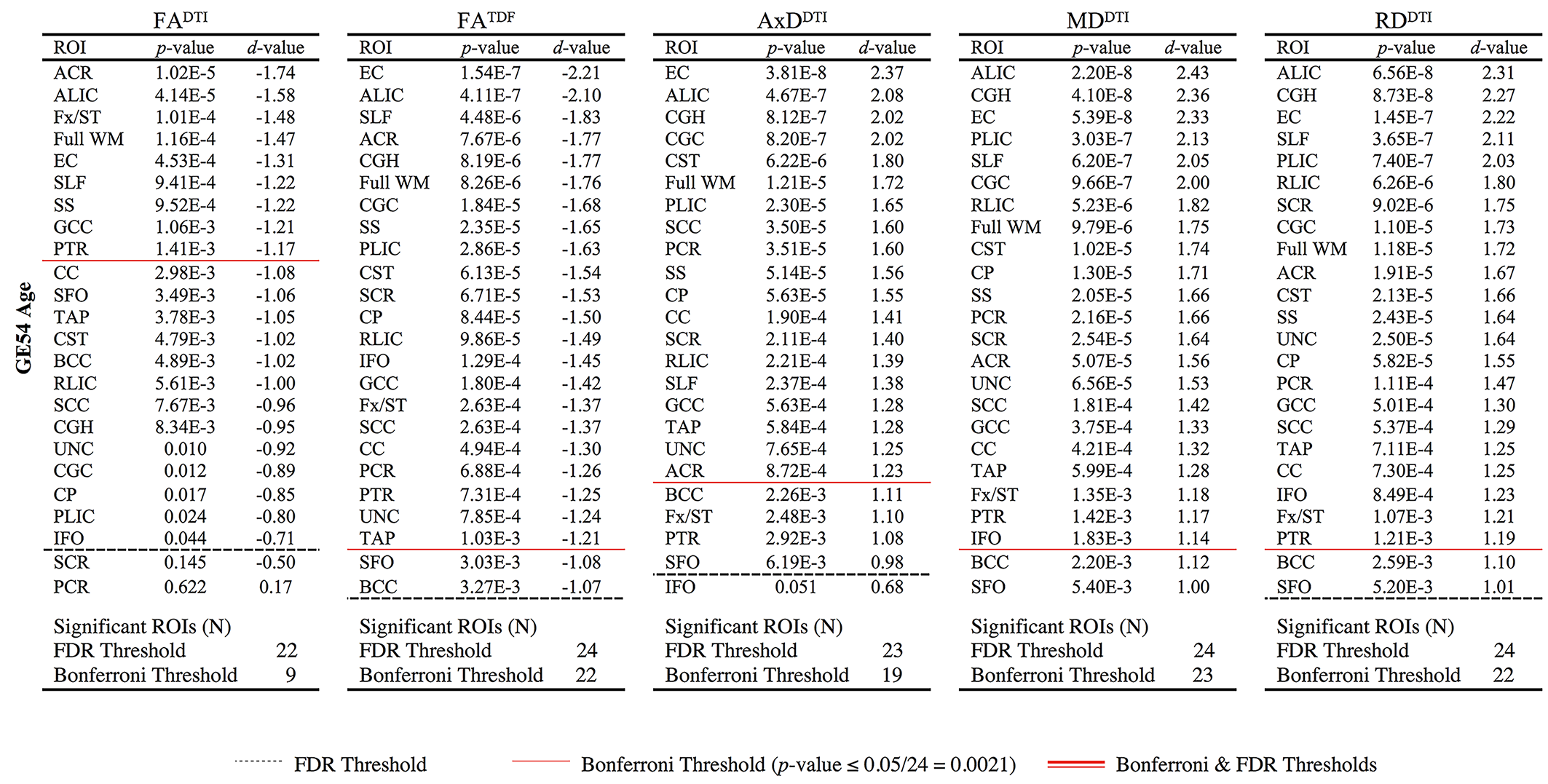

**2.2.7	ADNI2 Associations with Age**

**Supplementary Table 11.** For CN participants scanned with the ADNI2 protocol (N=59), *p*-values and corresponding effect sizes (*d*-values) are reported for associations between age and 5 dMRI anisotropy and diffusivity indices. For each test, the ROIs are ordered by *d*-value. Regions that were significant after FDR (*q* = 0.05) or Bonferroni (α = 0.05) multiple comparisons correction are delineated by a dotted or solid line respectively.

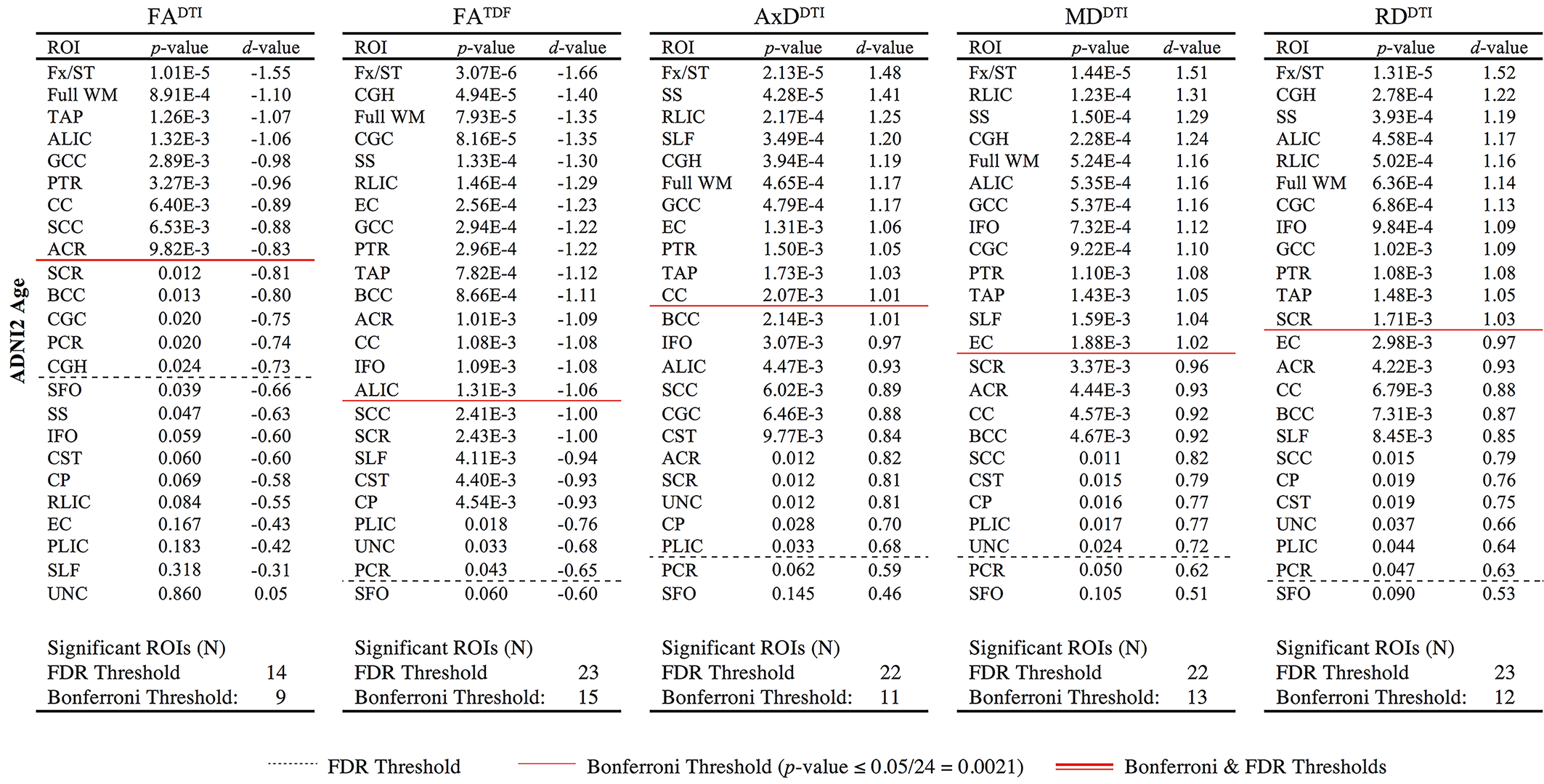

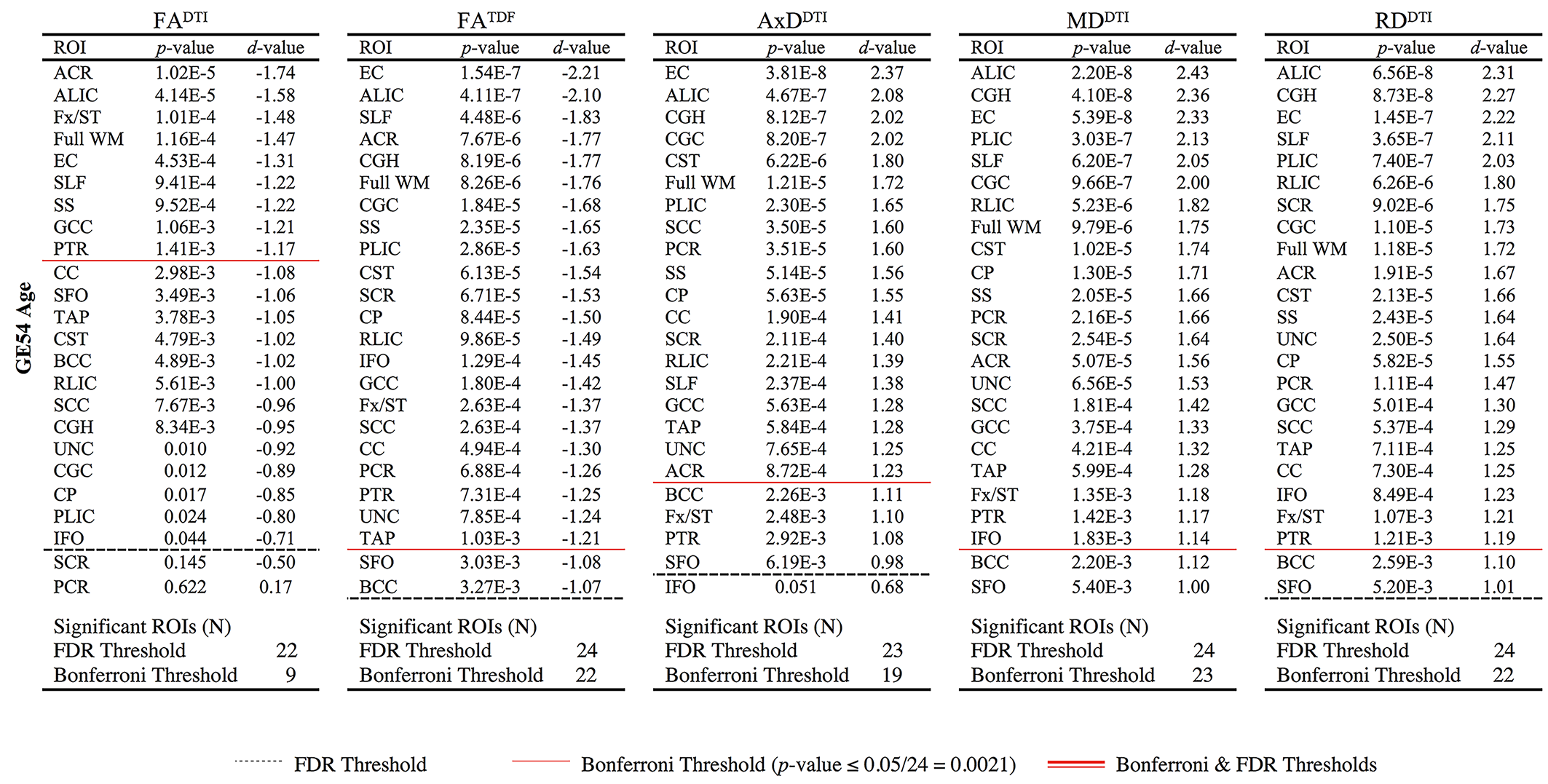

**2.3	dMRI Associations with Age *by Protocol* in Age and Sex Matched Subsets of N=12**

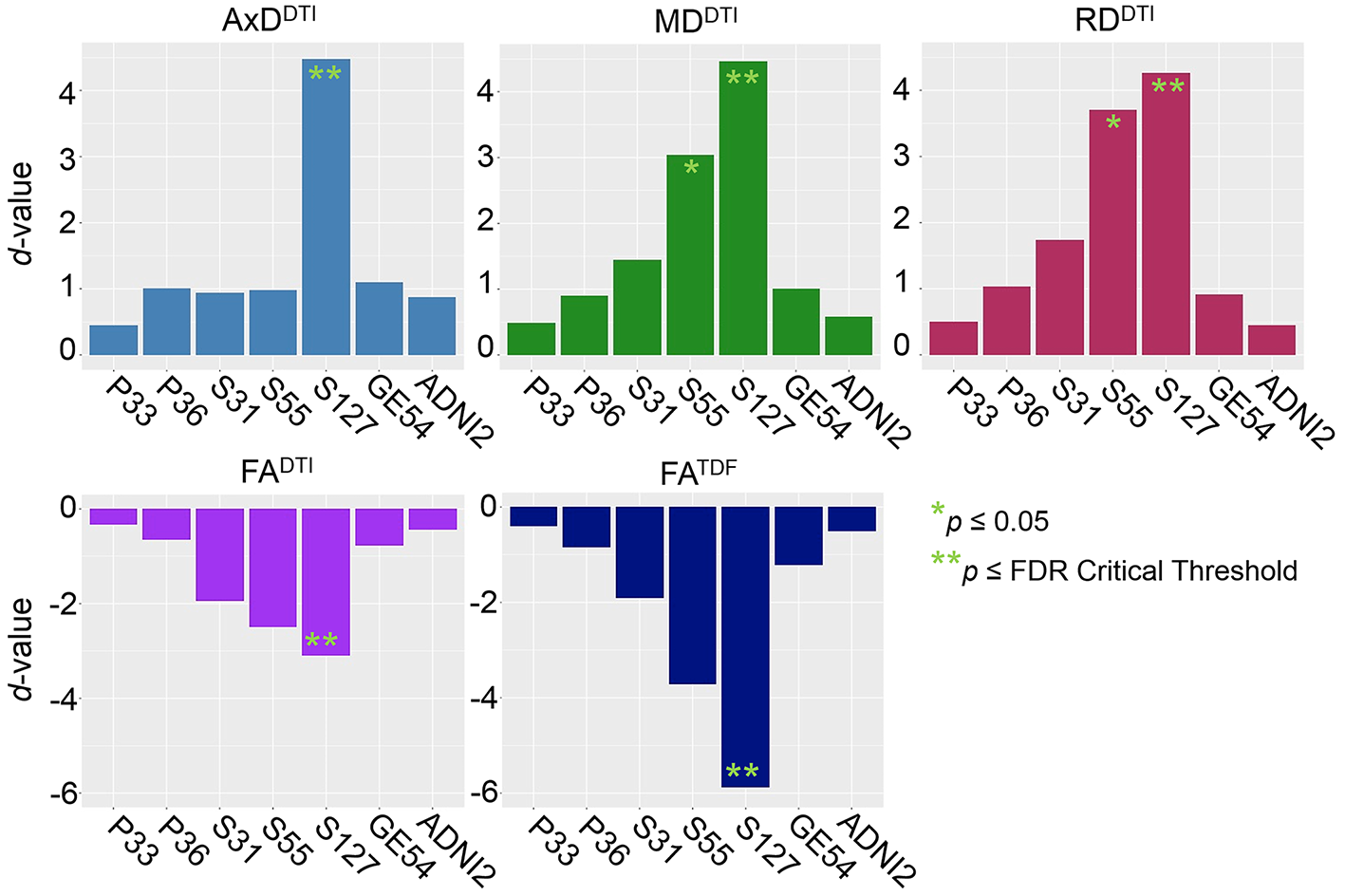

**Supplementary Figure 1.** Effect sizes (*d*-values) from 12 CN sex- and age-matched controls from each ADNI protocol (**Supplementary Table 3**) show the direction of dMRI associations with age in the full WM are consistent across protocols. While direct comparisons of effect sizes are underpowered, findings are suggestive of larger effect sizes for protocol S127, the protocol with greatest total number of diffusion-weighted (*b* = 1000 s/mm^2^) and non-diffusion sensitized (*b*_0_) gradients, followed by S55, the protocol with the second greatest number of diffusion-weighted and *b*_0_ gradients.

**3.	ComBat Harmonization of ADNI2 and ADNI3 Protocols**

**3.1	Protocol Differences in dMRI Indices After ComBat**

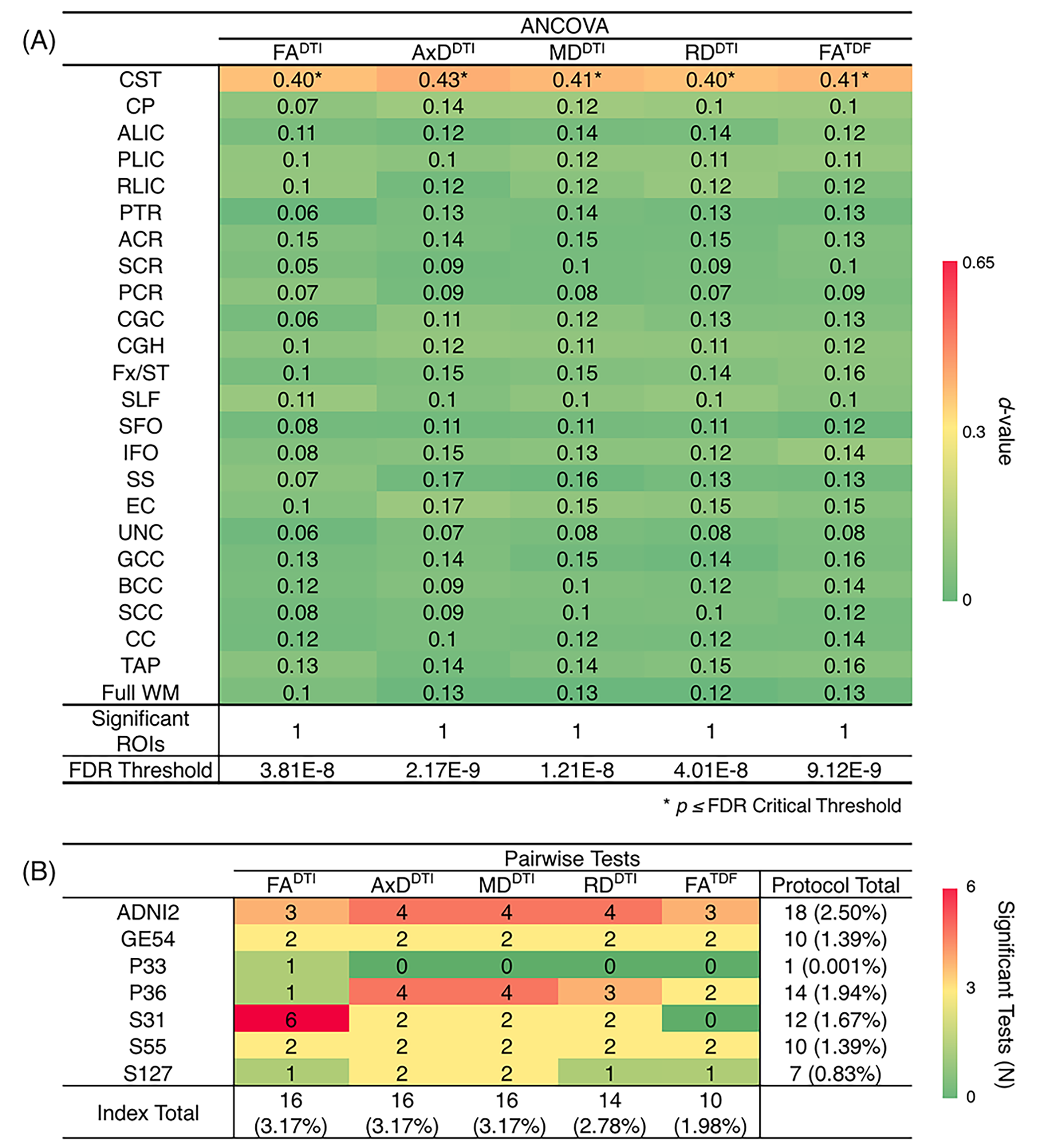

**Supplementary Figure 2. (A)** *d*-values from the ANCOVAs assessing differences in dMRI indices between protocols, for each of the 24 ROIs, after ComBat harmonization; the CST is the only region where significant protocol differences remain, across dMRI indices. **(B)** We report the number of times each protocol and each dMRI index showed significant differences in pairwise tests between protocols after ComBat harmonization (out of 504 tests per index and 720 tests per protocol). After running ComBat, the number of pairwise tests for which each protocol showed significant differences in dMRI indices decreased by 93.8%.

**3.2	ADNI2 and ADNI3 *Pooled* dMRI ROI Associations with Age after ComBat**

**3.2.1	Associations with Age after ComBat**

**Supplementary Table 12.** *P*-values and corresponding effect sizes (*d*-values) are reported for ROI associations between age and 5 dMRI anisotropy and diffusivity indices in ADNI2 and ADNI3 protocols pooled after harmonization with ComBat. For each test, the ROIs are ordered by *d*-value. Regions that were significant after FDR (*q* = 0.05) or Bonferroni (α = 0.05) multiple comparisons correction are delineated by a dotted or solid line respectively.

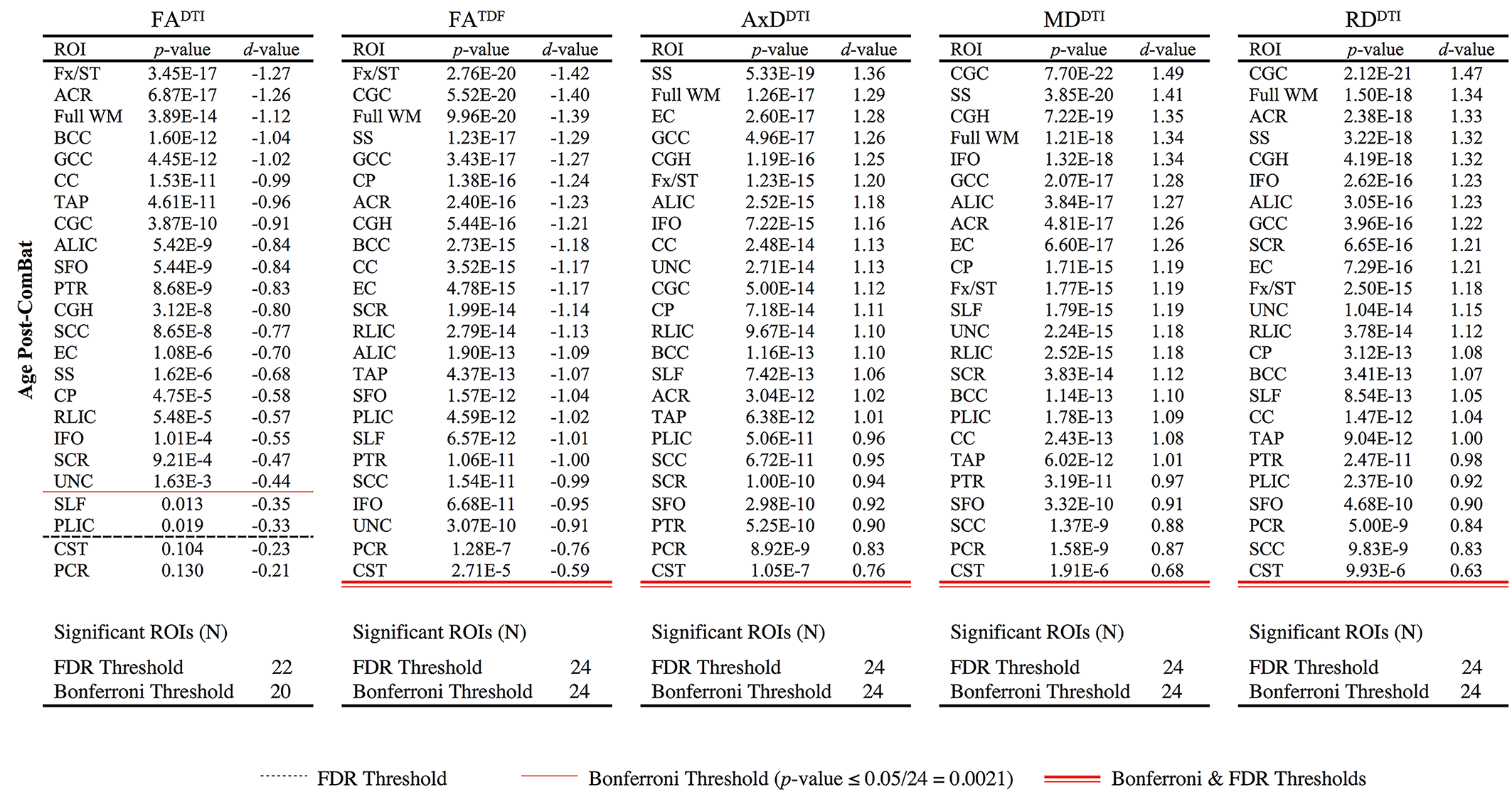

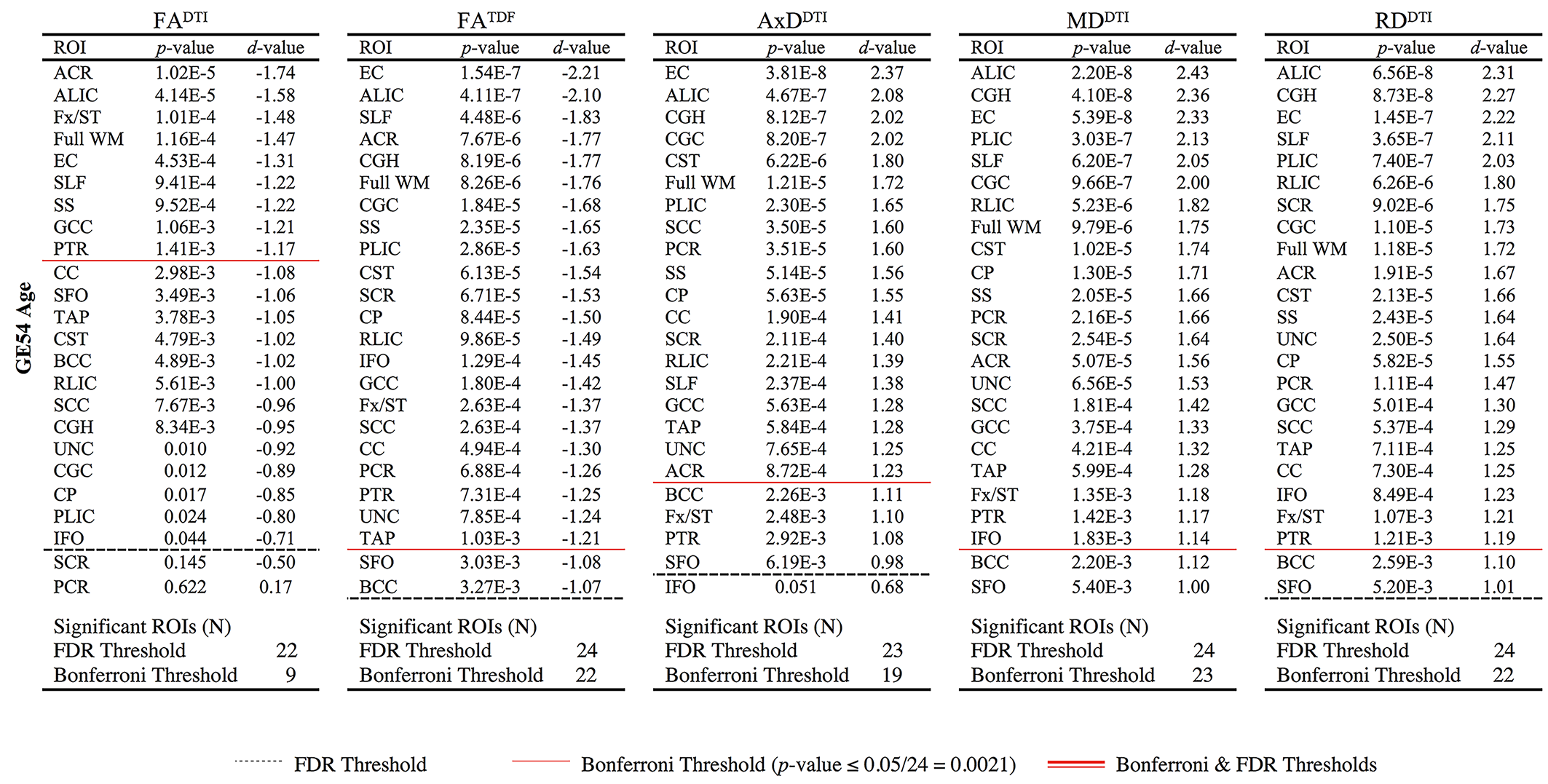

**3.2.2	dMRI Effect Sizes Before and After ComBat**

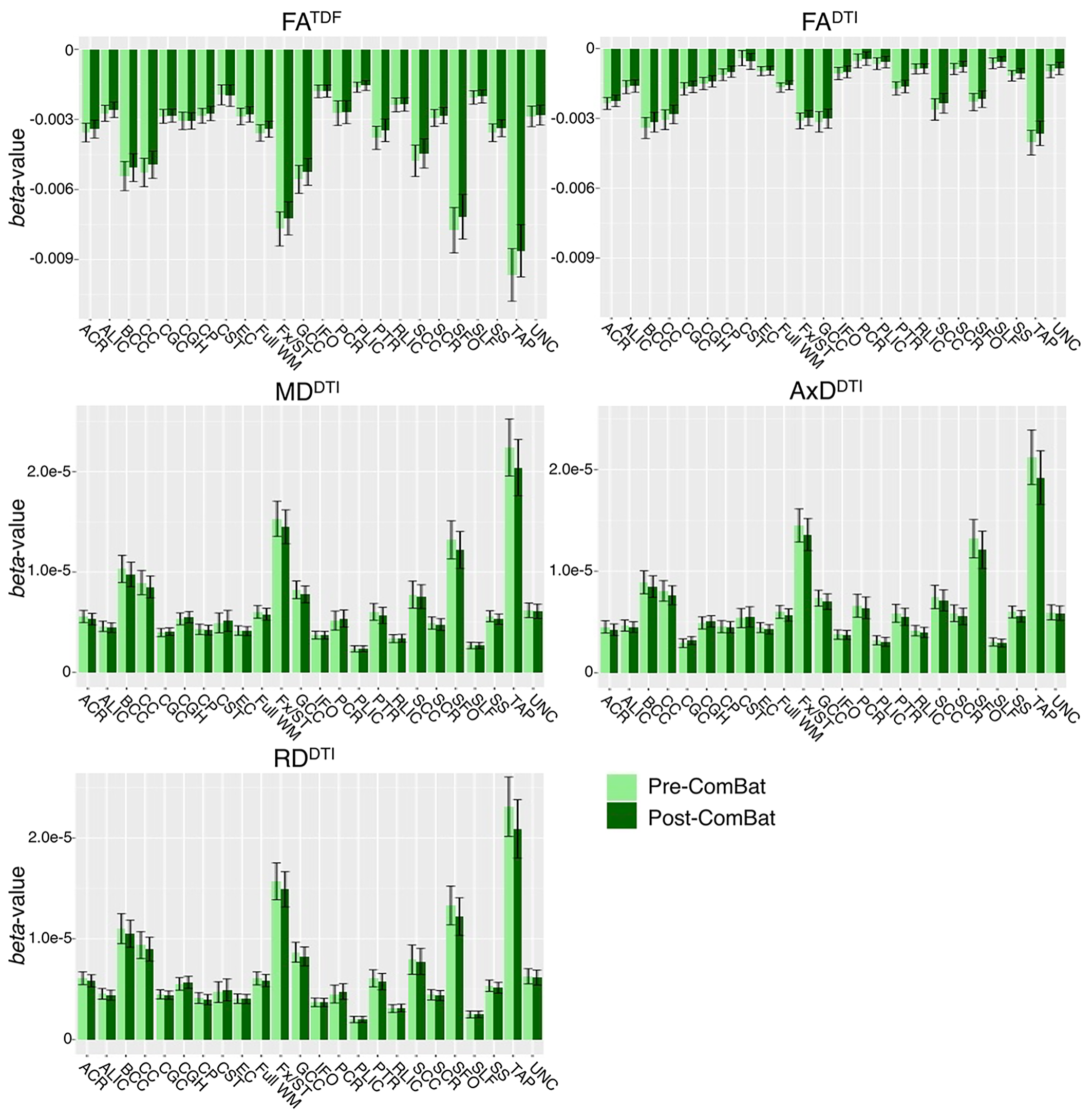

**Supplementary Figure 3.** Effect sizes (*beta*-values with error bars that represent the standard error) are plotted for the association between each diffusion index and age in elderly cognitively normal controls from ADNI2 and ADNI3 protocols pooled together before and after ComBat harmonization. Compared to pre-ComBat analyses, effect sizes are marginally different across indices, but still within the standard error bounds. All associations were significant (FDR *q* = 0.05) except for FA^DTI^ in the CST and PCR.

**3.3	dMRI ROI Associations with Age after ComBat *by Protocol***

**3.3.1	Full WM Associations with Age**

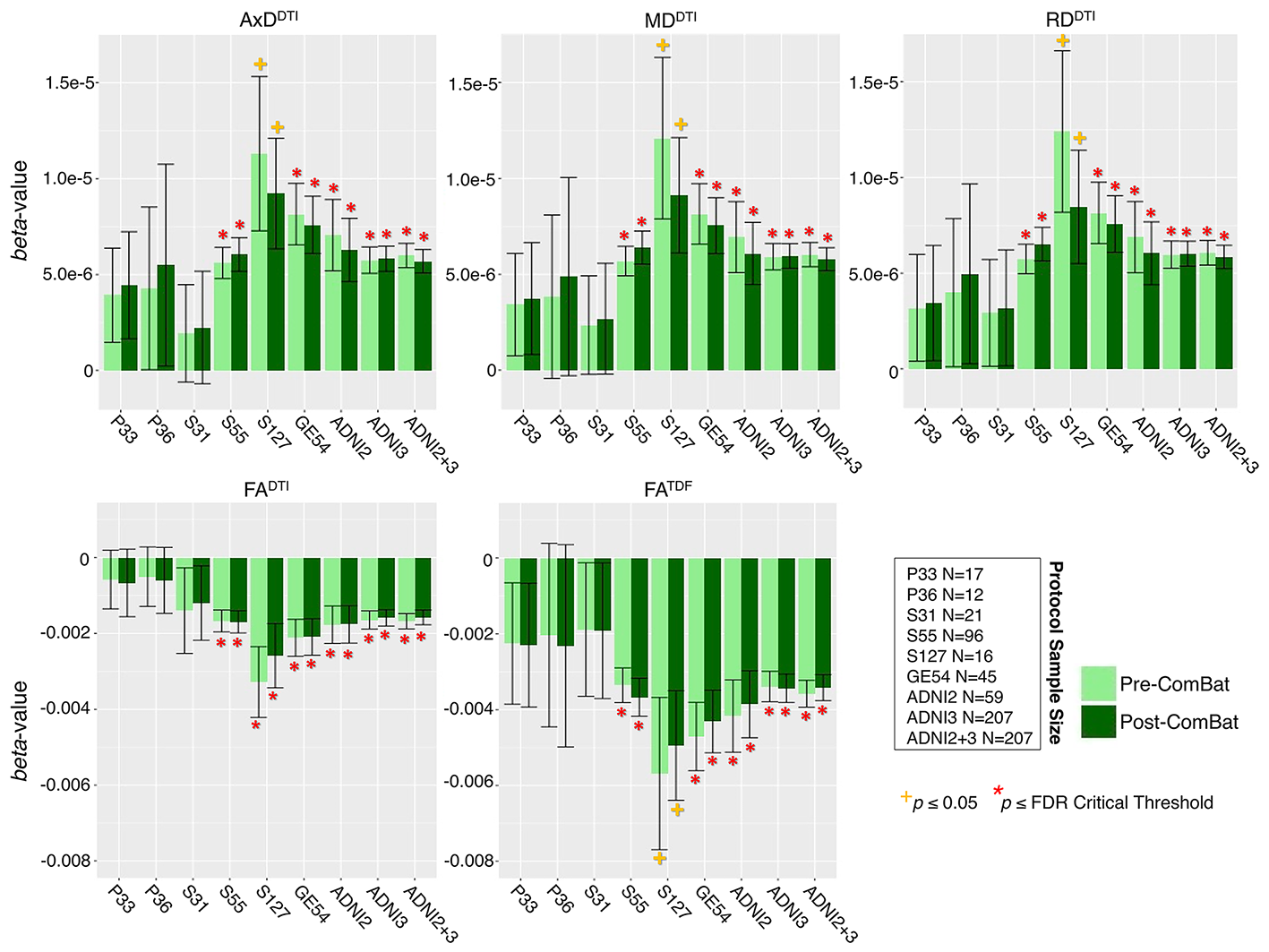

**Supplementary Figure 4.** For each protocol, the *beta*-values (error bars represent the standard error) are plotted for the association between each diffusion index in the full WM and age in elderly cognitively normal controls, before and after ComBat harmonization. Compared to pre-ComBat analyses, effect sizes are marginally different across indices, but still within the standard error bounds.

**3.3.2	Fx/ST dMRI Associations with Age**

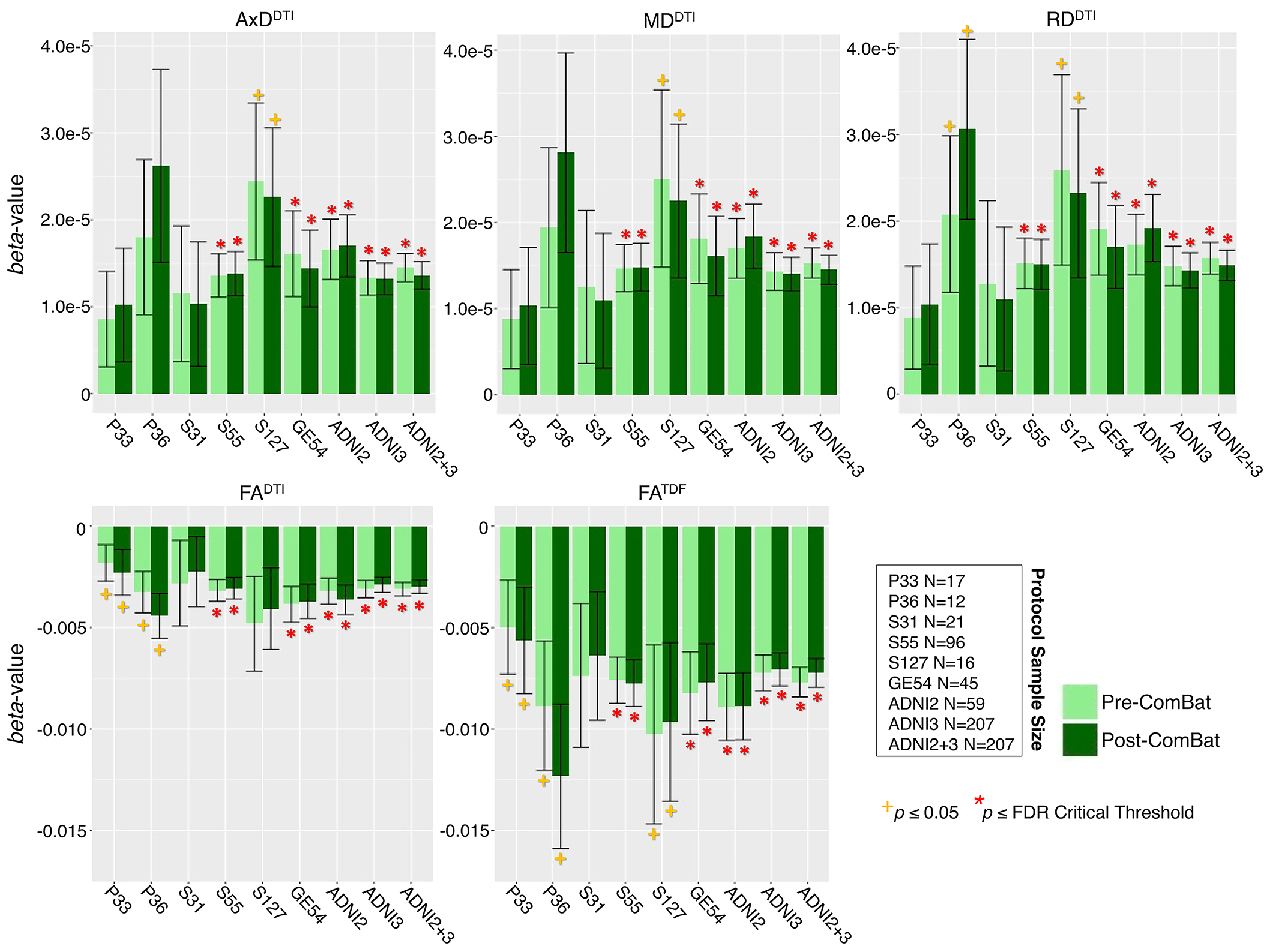

**Supplementary Figure 5.** For each protocol, the *beta*-values (error bars represent the standard error) are plotted for the association between each diffusion index in the fornix *(crus)* / *stria terminalis* (Fx/ST) and age in elderly cognitively normal controls, before and after ComBat harmonization. Compared to pre-ComBat analyses, effect sizes are marginally different across indices, but still within the standard error bounds.

**3.3.3	GCC dMRI Associations with Age**

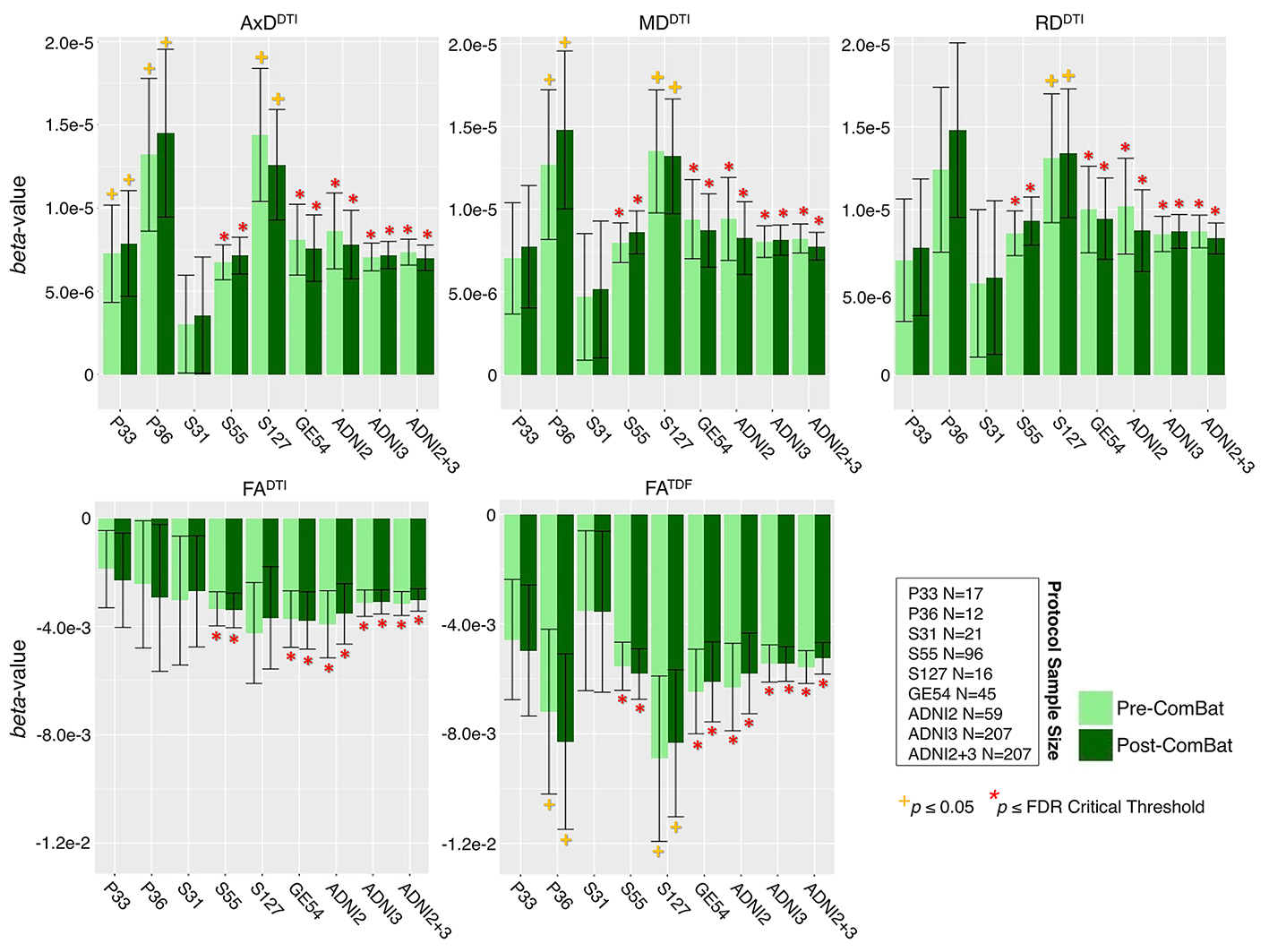

**Supplementary Figure 6.** For each protocol, the *beta*-values (error bars represent the standard error) are plotted for the association between each diffusion index in the corpus callosum genu (GCC) and age in elderly cognitively normal controls, before and after ComBat harmonization. Compared to pre-ComBat analyses, effect sizes are only marginally different across indices and protocols, but still within the standard error bounds.

**4.	ADNI3 dMRI Associations with Clinical Measures**

**4.1	Regional dMRI Measures: Associations with Cognitive Measures**

**4.1.1	CDR-sob Associations**

**Supplementary Table 13.** *P*-values and corresponding effect sizes *(d*-values) are reported for ROI associations between CDR-sob and dMRI indices across all pooled ADNI3 participants (N=316). For each test, the ROIs are ordered by *d*-value. Regions that were significant after FDR (*q*=0.05) or Bonferroni (α=0.05) multiple comparisons correction are delineated by a dotted or solid line respectively.

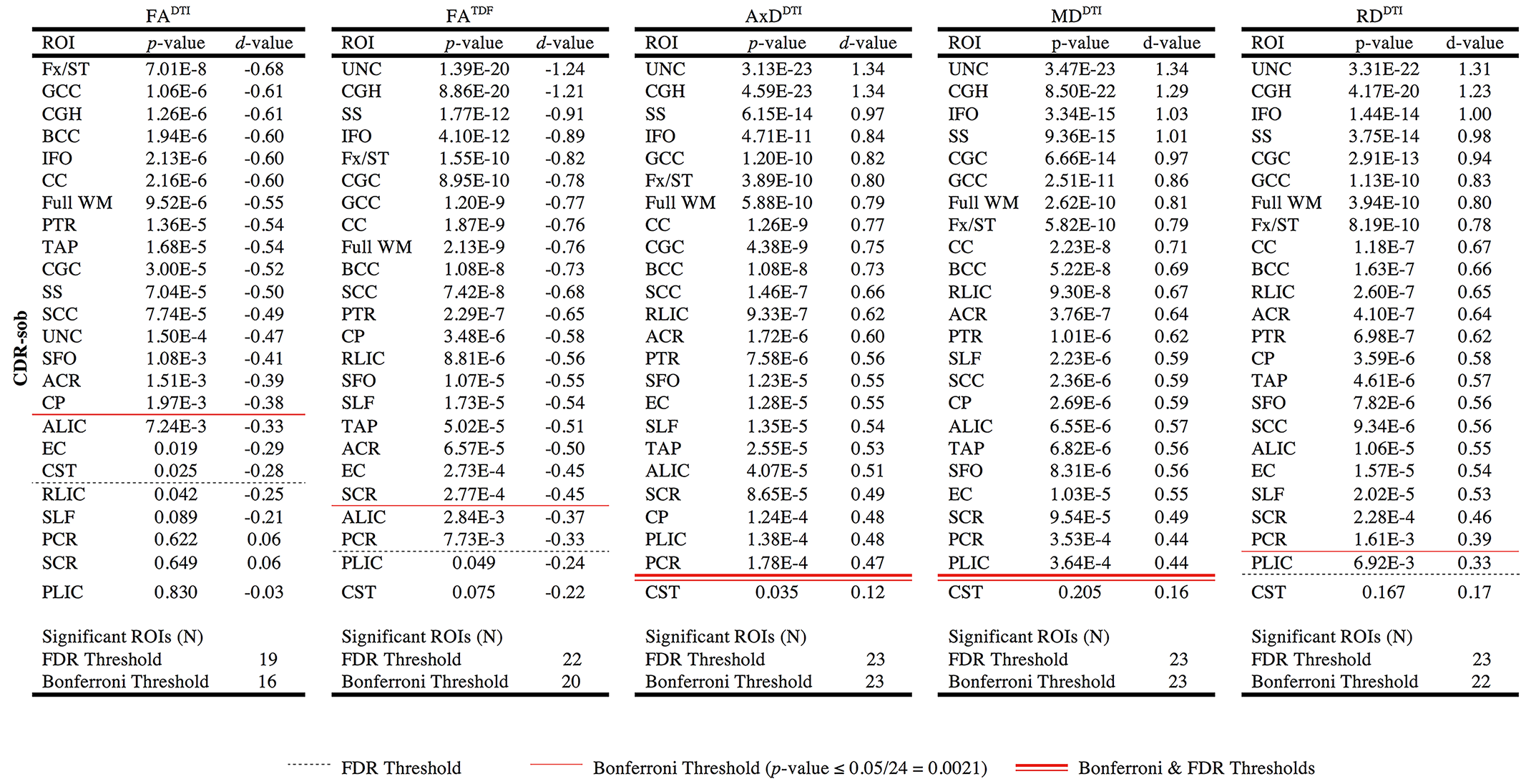

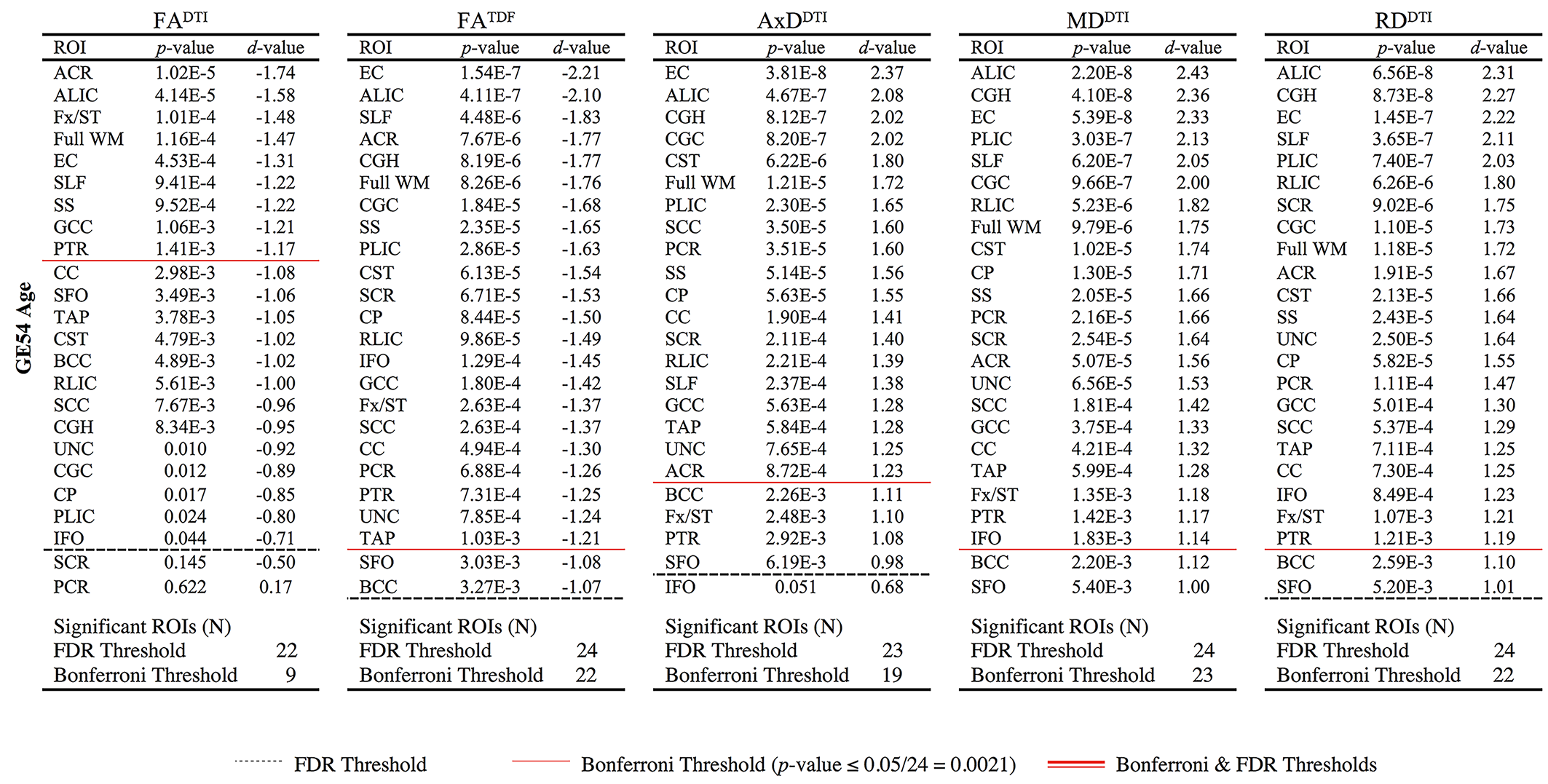

**4.1.2	ADAS-cog Associations**

**Supplementary Table 14.** *P*-values and corresponding effect sizes *(d*-values) are reported for ROI associations between ADAS-cog diagnosis and dMRI indices across all ADNI3 participants pooled together (N=278). For each test, the ROIs are ordered by *d*-value. Regions that were significant after FDR (*q*=0.05) or Bonferroni (α=0.05) multiple comparisons correction are delineated by a dotted or solid line respectively.

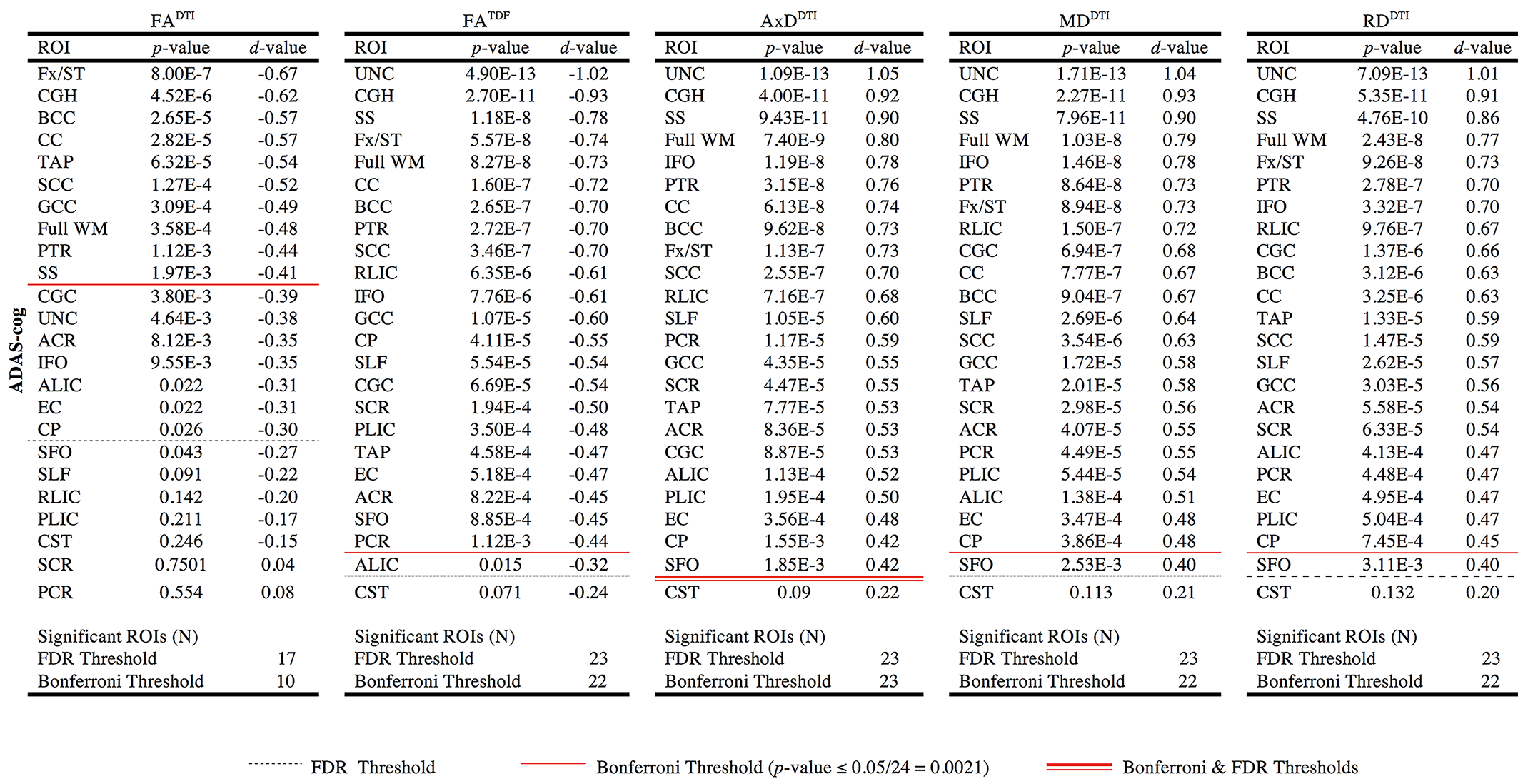

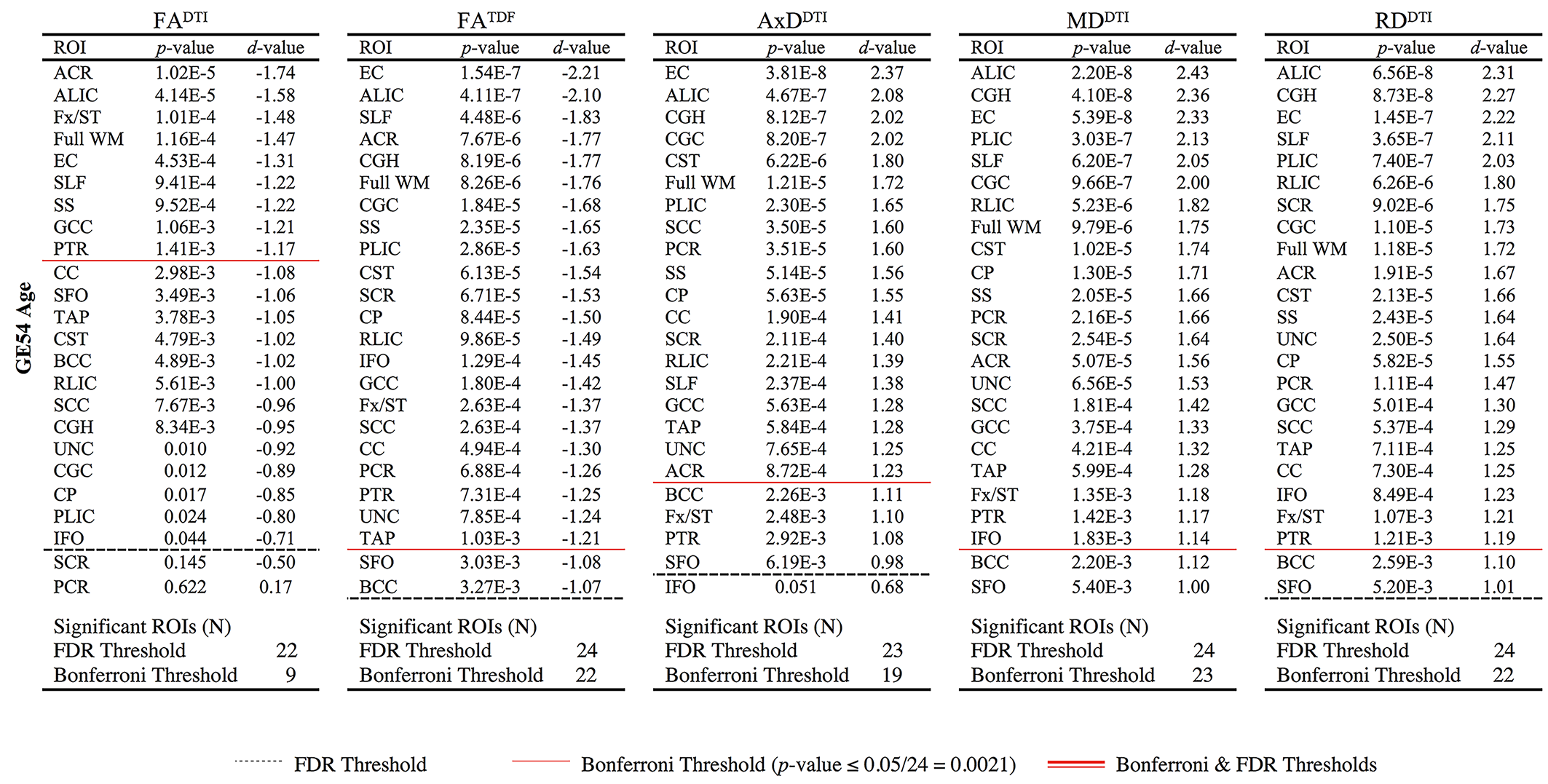

**4.1.3	MMSE Associations**

**Supplementary Table 15.** *P*-values and corresponding effect sizes *(d*-values) are reported for ROI associations between MMSE and dMRI indices across all ADNI3 participants pooled together (N=315). For each test the ROIs are ordered by *d*-value. Regions that were significant after correcting for multiple comparisons (FDR *q*=0.05) are delineated by a dotted line.

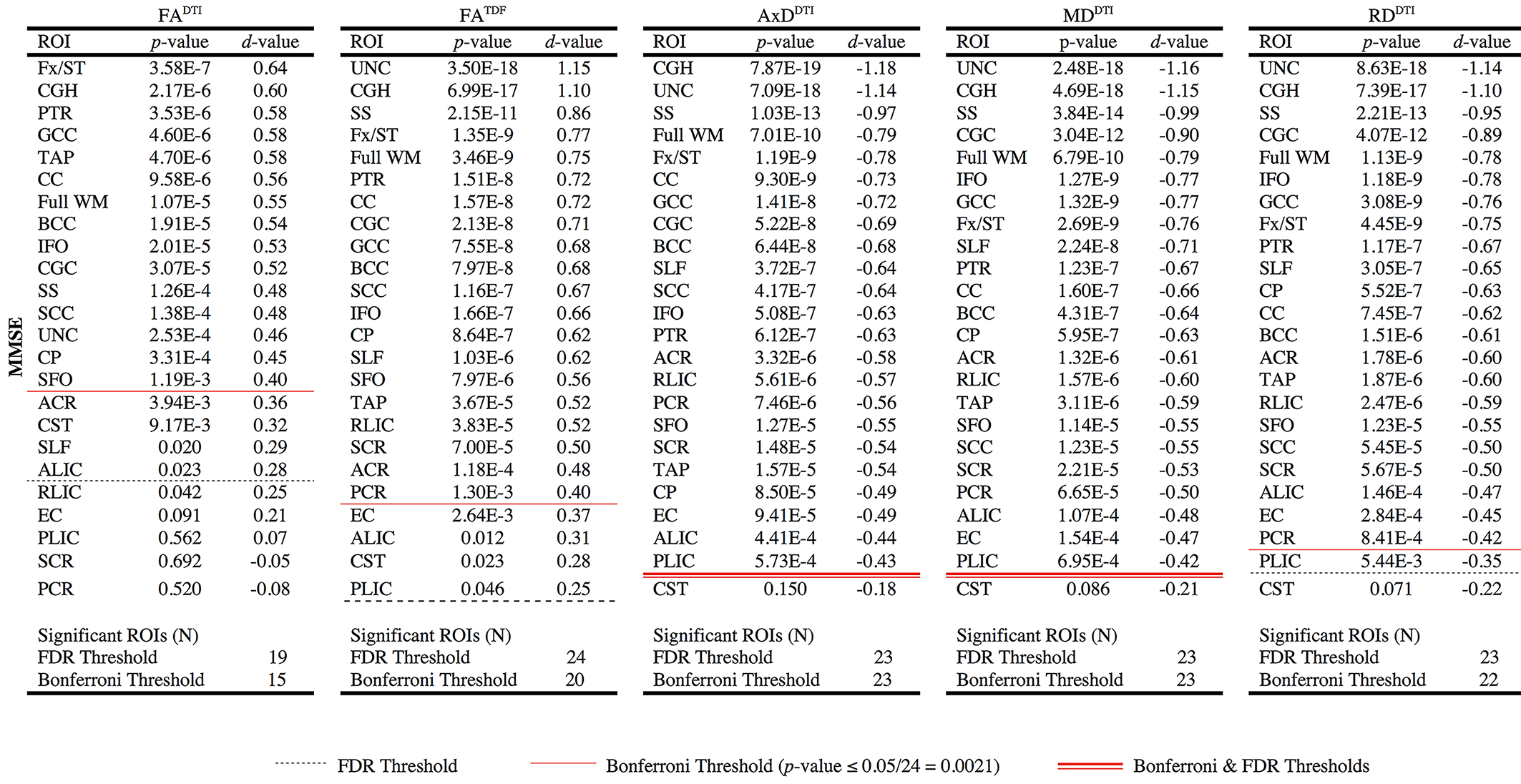

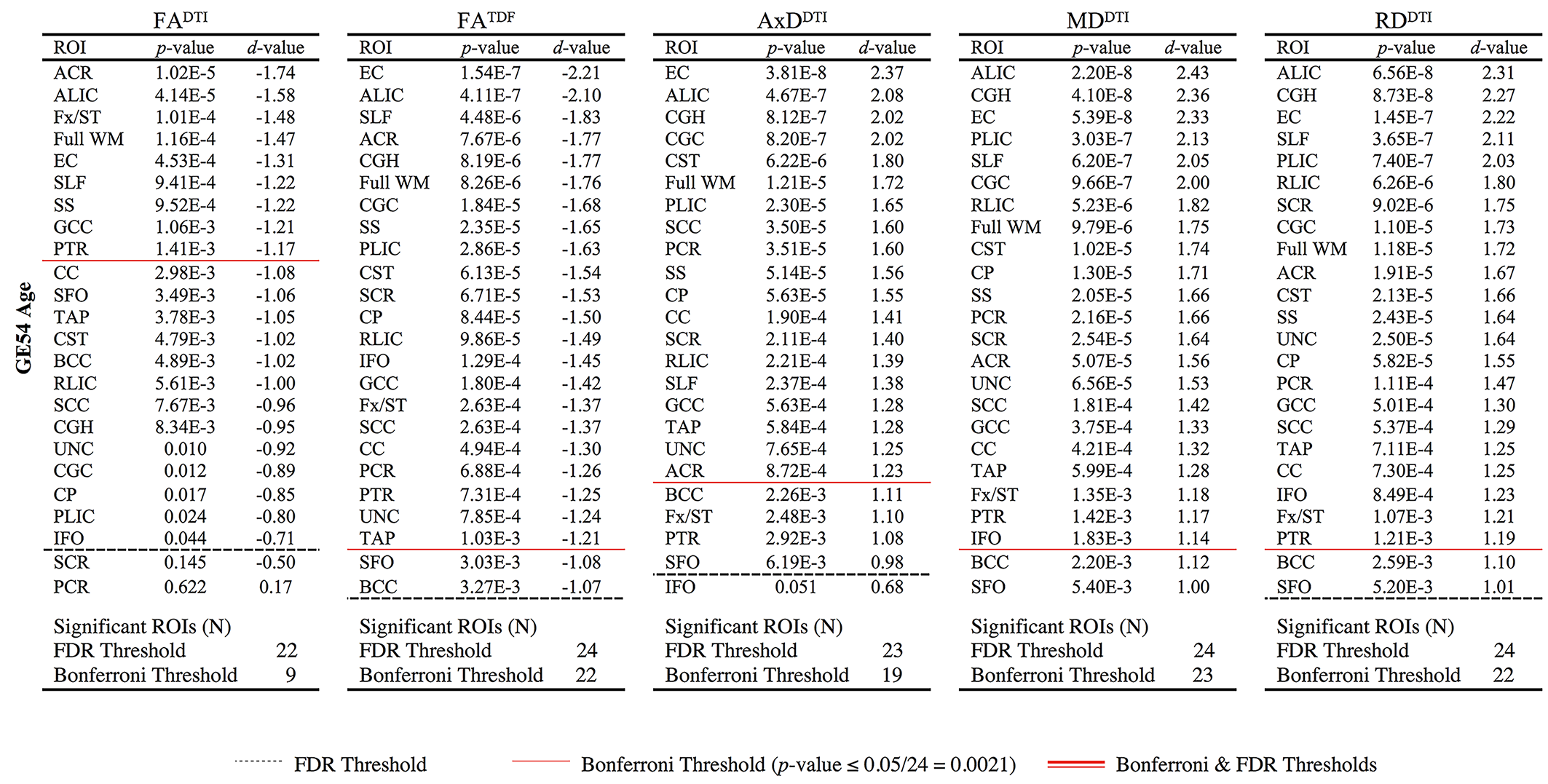

**4.2.	dMRI Associations with Cognitive Measures *by Protocol***

**4.2.1 Full WM Clinical Associations**

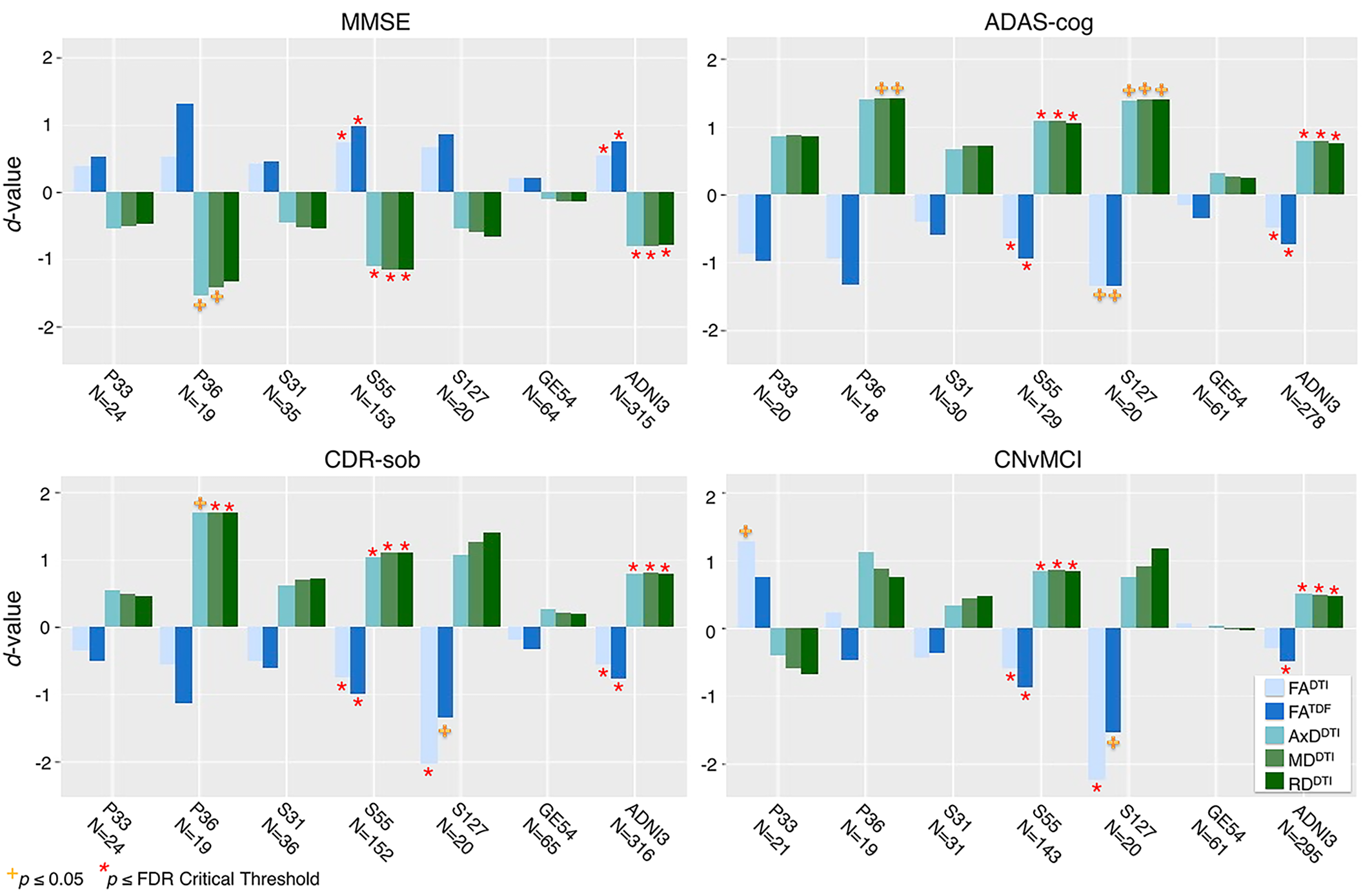

**Supplementary Figure 7.** Effect sizes (*d*-values) from associations between cognitive scores or diagnosis and dMRI indices in the full WM for each of six ADNI3 protocols individually and pooled. We note that due to differences in sample size between protocols, effect sizes should not be directly compared. However, the direction of associations was consistent across protocols.

**4.2.2	CGH Clinical Associations**

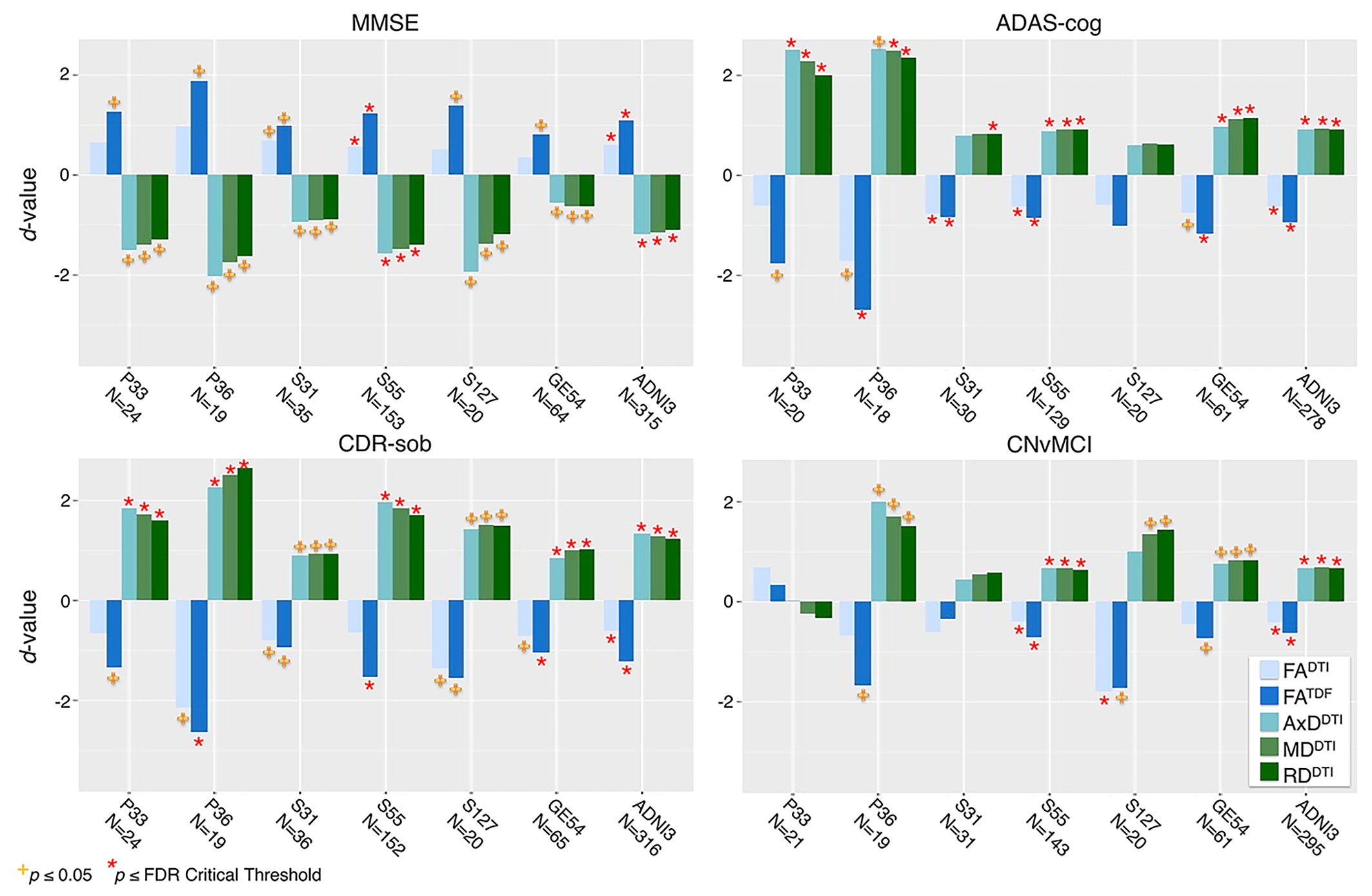

**Supplementary Figure 8.** Effect sizes (*d*-values) from associations between cognitive scores or diagnosis and dMRI indices in the CGH, one of two ROIs that consistently showed significant associations across all four clinical tests and dMRI indices, for each of six ADNI3 protocols individually and pooled. We note that due to differences in sample size between protocols, effect sizes should not be directly compared. However, the direction of associations was consistent across protocols.

**4.2.3	Fx/ST Clinical Associations**

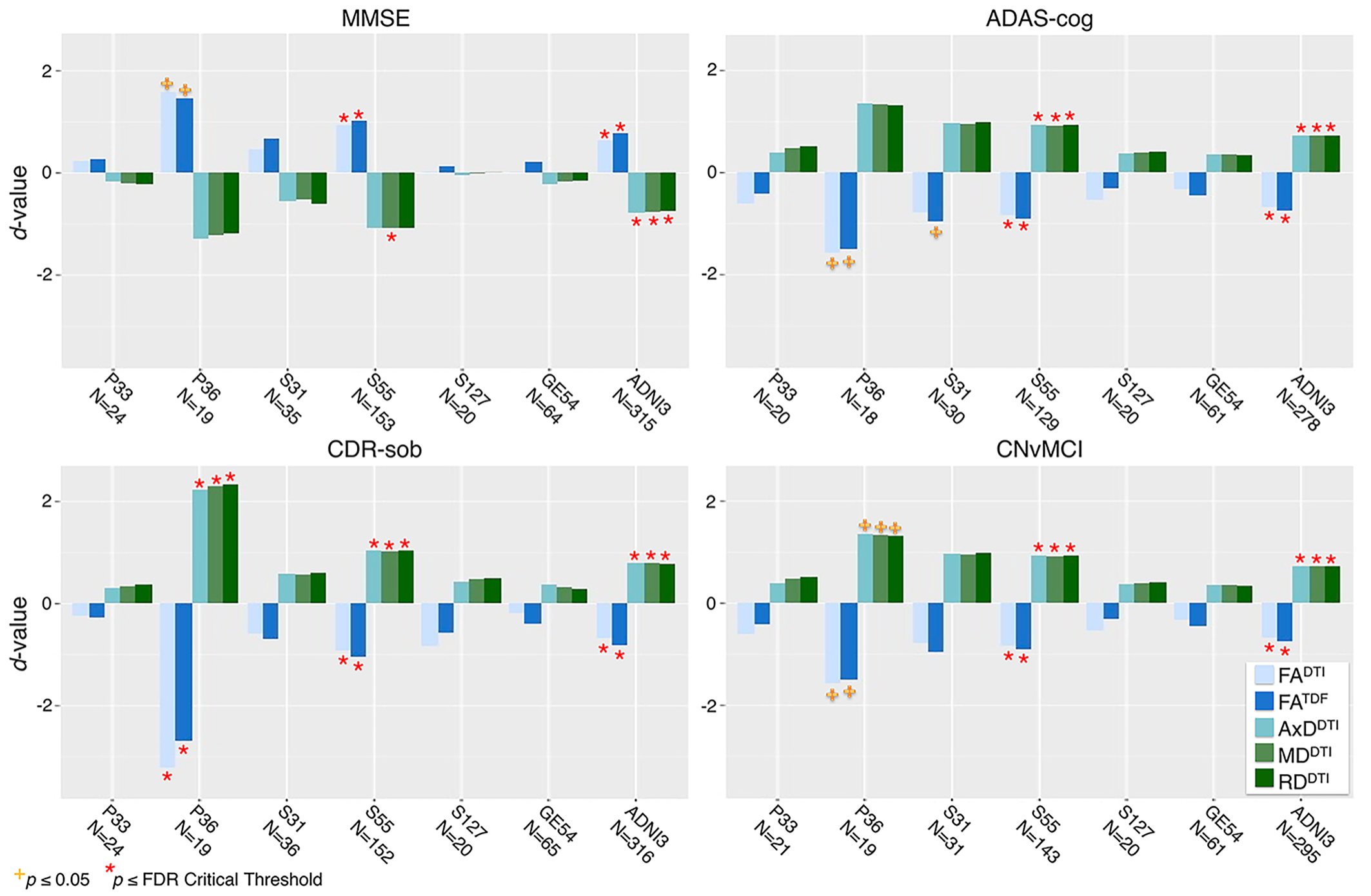

**Supplementary Figure 9.** Effect sizes (*d*-values) from associations between cognitive scores or diagnosis and dMRI indices in the Fx/ST, one of two ROIs that consistently showed significant associations across all four clinical tests and dMRI indices, for each of six ADNI3 protocols individually and pooled. We note that due to differences in sample size between protocols, effect sizes should not be directly compared. However, the direction of associations was consistent across protocols.

**4.2.4	FA^DTI^ vs FA^TDF^ Associations with CDR-sob**

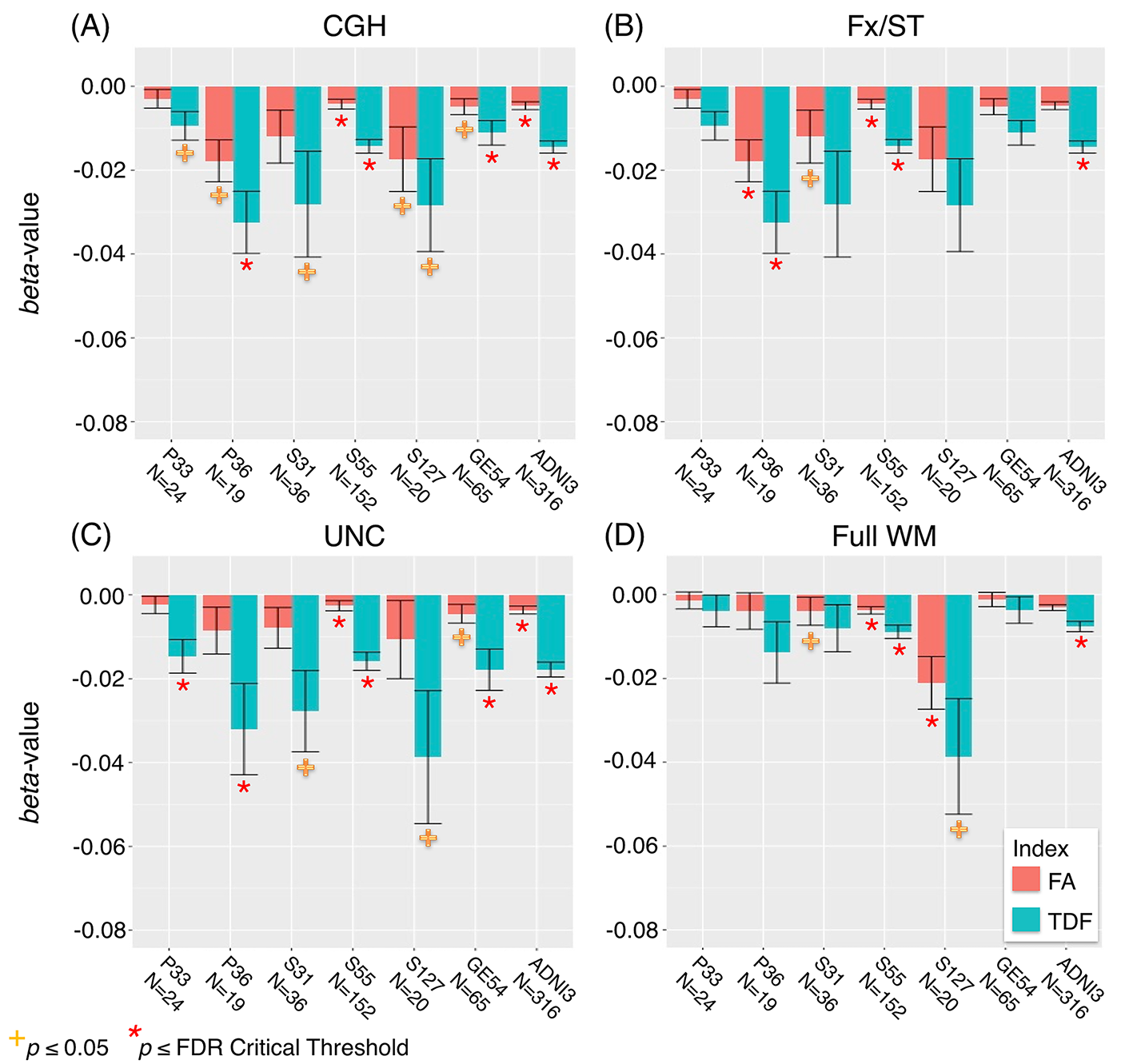

**Supplementary Figure 10.** A comparison of FA^DTI^ and FA^TDF^ effect sizes (*beta*-values and standard error) for CDR-sob associations in the CGH, UNC, Fx/ST, and full WM, for each of six ADNI3 protocols individually and pooled; FA^TDF^ consistently detects larger effect sizes across protocols.

**4.3	Brain Maps of dMRI Associations with Cognitive Measures**

**4.3.1**	**ADAS-cog Associations**

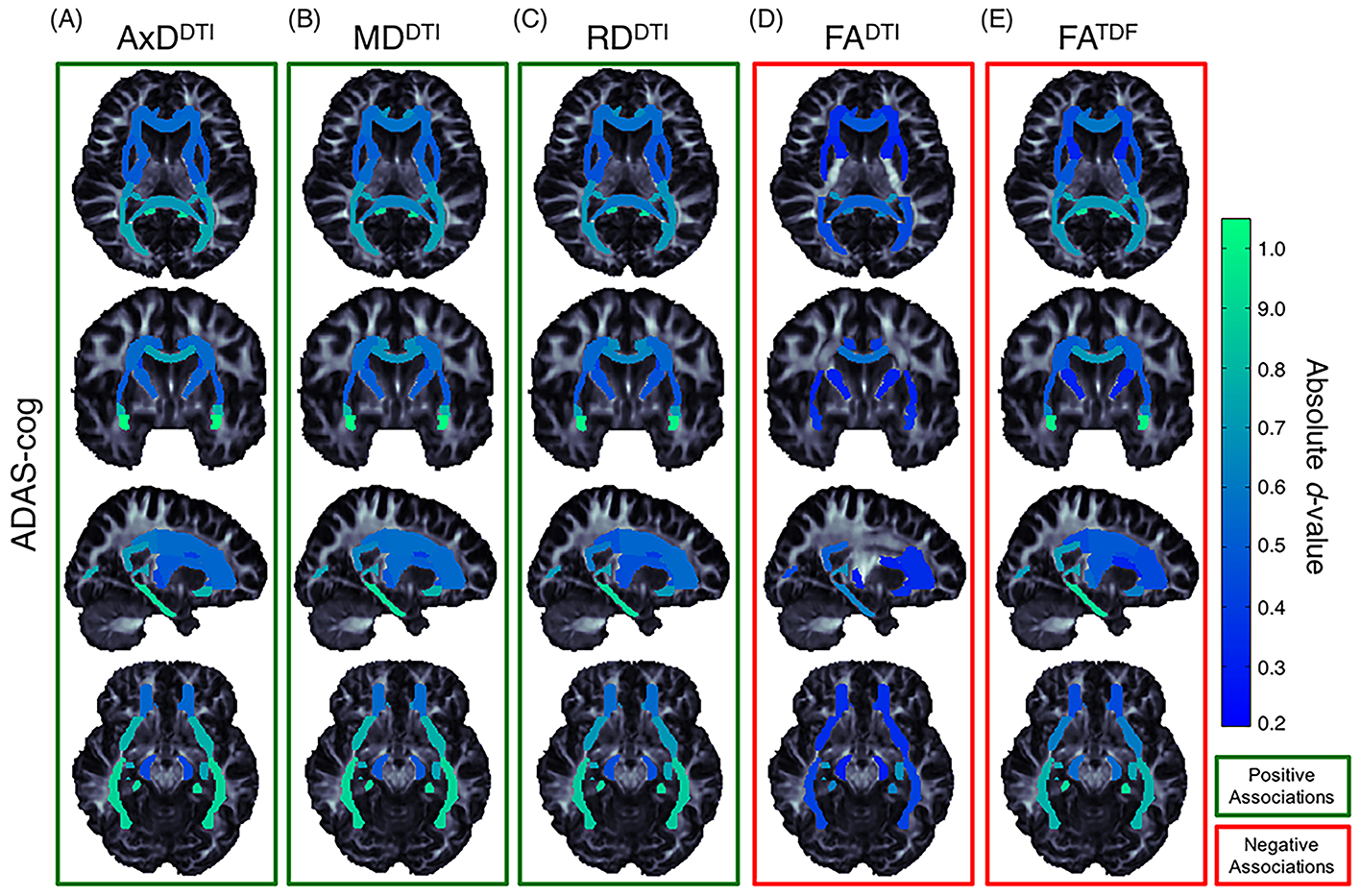

**Supplementary Figure 11.** Effect size (absolute *d*-value) maps of WM regions that show significant associations with ADAS-cog (FDR *q* = 0.05). Associations were detected between ADAS-cog and a) AxD^DTI^ b) MD^DTI^ and c) RD^DTI^, where higher diffusivity was significantly associated with greater cognitive impairment. Lower d) FA^DTI^ and e) FA^TDF^ were associated with greater impairment, but FA^DTI^ associations were detected in fewer regions with weaker effect sizes compared to FA^TDF^. *Light green* regions show the largest effect sizes.

**4.3.2	MMSE Associations**

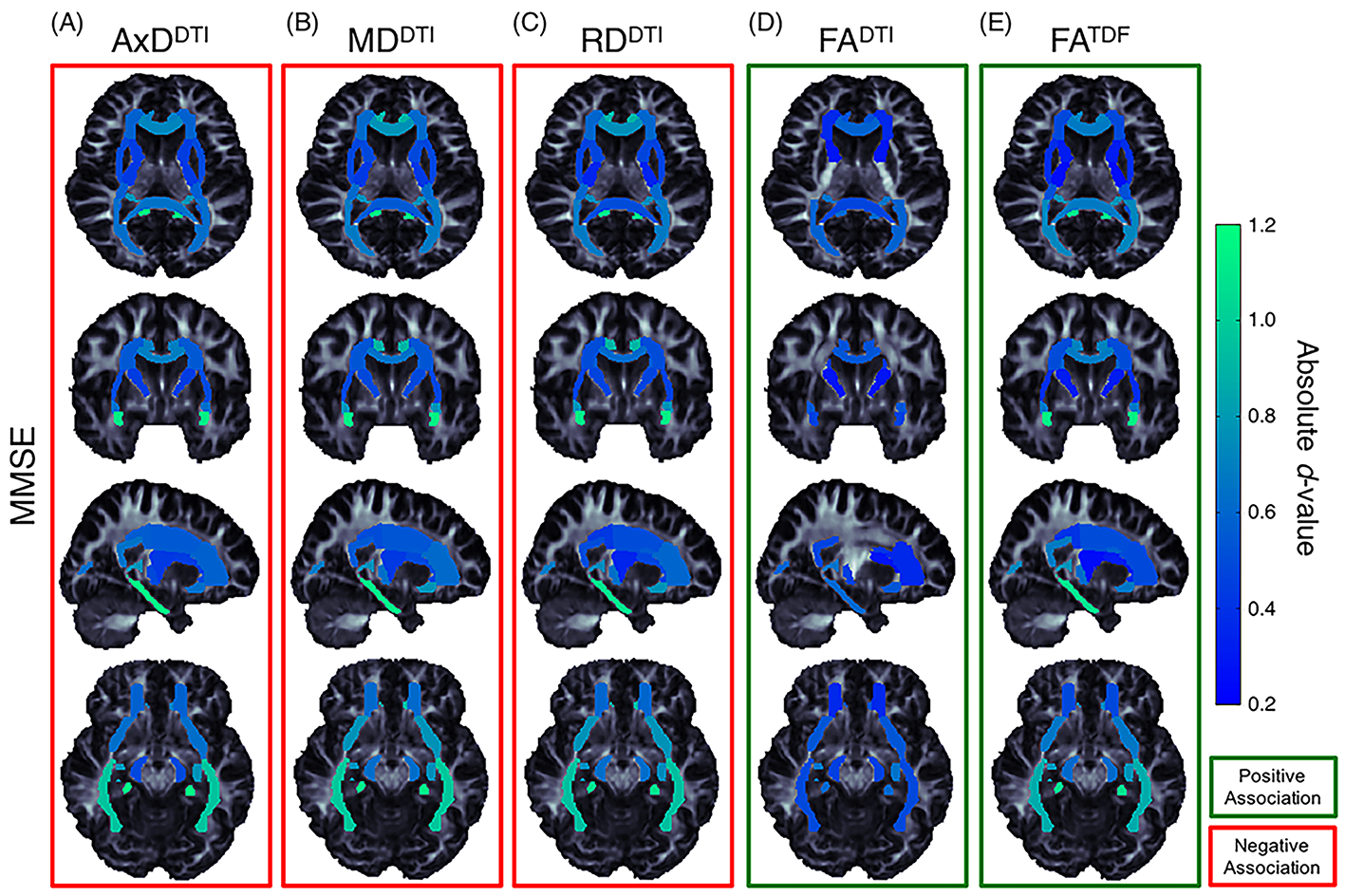

**Supplementary Figure 12.** Effect size (absolute *d*-value) maps of WM regions that show significant associations with MMSE (FDR *q* = 0.05). Negative associations were detected between MMSE scores and a) AxD^DTI^ b) MD^DTI^ and c) RD^DTI^, where higher diffusivity was associated with lower MMSE (greater cognitive impairment). Lower d) FA^DTI^ and e) FA^TDF^ were associated with lower MMSE, but FA^DTI^ associations were detected in fewer regions with weaker effect sizes compared to FA^TDF^. *Light green* regions show the largest effect sizes.

**4.4	ROI Size vs CDR-sob Effect Size**

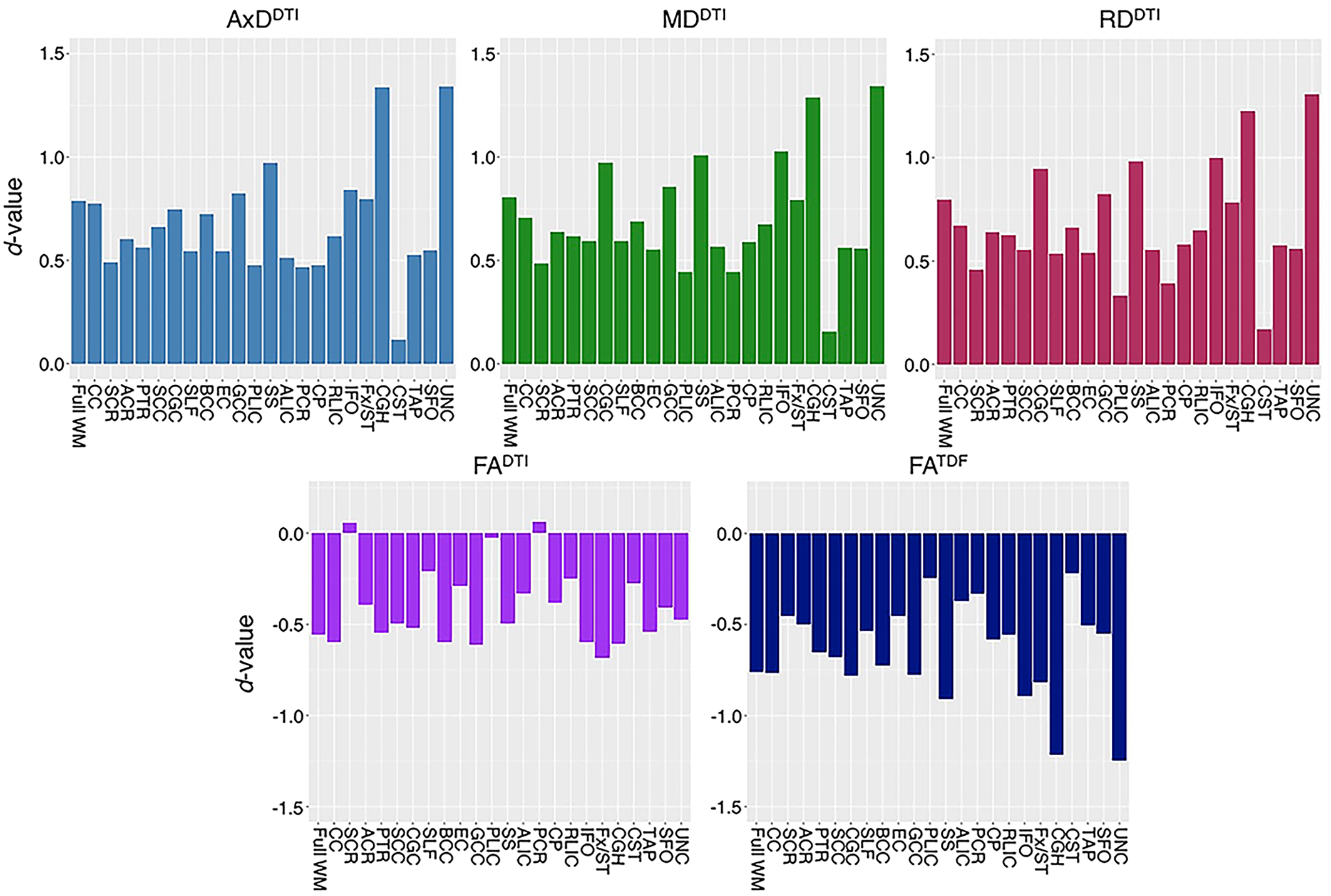

**Supplementary Figure 13.** CDR-sob effect sizes (*d*-values) for each ROI are ordered by ROI size (full WM, far left, is the largest ROI and UNC, far right, is the smallest). Effect size and the square root of ROI size (number of voxels) are not correlated (FA^DTI^: Pearson’s *r* = -1.1, *p* = 0.58; AxD^DTI^: *r* = 0.064 *p* = 0.77; MD^DTI^: *r* = 0.044, *p* = 0.84; RD^DTI^: *r* = -0.19, *p* = 0.93; FA^TDF^: *r* = -0.072, *p* = 0.74).

**4.5	Regional dMRI Measures: Associations with Diagnosis**

**Supplementary Table 16.** *P*-values and corresponding effect sizes *(d*-values) are reported for WM microstructural differences between CN and MCI participants when all ADNI3 dMRI data are pooled together. For each test, the ROIs are ordered by *d*-value. Regions that were significant after FDR (*q* = 0.05) or Bonferroni (α = 0.05) multiple comparisons correction are delineated by a dotted or solid line respectively.

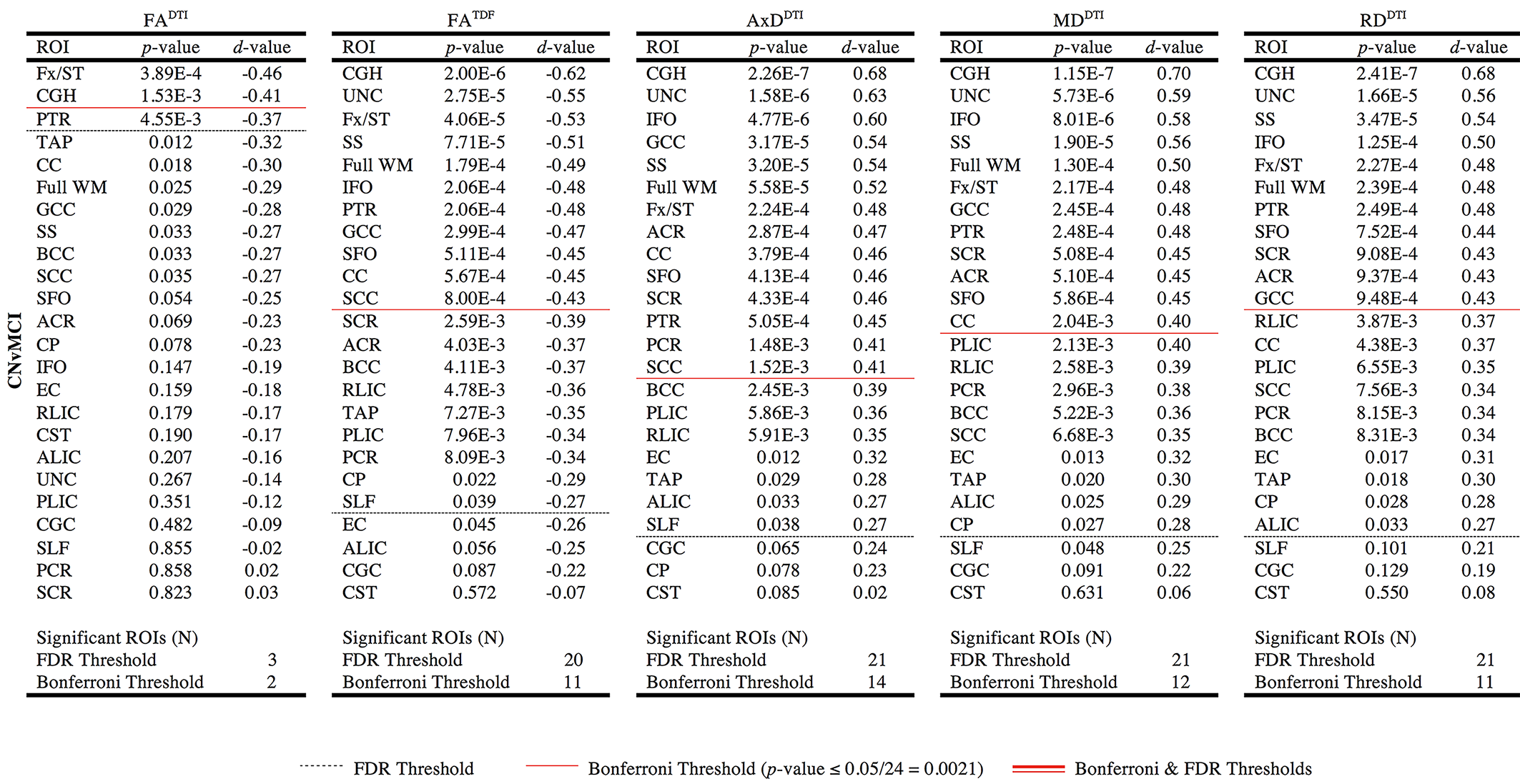

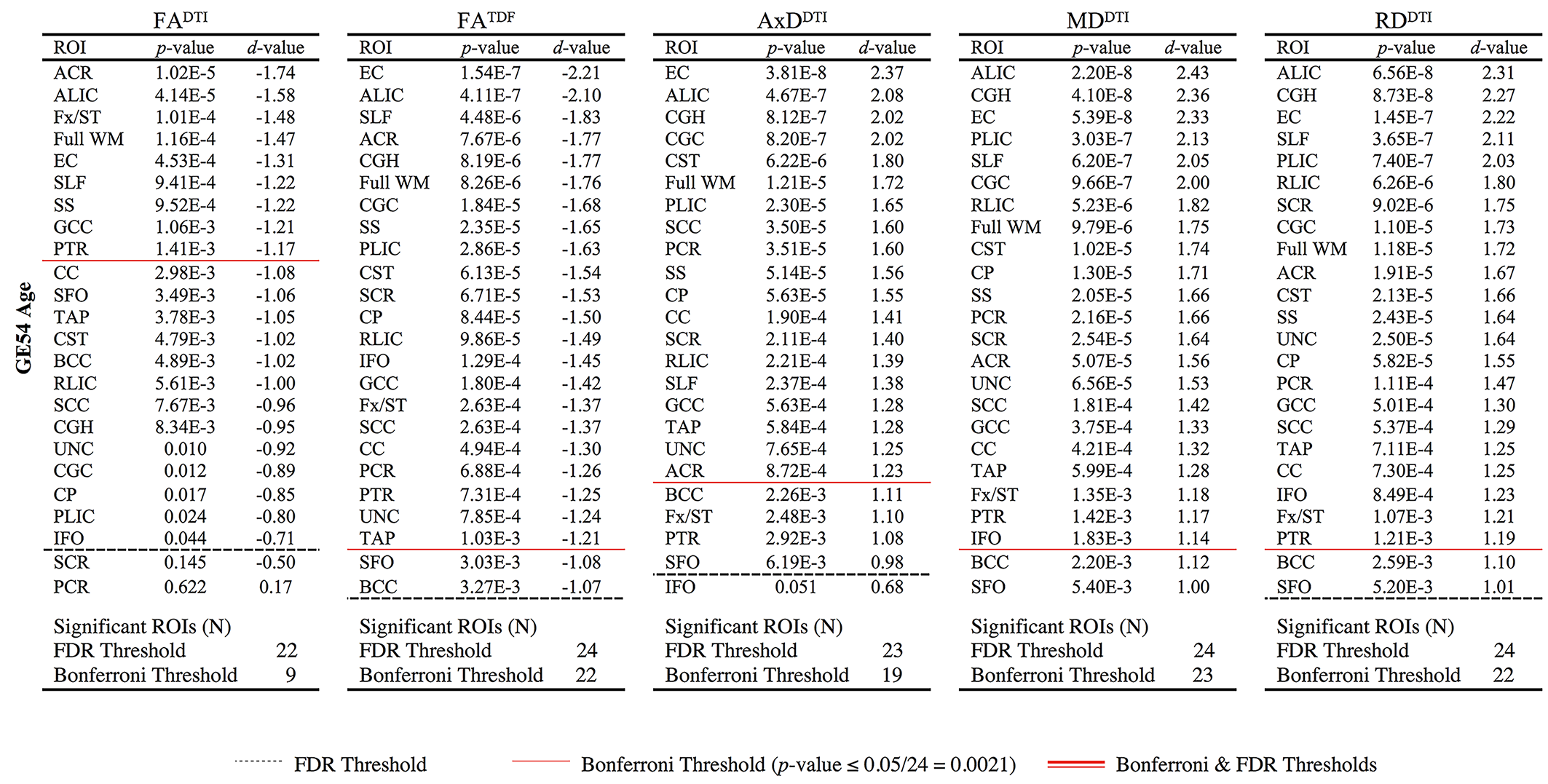

**4.6	Brain Maps of dMRI Associations with Diagnosis**

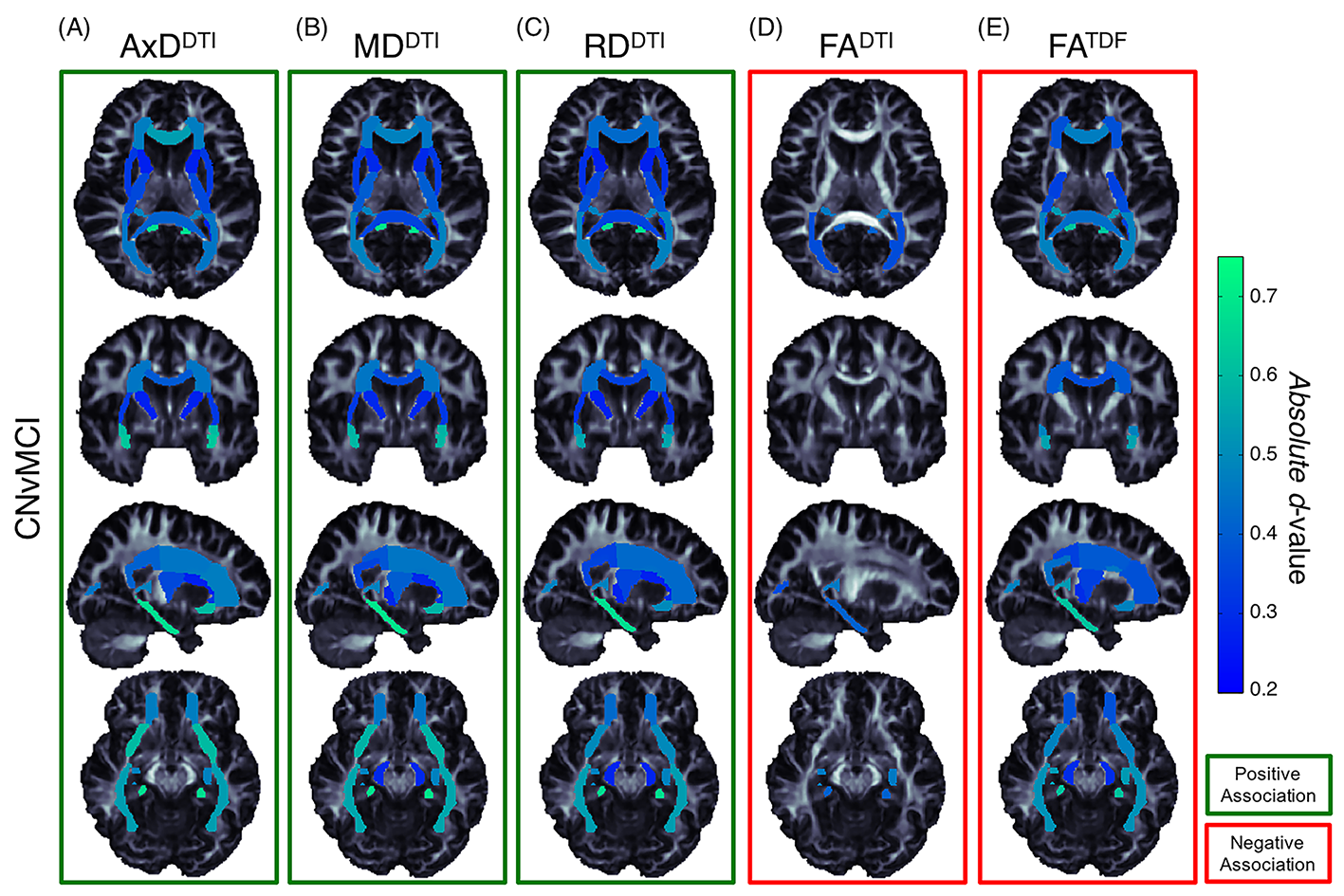

**Supplementary Figure 14.** Effect size (absolute *d*-value) maps of WM regions that show significant differences between CN participants and those with MCI (FDR *q* = 0.05). Positive associations were detected between MCI diagnosis and a) AxD^DTI^ b) MD^DTI^ and c) RD^DTI^, where higher diffusivity was associated with greater cognitive impairment. Lower d) FA^DTI^ and e) FA^TDF^ were associated with greater impairment, but FA^DTI^ associations were detected in fewer regions with weaker effect sizes compared to FA^TDF^. *Light green* regions show the largest effect sizes.
